# In-cell proximity target validation methods for heterobifunctional molecules with CRBN- or VHL-binder using AirID

**DOI:** 10.1101/2025.04.29.651368

**Authors:** Kohdai Yamada, Satoshi Yamanaka, Hiroyuki Yamakoshi, Aki Kohyama, Yoshiharu Iwabuchi, Hidetaka Kosako, Tatsuya Sawasaki

## Abstract

Heterobifunctional molecules, such as proteolysis-targeting and autophagy-targeting chimera, represent new drug concepts. They are composed of two protein binders that can induce proximity interactions between two proteins and protein catalysis. Currently, cereblon (CRBN)- and von Hippel-Lindau (VHL)-binders with thalidomide- and VH032-backbones are widely used as E3 ligase binders. Here we developed a method to validate proteins that interact with heterobifunctional molecules in cells using AirID, a proximity biotinylation enzyme. Interactome of target proteins was validated for six heterobifunctional molecules. ThBD-AirID, a fusion of the thalidomide-binding domain (ThBD) of CRBN and AirID, effectively biotinylated the target proteins. AirID fused to full-length VHL also exhibited highly effective biotinylation. Heterobifunctional molecules with the same target binder but different E3 binders showed different proximity interactome profiles in cells. Analysis using ThBD-AirID revealed a nuclear interaction between androgen receptor and ARV-110. AirID-fused ThBD and VHL could be useful for validating the heterobifunctional molecular interactome in cells.

## Introduction

Targeted protein degradation (TPD) is a therapeutic modality of targeting proteins that are difficult to target using small molecule compounds^1^. Heterobifunctional molecules comprising three components—an E3 ubiquitin ligase-binding binder (E3 binder), a target binder, and a linker connecting the two binders—provide an effective TPD drug design. The main strategy in TPD is to recruit a protein of interest (POI) to E3 ubiquitin ligase or cullin-based complexes, such as the cullin–RING E3 complex, through the E3 and target binders, and then degrade the POI via the ubiquitin-proteasome system. For example, proteolysis-targeting chimeras (PROTACs) can target many proteins previously considered ‘Undruggable Targets’ for conventional small molecule-mediated pharmacology^2,3^. PROTACs have attracted considerable attention in the drug discovery industry and academia. More than 200 PROTACs have been tested for their ability to degrade target proteins^4^, and approximately 6,000 PROTACs are listed in the database with the number expected to increase in the future^5^. ARV-110, an oral PROTAC that targets the androgen receptor (AR), is in clinical trials^6,7^. The PROTAC market is expected to grow further.

A variety of E3 binders that bind to cereblon (CRBN), von Hippel-Lindau (VHL), mouse double minute 2 (MDM2), and other proteins have been used in PROTACs^8,9,10^. However, E3 binders of CRBN and VHL are mainly used for PROTACs because of their efficient POI degradation^11,12^. CRBN and VHL are used as E3 ligases for PROTACs because of the availability of potent E3 binders, such as thalidomide and VH032^13^; however, it is not known whether they have superior capabilities compared to other E3 ligases. Different E3 ligases can be related to different degradation abilities of PROTACs, despite the use of same target binders^14,15,16^. Although the difference in the drug efficacy of these E3 ligases has been attributed to differences in their properties, the underlying mechanism remains unclear. Generally, the main reason for failure in the development of small-molecule compounds is a poor understanding of the POI interactome induced by the molecule, which hampers validation of the target proteins^17,18^. The impact of new interactome formation induced not only by POI binding but also by off-target binding of PROTAC has not been considered^19^. Because an understanding in this regard should promote the development of effective PROTACs, developing PROTAC-induced interactome technologies that enable the analysis of PROTAC targets and off-targets is important.

Proximity biotin labelling allows a comprehensive understanding of protein networks in cells via biotin labelling of proteins in proximity of POI^20,21^. Recently, target identification methods based either on photoaffinity, such as the µMap, or on proximity biotinylation enzymes, such as PROCID, and BioTac have been developed^22,23,24^, significantly expanding the methods for the evaluation of small-molecule compounds. However, the photoaffinity-based methods are not suitable for evaluating PROTACs because they require considerable chemical modifications of polymer-chimeric compounds.

Recently, we developed an original proximity biotinylation enzyme AirID (ancestral BirA for proximity-dependent biotin identification)^25^ and fused it with CRBN to create AirID-CRBN^26^. In addition, we successfully devised a method to analyse the intracellular interactome of thalidomide and its derivatives, known as molecular glue, and ARV-825, a PROTAC^26^. However, AirID-CRBN cannot be used to evaluate PROTACs that use E3 ligases other than CRBN. Furthermore, evaluation of a system that fuses a full-size E3 ligase with a proximity biotinylation enzyme is hampered by competition between ubiquitination and biotinylation because of the same protein surface lysine residues involved. Therefore, further improvements are needed to accurately evaluate TPD. Here, we applied VHL-AirID as an E3 ligase and analysed the PROTAC-dependent interactome. Additionally, for a more accurate evaluation of the target binder in PROTACs, we developed a system that fuses only the E3 binder-binding region, rather than the entire molecule, to AirID. We could analyse the interactome of PROTACs with CRBN binders using this system. Furthermore, AirID, a fusion protein of the E3 binder-binding domain, could be used to identify the proteins interacting with various PROTACs, including those undergoing clinical trials and heterobifunctional molecules. These results establish this approach as a powerful method for identifying targets to better understand the mechanism of action of PROTACs in living cells, as well as for evaluating of different E3 ligases PROTACs-dependent interactome.

## Results

### Comparison of interactomes of BRD binder-dependent PROTACs with different E3 binders

The bromodomain extra-terminal (BET) family proteins, including BRD4 and BRD3, are promising anticancer drug targets^27,28^. ARV-825 and MZ1 were developed to degrade BET proteins in close proximity to E3 ligases, CRBN and VHL, respectively (Fig. 1a)^29,30^. However, MZ1 has not been shown to induce interactome changes in cells, and differences in the interactome induced by MZ1 and ARV-825 remain unknown. We previously reported ARV-825-dependent biotinylation of BET family proteins using stably AirID-CRBN-expressing cells^26^. VHL is a widely used E3 ligase in PROTACs, as is CRBN^1^. However, the PROTAC (comprising CRBN or VHL binder)-dependent interactomes with the same target binder using the same system have not been compared. To compare the two E3 binders under the same conditions, we generated a cell line expressing AirID-fused VHL as described in our previous CRBN analysis. MZ1, which was constructed with a target binder of JQ-1 as in ARV-825, was used as a PROTAC for VHL analysis. First, MM1.S human multiple myeloma cell lines stably expressing AirID-VHL or VHL-AirID were generated to determine the optimal fusion position of AirID in VHL, and MZ1-dependent biotinylation of BRD4 was examined using streptavidin pull-down assay (STA-PDA). MM1.S cells stably expressing AirID-fused VHL were treated with MZ1, and as a negative control with DMSO and VH032, the VHL ligand used in MZ1. Biotinylation of BRD4 was observed only in MZ1-treated cells expressing AirID-VHL and VHL-AirID (Fig. 1b) but not in the controls. Notably, BRD4 biotinylation by VHL-AirID was higher than that by AirID-VHL, indicating that VHL-AirID has a higher ability to biotinylate target proteins. Therefore, we used VHL-AirID for further analysis.

**Fig. 1.**
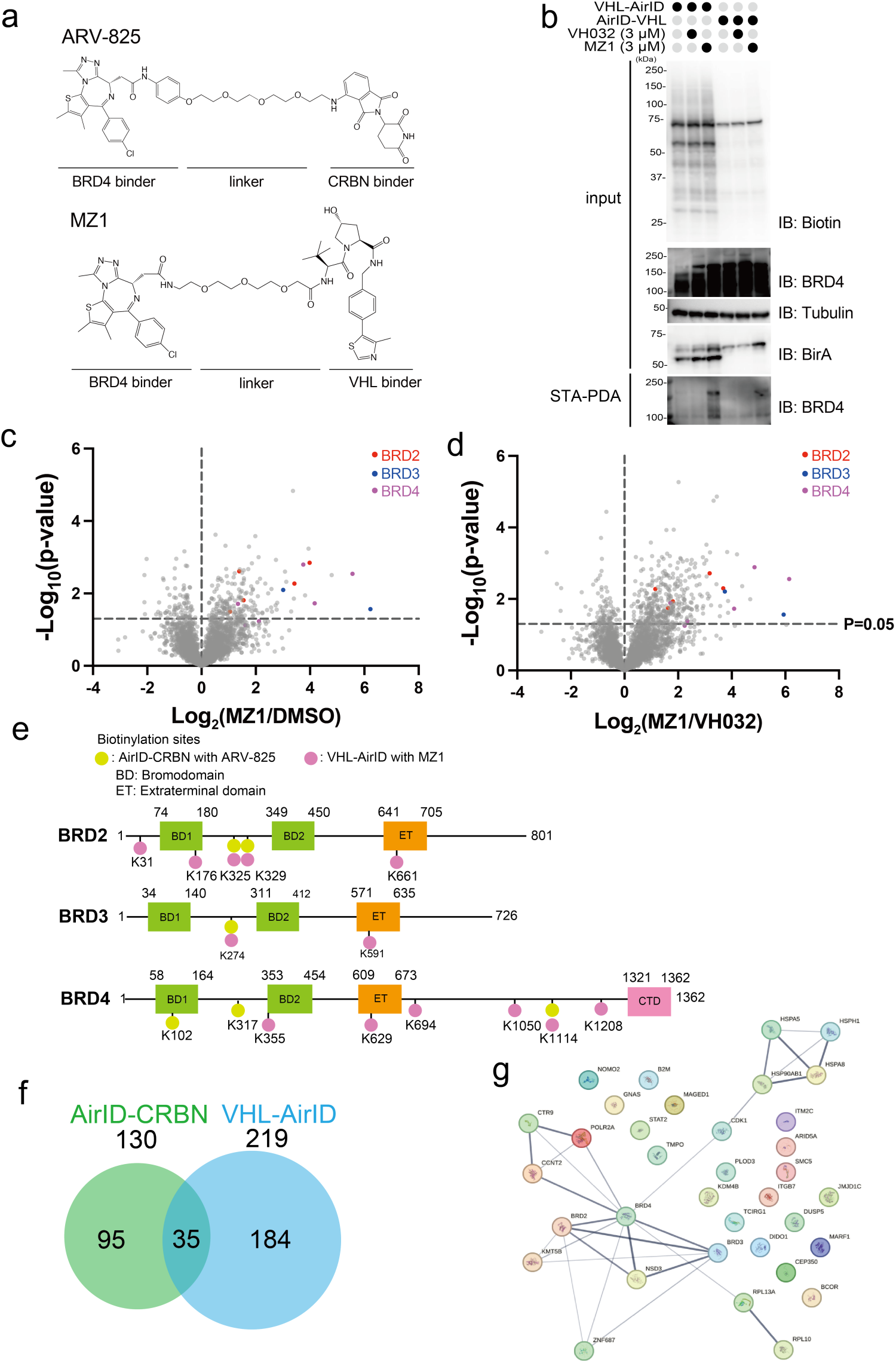
MZ1-dependent biotinylation by VHL-AirID compared with ARV-825-dependent biotinylation by AirID-CRBN. **a** Chemical structures of ARV-825 and MZ1. **b** Streptavidin pull-down assay of MZ1-dependent biotinylation in VHL-AirID- and AirID-VHL-expressing MM1.S cells. Three micromolar MZ1 or VH032 together with 5 µM MG132 and 10 µM d-biotin was added to the cells and incubated for 6 h for biotinylation reaction. **c, d** Volcano plots of biotinylated peptides detected via LC-MS/MS in MM1.S cells stably expressing VHL-AirID. Significant changes in the volcano plots were calculated using Student’s two-sided *t*-test, and the false discovery rate (FDR)-adjusted *P*-values calculated using the Benjamini–Hochberg method are shown in Supplementary Data 1. c Volcano plot comparing MZ1 treatment with DMSO treatment; d MZ1 treatment with VH032 treatment. **e** Diagram of ARV-825- or MZ1-dependent biotinylation sites using AirID-CRBN or VHL-AirID for BET family proteins (BRD2, BRD3, and BRD3). The yellow-green colour represents the ARV-825-dependent biotinylation site for AirID-CRBN, and the pink colour represents the MZ1-dependent biotinylation site for VHL-AirID. **f** Venn diagram of ARV-825- or MZ1-dependent proteins biotinylated by AirID-CRBN or VHL-AirID (PROTAC/E3 ligand ratios of biotinylated peptides detected via LC-MS/MS >2 and *P* < 0.05). **g** Interaction analysis of 35 proteins obtained from the Venn diagram shown in Fig. 1f.

The proteins biotinylated by the treatment of VHL-AirID-expressing MM1.S cells with MZ1 were identified using liquid chromatography-tandem mass spectrometry (LC-MS/MS) (Fig. 1c, d, Supplementary Data 1). The results showed that MZ1 targeted BRD proteins were enriched to higher levels in the MZ1-treated compartment than in the DMSO-treated compartment (Fig. 1c). In addition, BRDs were biotinylated by treatment of cells with MZ1 but not with the VHL ligand VH032 (E3 binder for MZ1) (Fig. 1d). Furthermore, we identified biotinylation sites in biotinylated proteins using our previously reported method employing Tamavidin-2 REV for enriching biotinylated peptides^31^. Biotinylation sites for AirID-CRBN (yellow-green spots in Fig. 1e) and VHL-AirID (pink spots) were spotted on BRD2, BRD3, and BRD4. Notably, four biotinylation sites (yellow and pink overlapping spots) were common among the three BRD proteins, and the number of sites for VHL-AirID was higher than that for AirID-CRBN (Fig. 1e). These results indicate that different chemical compositions of E3 binders and E3-binding proteins in PROTACs can lead to different modes of interaction with the target protein, even with the same target binders.

These results were compared with our previous report on biotinylated proteins when AirID-CRBN-expressing MM1.S cells was supplemented with ARV-825 (Fig. 1f)^26^. In total, 219 and 139 proteins were biotinylated by MZ1 and ARV-825, respectively (Supplementary Fig. 1a, b). Analysis of the protein–protein interactome of MZ1-dependent proteins biotinylated by VHL-AirID using the STRING database showed that 33 BRD4-interacting proteins were biotinylated (Supplementary Fig. 2a)^32^. In contrast, 26 BRD4-interacting proteins were biotinylated by AirID-CRBN in an ARV-825-dependent manner (Supplementary Fig. 2b). Thirty-five common proteins were biotinylated by MZ1 and in an ARV-825-dependent manner (Supplementary Fig. 2c). Gene ontology (GO) analysis using ShinyGO of these 35 proteins showed enrichment of GO terms for histone methylation. GO analysis revealed that proteins closely related to the BET family were biotinylated (Supplementary Fig. 2d)^33^. Interactome analysis of the 35 proteins revealed that eight of them interact with the BET proteins (Fig. 1g). Taken together, these results show that even for VHL, an E3 ligase different from CRBN, fusion with AirID can lead to target protein biotinylation in a PROTAC-dependent manner, indicating that the AirID-fusion system can be used for other E3 ligases in PROTAC validation.

### VHL ligand-binding domain does not function in cells

Full-length E3 ligases fused to AirID can be used to analyse the PROTAC-induced interactome. In addition to the ligand-binding domain, full-length E3 ligases have multiple domains that form complexes with other proteins^10^. For AirID fusion with full-length E3 ligase, proteins that bind to regions other than the ligand-binding domain are also biotinylated; thus, fusing the minimal ligand-binding domain to AirID should provide a more accurate analysis of the interactome of heterobifunctional molecules. Biotinylation and ubiquitination likely use the same surface lysine residues on the proximal protein. Because full-length E3 ligases are capable of ubiquitination, biotinylation and ubiquitination can compete within the same complex if full-length E3 ligases are used. Therefore, the minimal ligand-binding domain of the E3 ligase was fused to AirID. First, because the 54–155 aa region of VHL is the VHL ligand-binding domain (VBD)^34,35^, it was fused to AirID with an AGIA tag (Fig. 2a). Because the AGIA tag has no lysine residues, it was used to detect proximity biotinylation^36^. As the fusion site of AirID to VBD could influence the interaction between VBD and the VHL ligand, AirID was fused at the N- and C-terminus, respectively, and MZ1-dependent biotinylation of BRD4 was compared. Two types of AirID-fusion VBDs, N-terminal (AGIA-AirID-VBD) and C-terminal (VBD-AGIA-AirID), and full-length VHL-AirID were synthesised in a wheat cell-free protein production system and compared for biochemical MZ1-dependent biotinylation using the AlphaScreen system (Fig. 2b). In vitro biotinylation was performed by mixing the AirID-fusion protein with FLAG-GST-BRD4 (FG-BRD4) and incubating for 3 h. The biotinylation ability of the AirID-fused VBD or full-length VHL was determined by measuring the biotinylation rate of FG-BRD4 using the AlphaScreen system. Results of biochemical analysis showed that FG-BRD4 was biotinylated in a concentration-dependent manner by MZ1 in all AirID-fusion proteins (Fig. 2c), but not by the VHL binder alone, VH032. MZ1-dependent biotinylation of FG-BRD4 by AirID-fused VBD or VHL protein was also confirmed in a STA-PDA using streptavidin-conjugated beads (Fig. 2d). These results indicated that AirID-fused VBD can be used for MZ1-dependent BRD4 biotinylation in vitro.

**Fig. 2.**
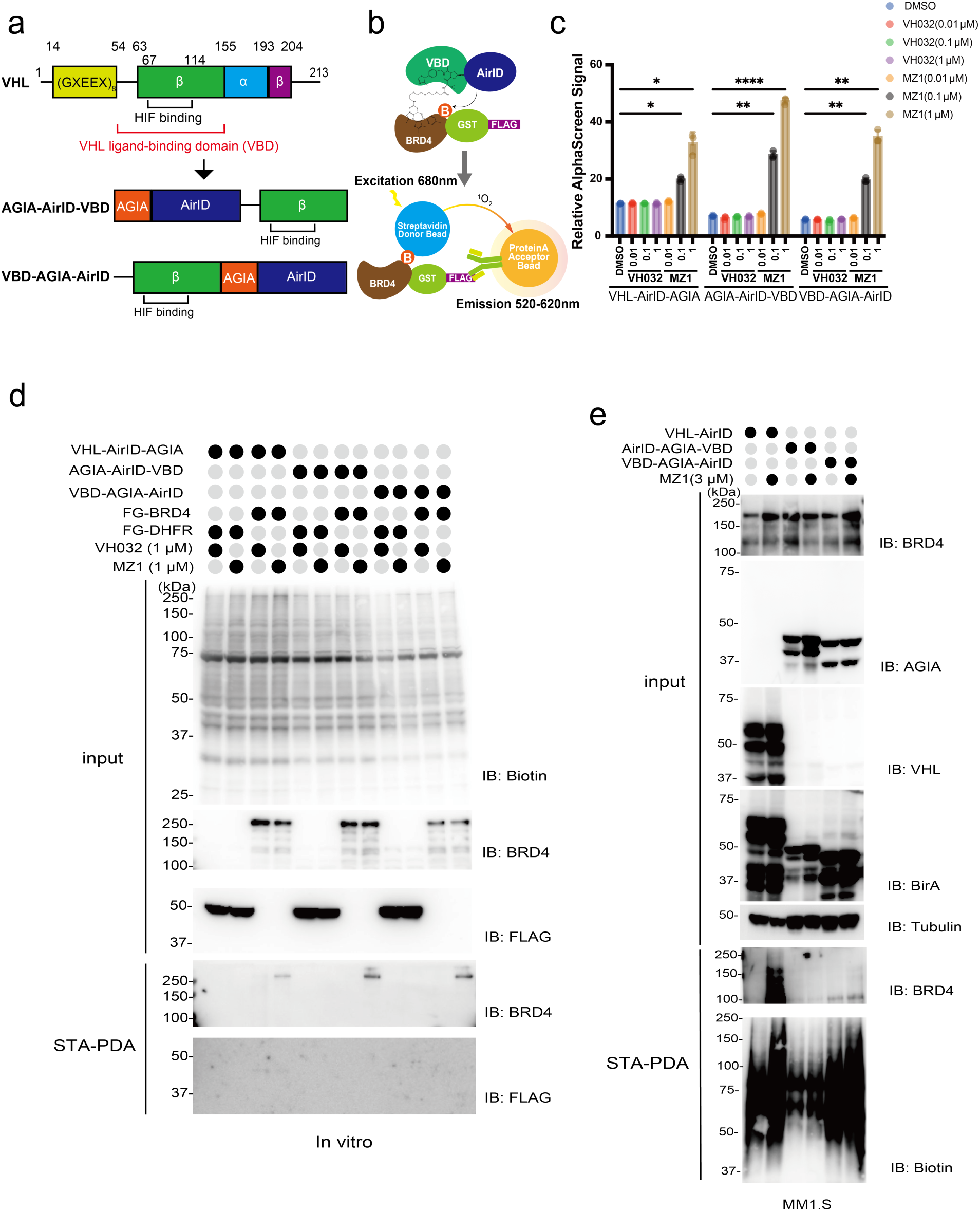
MZ1-dependent proximity biotinylation ability of AirID-fused VBD. **a** Schematic of VBD fused to AirID. AirID was fused to N- or C-terminal VBD. **b** Schematic of the detection of BRD4 biotinylated by AirID-fused VBD with MZ1 using the AlphaScreen assay. **c** AlphaScreen assay for detecting biotinylated BRD4 using AirID. All relative AlphaScreen signals represent the relative values of the AirID-fused VHL and FLAG-GST-DHFR for each drug treatment. Error bars denote standard deviations (independent experiments, *n* = 3). Statistical significance was determined using a two-way ANOVA with Tukey’s multiple comparison test (\*\*\*\**P* < 0.0001, \*\*\**P* < 0.0002, \*\**P* < 0.0021, \**P* < 0.0332). **d** Streptavidin pull-down assay of MZ1-dependent BRD4 biotinylation by VHL-AirID-AGIA, AGIA-AirID-VBD, and VBD-AGIA-AirID in vitro. **e** Streptavidin pulldown assay of MZ1-dependent proteins biotinylated in MM1.S cells stably expressing VHL-AirID, AGIA-AirID-VBD, or VBD-AGIA-AirID. Three micromolar MZ1 together with 5 µM MG132 and 10 µM d-biotin was added to the cells and incubated for 6 h for biotinylation reaction.

Next, MZ1-dependent biotinylation of BRD4 was examined using cells expressing AirID-AGIA-VBD or VBD-AGIA-AirID. First, we generated MM1.S cell lines stably expressing AirID-AGIA-VBD or VBD-AGIA-AirID and compared their MZ1-dependent biotinylation of BRD4 (Fig. 2e). MM1.S cells stably expressing the AirID-fused VBD were treated with MZ1 plus biotin; VH032 was used as a negative control. After biotinylation and cell extraction, STA-PDA was performed and BRD4 biotinylation in the respective cell lines was detected using immunoblotting. MZ1-dependent biotinylation of BRD4 was observed only for AirID-fused full-length VHL (IB: BRD4 of STA-PDA panel in Fig. 2e) and not for both AirID-fused VBDs, VBD-AGIA-AirID and AirID-AGIA-VBD. In cells stably expressing AirID-fusion proteins, only full-length VHL had the ability to biotinylate BRD4 in an MZ1-dependent manner, whereas both AirID-fused VBDs had biotinylation capacity in vitro but not in cells.

### Validation of IMiD-dependent biotinylation of AirID-fused thalidomide binding domain

Because the biochemically functional AirID-fused VBDs could not biotinylate BET family proteins in cells, we next investigated whether the thalidomide-binding domain of CRBN could be used. We previously performed structural analysis of the thalidomide-binding domain^37^. According to the report, the 318–424 aa region of CRBN was selected, and the C366S mutant was used as the thalidomide-binding domain (ThBD) because the C366S mutation stabilises ThBD. AirID fused to the N- or C-terminus of ThBD, AirID-AGIA-ThBD or ThBD-AGIA-AirID respectively, were constructed (Fig. 3a). To compare their biochemical biotinylation abilities, the two AirID fusion ThBDs and an AirID-fused full-length CRBN (AirID-CRBN) were produced using a wheat cell-free protein production system. SALL4 was used as a substrate for the FLAG-GST fusion (FG-SALL4) because it interacts with CRBN in an IMiD-dependent manner^37,38^. Pomalidomide-dependent biotinylation of FG-SALL4 by AirID-fusion proteins was detected using the AlphaScreen method (Fig. 3b). For biochemical assay, each AirID-fusion protein was mixed with FG-SALL4 and biotin, and then incubated for 3 h. Pomalidomide-dependent biotinylation of FG-SALL4 was observed for AirID-CRBN and ThBD-AGIA-AirID (Fig. 3c), but not for AirID-AGIA-ThBD. Biotinylation of FG-SALL4 by ThBD-AGIA-AirID showed a higher signal than that by AirID-CRBN in the presence of 10 µM pomalidomide. Pomalidomide-dependent FG-SALL4 biotinylation by ThBD-AGIA-AirID was confirmed using immunoblot analysis (Fig. 3d). Biotinylation of FG-SALL4 with pomalidomide by ThBD-AGIA-AirID produced the clearest band among the four AirID-fusion proteins (right lane in Fig. 3d). Next, biotinylation of SALL4 by ThBD-AGIA-AirID using other IMiDs was performed. The AlphaScreen assay showed that thalidomide (Tha)- and lenalidomide (Len)-dependent biotinylation of FG-SALL4 was induced by ThBD-AGIA-AirID (Fig. 3e), indicating that the IMiD-dependent interaction-inducing capability of the C-terminal AirID-fused ThBD is almost equivalent to that of the full-length CRBN.

**Fig. 3.**
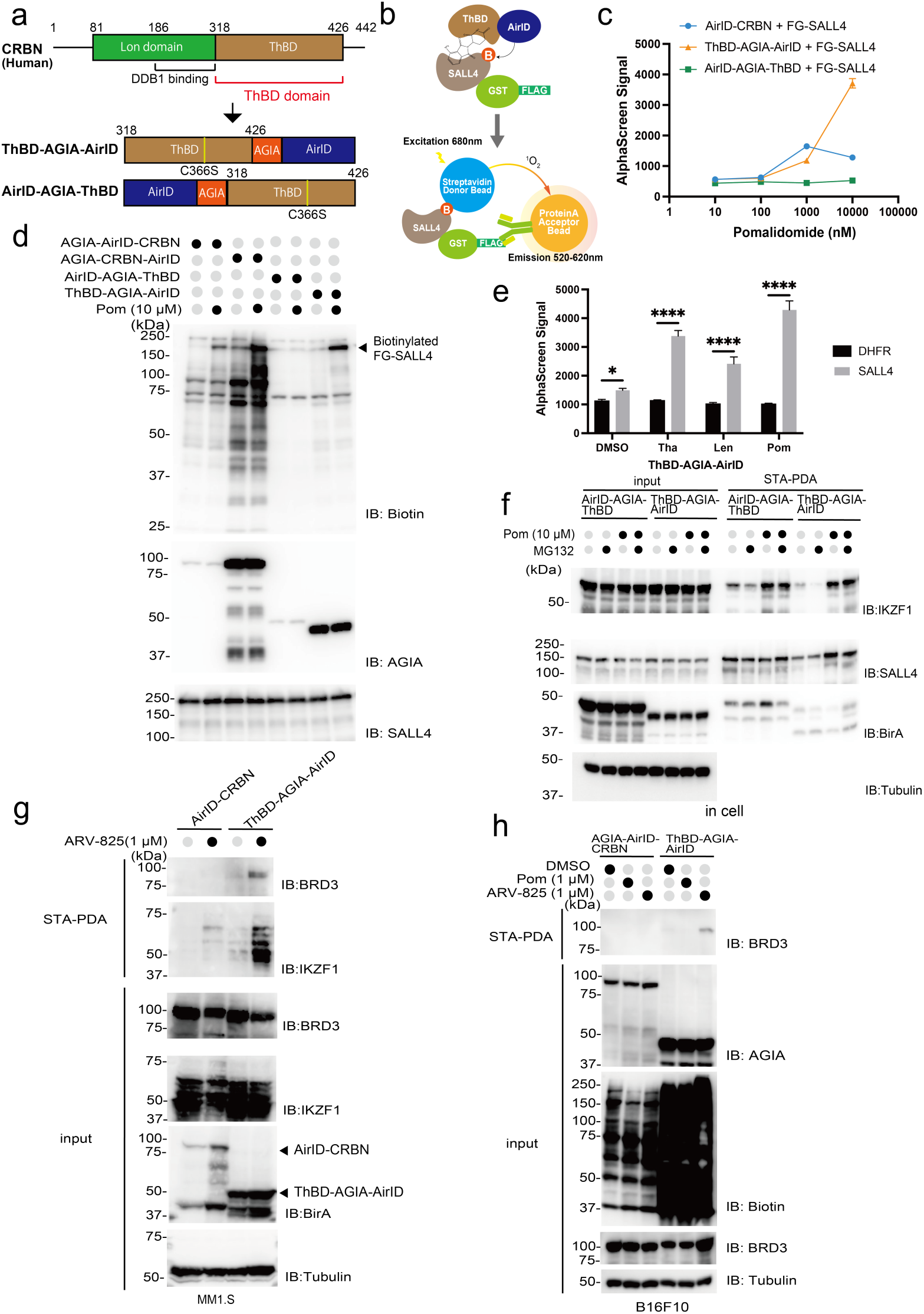
Proximity biotinylation ability of AirID-fused ThBD with molecular glue and PROTAC. **a** Schematic of ThBD fused to AirID. AirID was fused to N- or C-terminal ThBD. **b** Schematic of the detection of SALL4 biotinylated by AirID-fused ThBD with pomalidomide using the AlphaScreen assay. **c** AlphaScreen assay detection of SALL4 biotinylated by AirID-fused CRBN or ThBD. Error bars denote standard deviations (independent experiments, *n* = 3). **d** Immunoblot assay of pomalidomide-dependent SALL4 biotinylation by AGIA-AirID-CRBN, AGIA-CRBN-AirID, AirID-AGIA-ThBD, and ThBD-AGIA-AirID in vitro. **e** In vitro detection of SALL4 biotinylation by thalidomide, lenalidomide, and pomalidomide-dependent ThBD-AGIA-AirID using the AlphaScreen assay. IMiDs were subjected to the biotinylation reaction at a concentration of 20 µM. Error bars denote standard deviations (independent experiments, *n* = 3). Statistical significance was determined using two-way ANOVA with Bonferroni’s multiple comparison test (\*\*\*\**P* < 0.0001, \**P* < 0.0332). **f** Streptavidin pull-down assay of pomalidomide-dependent IKZF1 and SALL4 biotinylated in HEK293T cells stably expressing AirID-AGIA-ThBD or ThBD-AGIA-AirID. **g** Streptavidin pull-down assay of ARV-825-dependent biotinylation of IKZF1 and BRD3 in MM1.S cells stably expressing AGIA-AirID-CRBN or ThBD-AGIA-AirID. **h** Streptavidin pull-down assay of ARV-825-dependent biotinylation of BRD3 in B16F10 cells stably expressing AGIA-AirID-CRBN and ThBD-AGIA-AirID.

### IMiD- and PROTAC-dependent biotinylation of target proteins by ThBD-AGIA-AirID in cells

Next, we examined the ability of ThBD-AGIA-AirID to induce pomalidomide-dependent biotinylation of neosubstrates. HEK293T cells were constructed with steady AirID-AGIA-ThBD and ThBD-AGIA-AirID expression levels. IKZF1 and SALL4 were overexpressed as neosubstrates in these cells and treated with pomalidomide and biotin^38,39^. After STA-PDA of cell lysates, the samples were subjected to immunoblotting. The results showed that biotinylation of IKZF1 was induced by both AirID-AGIA-ThBD and ThBD-AGIA-AirID (Fig. 3f), and pomalidomide-dependent biotinylation of SALL4 was observed only for ThBD-AGIA-AirID. These results were consistent with those of in vitro biotinylation. Furthermore, as ThBD-AGIA-AirID lacks the DDB1-binding domain of CRBN, biotinylation of SALL4 was detected even in the absence of treatment with the proteasome inhibitor MG132. These results indicate that ThBD-AGIA-AirID can be used to analyse the molecular glue-dependent biotinylation of neosubstrates in cells without the addition of proteasome inhibitors. However, the conditions under which ThBD-AGIA-AirID can be used without proteasome inhibitors, because of the endogenous expression of CRBN in cells, are currently limited to those where the neosubstrate is overexpressed or where endogenous CRBN is knocked out.

To investigate whether ThBD-AGIA-AirID has PROTAC-dependent biotinylation activity in cells, biotinylation analysis of the BRD protein using ARV-825 was performed. MM1.S cells stably expressing AirID-CRBN (full-length form) or ThBD-AGIA-AirID were treated with or without ARV-825. Immunoblotting after STA-PDA showed that ThBD-AGIA-AirID induced biotinylation of BRD3 and IKZF1 in the presence of ARV-825 (Fig. 3g). In contrast, in AirID-CRBN-expressing cells, although IKZF1 biotinylation was detected, the band intensity was lower than that in ThBD-AGIA-AirID-expressing cells, and BRD3 biotinylation was not detectable. This difference in BRD3-biotinylation activity between full-length CRBN and ThBD-AGIA-AirID is apparently attributable to the higher expression level of ThBD-AGIA-AirID than that of full-length CRBN (arrowheads in IB: BirA panel of Fig. 3g).

To determine whether ThBD-AGIA-AirID could be used in non-human cell lines, we used the murine melanoma-derived B16F10 cell line. In the mouse cells, ThBD-AGIA-AirID biotinylated BRD3 in the presence of ARV-825 (Fig. 3h). Taken together, these results indicate that ThBD-AGIA-AirID is suitable for PROTAC-dependent biotin labelling in cells and can be used in animal cells other than human cell lines. Integration with the experimental results for AirID-fused VHL showed that ThBD-AGIA-AirID was suitable for the PROTAC-dependent biotinylation of POI; therefore, ThBD-AGIA-AirID was used as the AirID-fused E3 ligand domain in subsequent experiments.

### ARV-825-dependent intracellular biotinylated interactome analysis using ThBD-AGIA-AirID and LC-MS/MS

Since ThBD-AGIA-AirID was functional in cells, it was used for exploring ARV-825-dependent intracellular biotinylated interactome. MM1.S cells stably expressing ThBD-AGIA-AirID were treated with ARV-825 and biotin, and cell lysates were analysed via LC-MS/MS. DMSO and pomalidomide were used as negative controls to confirm ARV-825-independent biotinylation. LC-MS/MS analysis showed that biotinylation of BRD3, IKZF1, IKZF3, WIZ, and ZFP91 was induced by ARV-825 treatment but not by DMSO treatment (Fig. 4a, Supplementary Data 2). Comparison between pomalidomide- and DMSO-treated cells showed IMiD-dependent biotinylation of neosubstrates in CRBN, whereas not of BRD3 (Fig. 4b, Supplementary Data 2). Comparison of biotinylated proteins between ARV-825- and pomalidomide-treated cells showed that ARV-825 significantly induced biotinylation of BRD3; the ratio for the ARV-825 to pomalidomide treatment significantly reduced the biotinylation ratio of IKZF1, IKZF3, WIZ, and ZFP91 (Fig. 4c, Supplementary Data 2), which are neosubstrates for IMiD-dependent CRBN^38,39,40^. CRBN and VHL are conserved proteins that function as substrate recognition adapters in the CRL4^CRBN^ and CRL2^VHL^ E3 ubiquitin ligase complex, respectively^41,42^. ThBD is not thought to integrate into the CRL4^CRBN^ complex because it lacks the DDB1-binding domain of CRBN. Therefore, we examined whether biotinylated peptides derived from CUL4, DDB1, and RBX1 could be detected using ThBD-AGIA-AirID (Supplementary Fig. 3a). Six CRL4 complex-derived biotinylated peptides were detected by AirID-CRBN and three CRL2 complex-derived biotinylated peptides were detected by VHL-AirID. Compared with AirID-CRBN and VHL-AirID, ThBD-AGIA-AirID only detected only the biotinylated peptide K8 of CUL4A, which was the biotinylated site detected in all treated samples. Therefore, the K8 site was considered to have background biotinylation rather than specific biotinylation. These results indicate that ThBD is not incorporated into the CRL4 complex. Furthermore, to compare biotinylated interactomes induced by PROTACs with the same target binder but different E3 binders, we analysed the results for ThBD-AGIA-AirID and AirID-CRBN with ARV-825 treatment, and VHL-AirID with MZ1 treatment. PROTAC-dependent biotinylated proteins were defined as biotinylated peptides with an abundance ratio >2 and a *P*-value <0.05 compared to the treatment with competitive compound. Twenty-three ARV-825-dependent biotinylated proteins were identified for ThBD-AGIA-AirID using this threshold (Supplementary Fig. 3b). To determine whether biotinylation by ThBD-AGIA-AirID allowed analysis of the ARV825-dependent BRD4 protein interactome, the 23 proteins were mapped based on protein–protein interaction data from the STRING database (Supplementary Fig. 3c)^32^. Mapping represented only interactions with other proteins, and the results showed that five BRD4-interacting proteins were biotinylated by ThBD-AGIA-AirID in an ARV-825-dependent manner. Venn diagrams were generated for the 23 biotinylated proteins and compared with the PROTAC-dependent proteins biotinylated by AirID-CRBN and VHL-AirID (Fig. 4d). Four PROTAC-dependent biotinylated proteins, including BRD3, were common among the AirID-fused CRBN, ThBD, and VHL. In addition, one and three of the 23 proteins were common between ThBD and CRBN and ThBD and VHL. Interestingly, there were a maximum of 35 proteins between full-length CRBN and VHL, suggesting that these full-length forms have several domains that interact with other proteins to form the E3 complex.

**Fig. 4.**
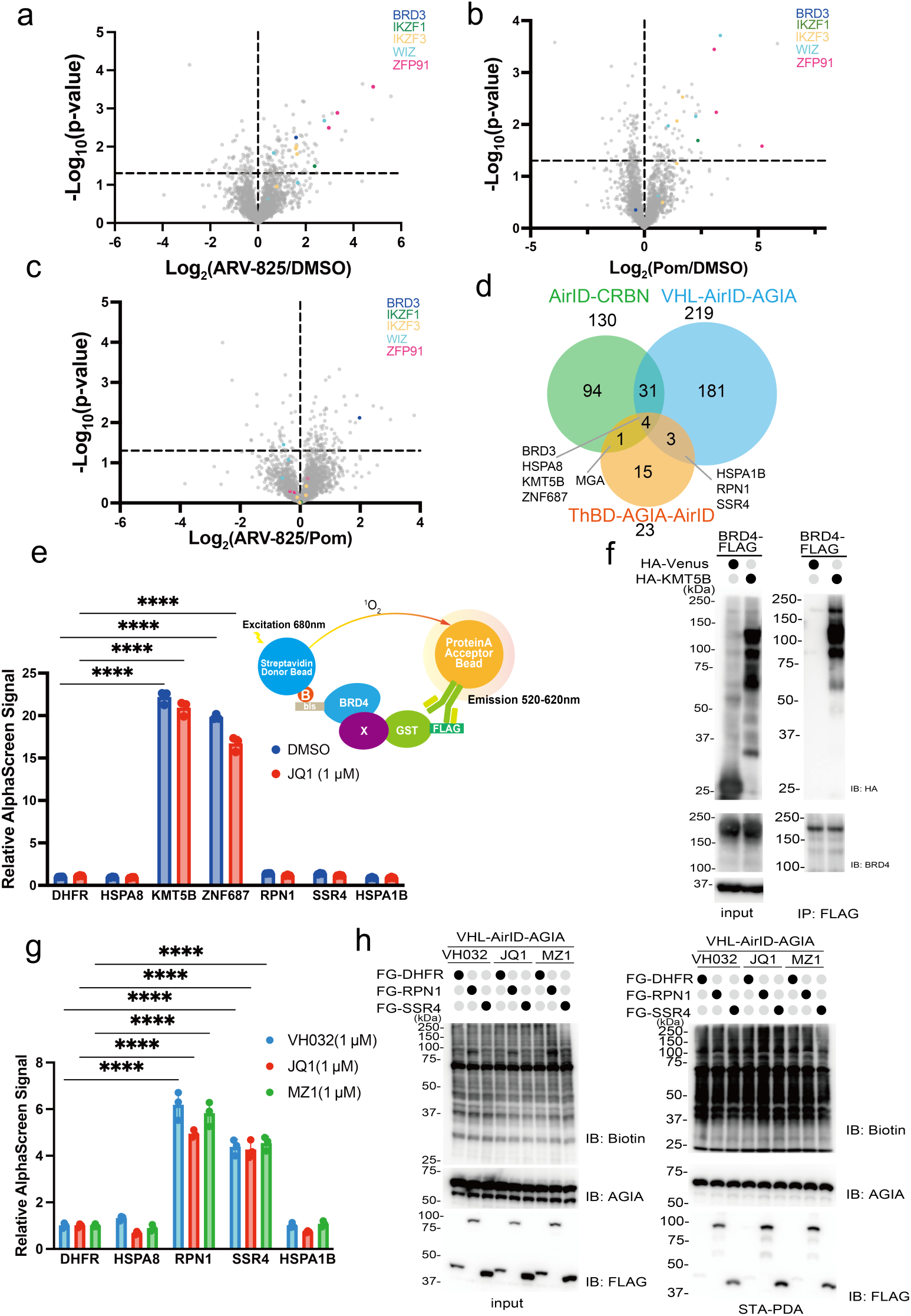
Comparison of in-cell proximity of biotinylated proteins among ThBD-AGIA-AirID, AirID-CRBN, and VHL-AirID with PROTAC. **a, b, c** Volcano plots of ARV-825-dependent biotinylated peptides detected via LC-MS/MS in MM1.S cells stably expressing ThBD-AGIA-AirID. Significant changes in the volcano plots were calculated using Student’s two-sided *t*-test, and the false discovery rate (FDR)-adjusted *P*-values calculated using Benjamini–Hochberg method are shown in Supplementary Data 2. Volcano plot comparing a ARV-825 treatment with DMSO treatment, b Pomalidomide treatment with DMSO treatment, c ARV-825 treatment with Pomalidomide. **d** Venn diagram of ARV-825- or MZ1-dependent proteins biotinylated by ThBD-AGIA-AirID, AirID-CRBN, or VHL-AirID (PROTAC/E3 ligand ratios of biotinylated peptides detected via LC-MS/MS >2 and *P* < 0.05). **e** AlphaScreen assay detection of bls-BRD4 interaction protein. All bls-BRD4 interaction analyses with candidate BRD4-interacting proteins were performed in the presence of DMSO or JQ1. All AlphaScreen signals represent the relative values of FG-DHFR and bls-BRD4 proteins. Error bars denote standard deviations (independent experiments, *n* = 3). Statistical significance was determined using two-way ANOVA with Dunnett’s multiple comparison test (\*\*\*\**P* < 0.0001). Error bars represent standard deviation. **f** HEK293T cells were transfected with each gene. The transfected Venus gene was used as a control. Immunoprecipitation was performed using anti-FLAG antibody. The immunoprecipitated proteins were detected using anti-FLAG and anti-BRD4 antibodies. **g** AlphaScreen detection of proteins biotinylated by VHL-AirID in vitro. All AlphaScreen signals represent the relative values of FG-DHFR and VHL-AirID for each drug treatment. Error bars denote standard deviations (independent experiments, *n* = 3). Statistical significance was determined using two-way ANOVA with Dunnett’s multiple comparison test (\*\*\*\**P* < 0.0001). **h** Streptavidin pull-down detection of RPN1 or SSR4 biotinylated by VHL-AirID in vitro.

To confirm the interaction of their common proteins, six common proteins, HSPA8, KMT5B, and ZNF687 among CRBN, ThBD, and VHL, and HSPA1B, RPN1, and SSR4 between ThBD and VHL, were selected for further analysis. The proteins were synthesised using a cell-free system. Expression and band size of the synthesised proteins were confirmed using immunoblotting (Supplementary Fig. 4a). First, direct interactions of the synthesised proteins with BRD4 were assessed using the AlphaScreen method in the presence and absence of JQ1 (Fig. 4e). KMT5B and ZNF687 directly interacted with BRD4 without the influence of JQ1, indicating that the binding of JQ1 to BRD4 even if they were bound did not affect their interaction. We used a co-immunoprecipitation assay to investigate whether KMT5B forms a complex with BRD4 in cells (Fig. 4f). The results suggested that BRD4 and KMT5B bind directly not only in vitro but also in cells. Second, the four remaining proteins were investigated for direct interactions with VHL using the AlphaScreen method in the presence of VH032, JQ1, or MZ1. For this analysis, a biotinylation assay using AirID-fused VHL was performed (Supplementary Fig. 4b). In this assay, proteins are biotinylated by VHL-AirID upon binding to VHL. Thus, in this case, AlphaScreen detects biotinylation of proteins after the binding of VHL-AirID. The AlphaScreen signals showed that VHL-AirID biotinylated RPN1 and SSR4 without the influence of VH032, JQ1, or MZ1 (Fig. 4g). In vitro biotinylation of RPN1 and SSR4 was confirmed using immunoblotting after STA-PDA (Fig. 4h). Furthermore, we investigated whether these four proteins could be biotinylated by AirID-CRBN or ThBD-AGIA-AirID in the presence of pomalidomide, JQ1, or ARV-825. No biotinylation was detected using the AlphaScreen method, although the positive control BRD4 was biotinylated in an ARV-825-dependent manner (Supplementary Fig. 4b, c, d), indicating that RPN1 and SSR4 did not interact with CRBN and ThBD. Taken together, these results indicate that RPN1 and SSR4 are novel direct interactors of VHL.

### EGFR-targeting PROTAC-dependent biotinylation by AirID-fused ThBD and VHL

EGFR is a widely recognised oncogene and an important drug target^43,44^. Therefore, multiple tyrosine kinase inhibitors (TKIs) targeting EGFR, such as gefitinib, are widely used for cancer treatment^45,46^. However, TKIs targeting EGFR are known to cause the emergence of drug-resistant cancer cells with continued dosing^47^. Proteolysis-inducing drugs are expected to be used as EGFR-targeting drugs to prevent resistance. Currently, two EGFR-targeting PROTACs, MS154 and MS39, have been developed with gefitinib as an EGFR binder and thalidomide backbone or VHL ligand as E3 binder, respectively (Fig. 5a)^48^. The proteins with which MS154 and MS39 interact in cells are poorly understood. In addition, there are a few examples of the identification of proteins other than EGFR with which gefitinib interacts in cells. Gefitinib causes several side effects such as interstitial pneumonia^49^. Identification of proteins to which gefitinib binds in cells may lead to an understanding of the mechanism of its side effects. Therefore, we analysed the interactome induced by MS154 or MS39 and the proteins that bind to gefitinib in these cells. HCC-827-Luc cells, which are human lung cancer-derived cells expressing luciferase, were used for analysis of MS154 and MS39. HCC-827 cells express EGFR del19, which is deleted at exon 19 of EGFR^50^. Gefitinib and TKIs targeting EGFR are known to bind to the kinase domain of these EGFR mutants better than to the wild-type EGFR. EGFR degradation by MS154 and MS39 was also detected in HCC-827 cells^48^. For these reasons, cell lines expressing ThBD-AGIA-AirID or VHL-AirID were established in HCC-827-Luc cells to search for interactomes in proximity to MS154 or MS39 and proteins that bind to gefitinib. Biotinylation reactions were performed in HCC-827-Luc cells expressing ThBD-AGIA-AirID by the addition of pomalidomide or MS154 (Fig. 5b). EGFR was detected using STA-PDA only in the MS154-treated section. This result indicated that ThBD-AGIA-AirID can biotinylate MS154 target proteins in HCC-827-Luc cells. Next, HCC-827-Luc cells expressing VHL-AirID or VBD-AGIA-AirID were established, and MS39-dependent biotinylation of EGFR was observed using STA-PDA (Fig. 5c). Biotinylated EGFR could only be detected with MS39-supplemented treatment of VHL-AirID-expressing HCC-827-Luc cells, but not of VBD-AGIA-AirID-expressing cells. This result is consistent with the higher efficiency of POI biotinylation in cells with VHL-AirID than in those with VBD-AGIA-AirID, as shown in Figure 2.

**Fig. 5.**
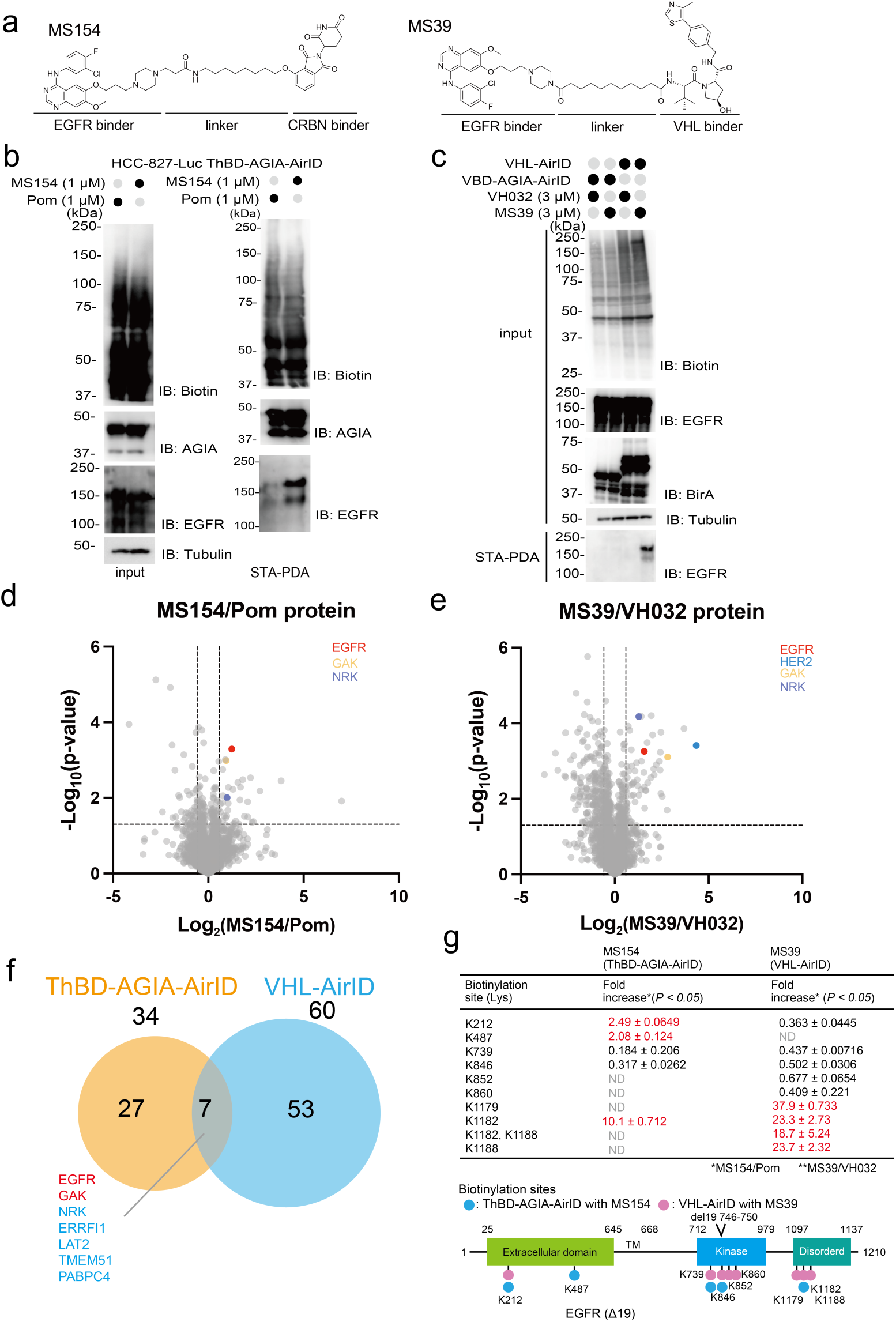
Analysis of proteins biotinylated by ThBD-AGIA-AirID or VHL-AirID in the presence of EGFR-targeted PROTACs MS154 and MS39. **a** Chemical structures of MS154 and MS39. **b** Streptavidin pull-down assay of MS154-dependent biotinylation of EGFR in HCC827-Luc cells stably expressing ThBD-AGIA-AirID. **c** Streptavidin pull-down assay of MS39-dependent biotinylated EGFR in HCC827-Luc cells stably expressing VBD-AGIA-AirID or VHL-AirID. **d**, **e** Volcano plots of MS154- or MS39-dependent biotinylated proteins detected via LC-MS/MS in MS154 or in HCC827-Luc cells stably expressing ThBD-AGIA-AirID or VHL-AirID. Significant changes in the volcano plots were calculated using Student’s two-sided *t*-test, and the false discovery rate (FDR)-adjusted *P*-values calculated using the Benjamini–Hochberg method are shown in Supplementary Data 3 and 5. d, Volcano plot diagram comparing MS154 treatment with pomalidomide treatments; e, Volcano plot diagram comparing MS39 and VH032 treatments. The vertical dashed lines represent a ratio of 1.5 and −1.5, respectively. **f** Venn diagram of MS154- or MS39-dependent protein biotinylation by ThBD-AGIA-AirID or VHL-AirID (PROTAC/E3 ligand ratios of biotinylated proteins detected via LC-MS/MS >1.5 and *P* < 0.05). Red text represents known interacting proteins of gefitinib. **g** Table and schematic diagram of EGFR lysine residues biotinylated in a ThBD-AGIA-AirID- or VHL-AirID-dependent manner by MS154 or MS39 treatment. The red text in the table represents EGFR lysine residues with MS154- or MS39-dependent biotinylation. The schematic diagram shows full-length EGFR del 19, with blue circles representing MS154-dependent biotinylation sites and pink circles representing MS39-dependent biotinylation sites.

### Comparison of MS154- and MS39-induced interactomes using AirID-fused ThBD and VHL via LC-MS/MS

Next, we verified the biotinylated interactome induced by MS154 and MS39 treatment using HCC-827-Luc cells stably expressing ThBD-AGIA-AirID or VHL-AirID, respectively. EGFR degradation by MS154 is accelerated in serum-free medium^48^. Therefore, the medium was replaced with serum-free medium prior to MS154 and biotin supplementation to prevent EGFR degradation. LC-MS/MS results confirmed MS154-dependent biotinylation of EGFR by comparison with pomalidomide- and DMSO-treated compartments (Fig. 5d, Supplementary Fig. 5a, b, c, Supplementary Data 3, 4). Gefitinib binds to cyclin G-associated kinase (GAK) and EGFR^51,52^. Biotinylated GAK was also increased in the MS154-treated compartment, demonstrating that the biotinylation of PROTAC-target proteins using ThBD-AGIA-AirID is useful for target identification. Next, the biotinylated interactome for VHL-AirID with MS39 was analysed. For MS39 analysis, cell lysates were obtained from cells cultured in biotin, MG132 and serum-containing medium^48^. Similar to that in MS154 treatment, LC-MS/MS of MS39-treated cell lysates also showed biotinylation of EGFR and GAK in comparison with VH032- and DMSO-treated sections (Fig. 5e, Supplementary Fig. 5d, Supplementary Data 5). Notably, in the MS39-induced biotinylated proteins, the ERBB family member HER2 was detected, although it was not included in the MS154-induced interactome. This difference indicated that EGF ligands in the serum under MS39-treated conditions induced the interaction between EGFR and HER2, which is widely known to be induced by EGF treatment. As seen from the volcano plots of biotinylated peptides, many EGFR and GAK lysine residues were biotinylated (Supplementary Fig. 5e, f, Supplementary Data 6). Next, the correlation between MS154- and MS39-dependent biotinylation of proteins was investigated using a Venn diagram. PROTAC-dependent biotinylated proteins were defined as proteins with a *P*-value <0.05, and a ratio to pomalidomide or VH032 >1.5. The number of MS154- and MS39-dependent biotinylated proteins was 35 (Supplementary Fig. 5g) and 64 (Supplementary Fig. 5h), respectively. GO analysis of MS154- and MS39-dependent biotinylated proteins using ShinyGO showed enrichment of terms related to kinase activation and kinase binding (Supplementary Fig. 5, j)^33^. These results indicated that biotinylated proteins were enriched for EGFR-associated proteins. The Venn diagram shows seven common proteins, including three protein kinases, EGFR, GAK, and Nik-related kinase (NRK) (Fig. 5f). Notably, EGFR receptor feedback inhibitor 1 (ERRFI1) interacts with EGFR family proteins to suppress kinase activity by inhibiting dimerization^53,54^. ERRFI1 was a common biotinylated protein induced in MS154 and MS39 treatment, suggesting that it may interact with EGFR upon gefitinib treatment and promote the suppression of EGFR signalling. Further studies on the roles of NRK and ERRFI1 in the processing of EGFR-targeting PROTAC and gefitinib drugs are required to better understand the mechanisms of these drug effects.

To understand biotinylation by ThBD-AGIA-AirID and VHL-AirID, each biotinylation site in EGFR was compared. In the intracellular region, seven biotinylation sites were identified: four in the kinase domain and three in the C-terminal disorder domain (Fig. 5g). Of the biotinylated sites, only three were present in ThBD treated with MS154, K739, and K846 within the kinase domain and K1182 in the disorder domain. This number was lower than that for VHL. Three sites, K1179, K1182, and K1186, within the disorder domain were significantly increased by MS154 or MS39 treatment, suggesting that these EGFR-targeting PROTACs may expose surface lysine sites and/or induce conformational changes in the disorder domain of EGFR. Surprisingly, two biotinylation sites were found in the extracellular region; however, further studies are required to understand the underlying reasons. Taken together, these results show that the known gefitinib-interacting proteins are biotinylated in an MS154- and MS39-dependent manner, indicating that ThBD-AGIA-AirID and VHL-AirID function as expected. Furthermore, ThBD-AGIA-AirID and VHL-AirID can be used for interactome analysis of EGFR-targeting PROTACs.

### Neosubstrate exploration of VHL-binder VH032

Thalidomide and its derivatives function as molecular glue and induce neosubstrates ^38,39^. Although VH032 is a commonly used VHL ligand, no neosubstrate are known to induce interaction with VH032. The AirID-fusion system could successfully identify the neosubstrates of thalidomide and its derivatives in previous^26^ and this study (Fig. 4). Therefore, we investigated whether the neosubstrate of VH032 was present in the LC-MS/MS data. In this study, the VH032 single treatment group was also analysed using MS of PROTAC-dependent biotinylation of proteins by VHL-AirID; therefore, we focused on proteins that were biotinylated in a VH032-dependent manner from the LC-MS/MS data. Biochemical assays using the AlphaScreen method confirmed the interaction between the target proteins and VHL. In MM1.S and HCC827-Luc cells, the number of proteins biotinylated by VH032 in a VHL-AirID-dependent manner was 2 and 11, respectively (Supplementary Fig. 6a, b). Of these, SCRN3 was not included in biochemical assays because it was unlikely to be biotinylated in a VH032-dependent manner, as its ratio value compared to MZ1 and VH032 was >2. Seven proteins were selected from the VH032-dependent biotinylated proteins for direct biochemical interaction analysis using the AlphaScreen method (Supplementary Fig. 6c). Interaction analysis was performed in addition to the RPN1 and SSR4 treatments, which were found to interact directly with VHL. These proteins were synthesised using a cell-free system, and their expression and band size were confirmed by immunoblotting (Supplementary Fig. 6d). The AlphaScreen results showed that SLC25A5, COPB2, MAD2L1BP, PARG, TIPRL, ZNF114, RPN1, and SSR4 bind directly to VHL (Supplementary Fig. 6e), and all interactions were unaffected by VH032 treatment, indicating that the selected proteins are not neosubstrates of VH032. These results indicated that the neosubstrates of VH032 were not present among the biotinylated proteins derived from MM1.S and HCC827-Luc cells. Neosubstrates of the CRBN binder were found in various cells; however, those of the VHL binder were not found in the mass spectrometry results in this study, suggesting that the VHL binder is mainly responsible for inducing the degradation of target proteins. This is a major advantage of using a VHL binder in the development of PROTACs.

### Biotinylation analysis of a heterobifunctional molecule consisting of geldanamycin and thalidomide derivative

Recently, heterobifunctional molecules other than PROTAC, such as DUBTAC and PhosTAC, have been developed^55,56,57^. Therefore, the analysis of proteins that interact with heterobifunctional compounds other than PROTAC is useful for drug development beyond TPD. In addition, during PROTAC development, nondegradable heterobifunctional molecules can be produced. This indicates that the developing heterobifunctional molecule cannot ubiquitinate the target protein. However, it is important in PROTAC development to determine why the developing heterobifunctional molecules does not bind to the target protein, the E3 ligase, or cross the membrane in the beginning. Therefore, we investigated whether AirID-fused ThBD could be used to analyse nondegradable heterobifunctional molecules. As CRBN binders are essential for the use of AirID-fused ThBDs, the heterobifunctional molecules to be evaluated use thalidomide derivatives (pomalidomide) as binders. Geldanamycin (GA), known to bind HSP90^58^, was used as the target binder. Click chemistry is a widely used method for the fusion of two chemical molecules^59^. Moreover, because thalidomide derivatives for click chemistry are commercially available, it was used to synthesise heterobifunctional molecules. Using click chemistry, heterobifunctional molecules (GA-TD), with five types of linkers, were synthesised (Fig. 6a). First, to investigate whether these GA-TDs induced an interaction between HSP90 and ThBD, five GA-TD, numbered 1, 3, 5, 6, and 7, were validated using the AlphaScreen method. After the synthesis of GST-bls-ThBD (bls: biotin ligation site) and FLAG-HSP90AA1 using the wheat cell-free protein synthesis system, these proteins were mixed with individual GA-TDs, and the signals were detected using the AlphaScreen assay. The highest AlphaScreen signal was detected in the 1 µM GA-TD-5 treatment (Fig. 6b). In addition, we observed the hook effect, which is specific to heterobifunctional molecules, with GA-TD-1, -3, and -5, whereas GA-TD-6 and -7 were not different from the negative control pomalidomide and no interaction was detected. Next, the ability of GA-TDs to induce the biotinylation of HSP90 in HEK293T cells stably expressing ThBD-AGIA-AirID was validated. After 6 hours, of GA-TD and MG132 addition to cell medium, the cells were lysed, and biotinylation of HSP90 was detected using STA-PDA. The STA-PDA results showed that HSP90 was biotinylated in GA-TD-5-, and GA-TD-6-treated cells compared to the DMSO-treated cells (arrowhead in Fig. 6c). As the AlphaScreen signal value of GA-TD-6 in Fig. 6b was low, GA-TD-5 were used for further analysis. IKKα is reduced in a geldanamycin-dependent manner^60^. We investigated whether GA-TD-3 and GA-TD-5 affect the expression levels of HSP90 and IKKα in cells using immunoblotting. At 10 µM, GA-TD-5 induced a decrease in IKKα levels, as did GA (Fig. 6d), whereas GA-TD-3 had no effect to IKKα levels. However, neither GA-TD-3 nor GA-TD-5 induced HSP90 degradation. Furthermore, cell proliferation rates were measured using the Cell-Titer-Glo assay to investigate the cytotoxicity of GA-TD-5 in HEK293T cells. Both GA-TD-3 and GA-TD-5 were more cytotoxic than pomalidomide at high concentrations (Fig. 6e), although the cytotoxicity was highest with GA. Interestingly, although GA-TD-3 showed cytotoxicity similar to that of GA-TD-5, it did not induce HSP90 biotinylation in cells (Fig. 6d, e). The inability of GA-TD-3 to induce IKKα degradation and HSP90 biotinylation suggests that it may not retain the functional properties of GA in cells. Further research is needed regarding the cytotoxicity of GA-TD-3 in cells. Next, we investigated whether GA-TD-5 could be used to check the degradation of HSP90 in detail, using a cycloheximide chase assay. However, GA-TD-5 did not degrade HSP90 in this assay (Fig. 6f). These results indicated that GA-TD-5 cannot degrade HSP90. Taken together, these results show that GA-TD-5 induces the interaction between HSP90 and ThBD-AGIA-AirID but not the degradation of HSP90.

**Fig. 6.**
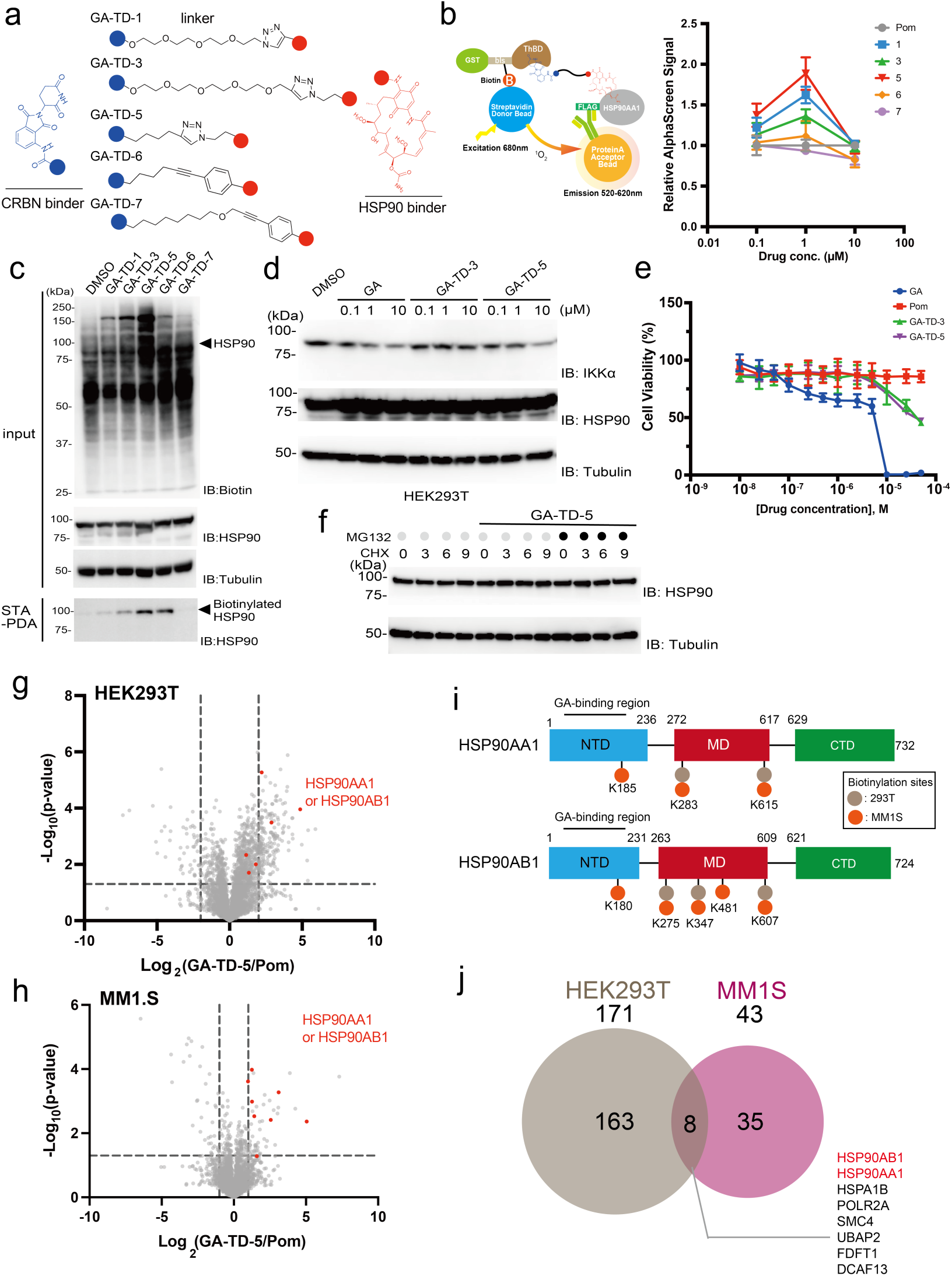
Proximity biotinylation of heterobifunctional molecule GA-TD with different linkers. **a** Chemical structures of GA-TD-1 through 7. **b** AlphaScreen detection of the interaction between ThBD and HSP90AA1 via GA-TD. Error bars denote standard deviations (independent experiments, *n* = 3). All relative AlphaScreen signals represent the relative values for each pomalidomide treatment. **c** Streptavidin pull-down detection of GA-TD-dependent HSP90 biotinylated by AirID-fused ThBD in HEK293T cells stably expressing ThBD-AGIA-AirID. GA-TDs were subjected to the biotinylation reaction at a concentration of 1 µM. **d** Immunoblot analysis of IKKα and HSP90. HEK293T cells were treated with geldanamycin (GA), GA-TD-3, or GA-TD-5 for 6 h. **e** Dose–response curve of the antiproliferative effect of GA-TDs on HEK293T cells. HEK293T cells were treated with DMSO, GA, GA-TD-3, or GA-TD-5 for 24 h and cell viability was analysed using the CellTiter-Glo assay kit. Cell viability was expressed as the luminescence signal relative to the luminescence signal of DMSO, which was considered as 100. Error bars denote standard deviations (biological replicates; *n* = 3). **f** Effect of GA-TD-5 on HSP90 protein levels. HEK293T cells were treated with DMSO or 10 µM GA-TD-5 in the presence of 100 µg/mL cycloheximide (CHX). Five micromolar MG132 or DMSO was added together with GA-TD-5 and cells were collected at indicated times. **g**, **h** Volcano plots of GA-TD-5-dependent biotinylated peptides detected via LC-MS/MS in HEK293T or MM1.S cells stably expressing ThBD-AGIA-AirID. The vertical dashed lines in volcano plot of HEK293T cells represent Log_2_(GA-TD-5/Pom) of 2 or −2, respectively. The vertical wavy lines in volcano plot of HEK293T cells represent Log_2_(GA-TD-5/Pom) of 2 or −2, respectively. The vertical dashed lines in the volcano plot of MM1.S cells represent Log_2_(GA-TD-5/Pom) of 1 or −1, respectively. Significant changes in volcano plots were calculated using Student’s two-sided *t*-test, and the false discovery rate (FDR)-adjusted *P*-values calculated using Benjamini–Hochberg method are shown in Supplementary Data 7, 8. **i** Schematic of HSP90 lysine residues biotinylated in a ThBD-AGIA-AirID-dependent manner following GA-TD-5 treatment. The schematic shows HSP90AA1 and HSP90AB1, with grey circles representing biotinylation sites in HEK293T cells and orange circles representing those in MM1.S cells. **j** Venn diagram of GA-TD-5-dependent proteins biotinylated by ThBD-AGIA-AirID in HEK293T and MM1.S cells (HEK293T cells: GA-TD-5/Pom ratios of biotinylated peptides detected via LC-MS/MS >4 and *P* < 0.05, MM1.S cells: GA-TD-5/Pom ratios of biotinylated peptides detected via LC-MS/MS >2 and *P* < 0.05).

### Analysis of GA-TD-5 or -3-dependent biotinylated interactome using ThBD-AGIA-AirID and LC-MS/MS

The experiments described above revealed that GA-TD-5 is a nondegradable heterobifunctional molecule and GA-TD-3 is a heterobifunctional molecule that can neither degrade nor biotinylate its target protein. Therefore, interactome analysis of these two nondegradable compounds was performed. HEK293T and MM1.S cells stably expressing ThBD-AGIA-AirID were used to comprehensively analyse biotinylated proteins in cells treated with GA-TD-5 or GA-TD-3. As a negative control, cells expressing ThBD-AGIA-AirID were treated with DMSO or pomalidomaide, together with MG132 and biotin. After incubation for 6 hours, biotinylated proteins in the cells were analysed via LC-MS/MS. Volcano plots were generated by comparing the biotinylation rates of cells treated with GA-TD-5 and those treated with pomalidomide (Pom) or DMSO in each cell line (Fig. 6g, h, Supplementary Fig. 7a, b, Supplementary Data 7, 8). In both cell lines, GA-TD-5 significantly induced the biotinylation of HSP90 (HSP90AA1 and HSP90AB1, red spots), whereas GA-TD-3 did not significantly induce them (Supplementary Fig. 7c, d). Biotinylation sites were detected in both HSP90AA1 and HSP90AB1 (Fig. 6i). The biotinylated sites of HSP90AA1 and HSP90AB1 were concentrated in the N-terminal ATPase domain (NTD) and the middle domain (MD), indicating that AirID-fusion ThBD mainly biotinylates the proximity of the GA-binding site. Next GA-TD-5-dependent biotinylated proteins from both the cell lines were analysed (Supplementary Fig. 7e, f). For HEK293T cells stably expressing ThBD-AGIA-AirID, biotinylated proteins had a (GA-TD-5/Pom) ratio >4 and *P*-value <0.05. At this threshold, 171 proteins were biotinylated in HEK293T cells. 43 proteins were biotinylated in MM1.S cells in a GA-TD-5-dependent manner, defined as biotinylated proteins with a (GA-TD-5/Pom) ratio >2 and *P*-value <0.05. A Venn diagram was created to identify the common biotinylated proteins (Fig. 6j). Eight proteins were biotinylated, including HSPA1B, which interacts with HSP90AA1, were identified^61,62^. In addition, a co-immunoprecipitation assay with HSP90 confirmed that DCAF13, an E3 ligase, interacted with HSP90 in cells (Supplementary Fig. 7g). These results and those of previous studies showed that proteins GA-TD-5-dependently biotinylated in the two cell lines stably expressing ThBD-AGIA-AirID include proteins that interact directly with HSP90. Taken together, AirID-fused ThBD was useful for the analysis of nondegradable heterobifunctional molecules. These findings suggest that AirID-fusion proteins could be useful for the development of protein–protein-induced heterobifunctional molecules, in addition to degradative agents such as PROTAC.

### ARV-110-dependent interactome analysis using ThBD-AGIA-AirID

ARV-110 is an oral PROTAC that targets the AR in the treatment of metastatic castration-resistant prostate cancer (mCRPC)^7^ and is a compound undergoing clinical trials (Fig. 7a). The DC_50_ of ARV-110 is 1 nM and it acts as a potent AR-degrading agent^6^. In the AR-degradation process, ARV-110-dependent interactions have not been investigated. AR interacts with a variety of proteins within cells^63,64^. Furthermore, as ARV-110 interacts with AR in cells, analysis of the interacting proteins induced by ARV-110, including AR in cells, is expected to provide useful information for determining the intracellular localisation where the ARV-110–AR interaction occurs and for considering the intracellular drug mechanism, such as whether ARV-110–AR interacts with other proteins. This information may allow us to investigate which drugs in combination with AR-targeted PROTACs, such as ARV-110, may efficiently inhibit mCRPC proliferation. Therefore, we used AirID-fused ThBD-expressing LNCaP cells to investigate ARV-110-dependent proximity biotinylation of AR. First, ARV-110 or pomalidomide was added to ThBD-AGIA-AirID-expressing LNCaP cells together with MG132 and biotin to determine whether the AR was biotinylated using STA-PDA (Fig. 7b). ThBD-AGIA-AirID biotinylated AR in an ARV-110-dependent manner. The ability of ARV-110 to degrade AR was also monitored using western blotting, to determine whether ARV-110 retained AR degradation when expressing ThBD-AGIA-AirID (Fig. 7c). These results confirmed that AR is also degraded in an ARV-110-dependen manner in LNCaP cells expressing ThBD-AGIA-AirID and that this degradation is inhibited by MG132. Next, we identified the biotinylated proteins via LC-MS/MS to observe ARV-110-dependent biotinylation by ThBD-AGIA-AirID in LNCaP cells (Fig. 7d, e, Supplementary Data 9, 10). Initially, a volcano plot focusing on the biotinylated peptides was generated, and four biotinylated peptides from AR were identified (red spots in Fig. 7d). Three of these four peptides were predominantly biotinylated by ARV-110. Four of these biotinylated sites are shown in the structural schematic for AR, which indicates that the N-terminus rather than the ligand-binding domain (LBD) is biotinylated (Fig. 7f). In addition, protein-focused volcano plots showed that biotinylation of AR and KAT7^65^, NFIB^66^, GRHL2^67^, MED1^68^, NCOA2^69^, BRD4^70^, FOXA1^71^ and SART3^72^, which are known to interact with AR, occurred via ARV-110 (Fig. 7e), suggesting that ARV-110 interacts with AR and other proteins. Interestingly, all eight AR-interacting proteins localise in the nucleus, which indicates that ARV-110 binds to nuclear ARs. In addition, 161 proteins were biotinylated in ARV-110-dependent manner in ThBD-AGIA-AirID-expressing LNCaP cells (Supplementary Fig. 8a). The biotinylated proteins were subjected to GO analysis using ShinyGO; the GO cellular component category was predominantly enriched in nucleus-related terms in an ARV-110-dependent manner (Fig. 7g). In addition, in the biological process and molecular function categories, RNA processing and RNA binding terms were enriched in an ARV-110-dependent manner. These results indicated that ARV-110-dependent biotinylation by ThBD-AGIA-AirID reflects the localisation of ARs in the nucleus (Supplementary Fig. 8b, c). Moreover, ThBD-AGIA-AirID was able to biotinylate targets and proteins that interacted with target proteins for drugs in clinical trials.

**Fig. 7.**
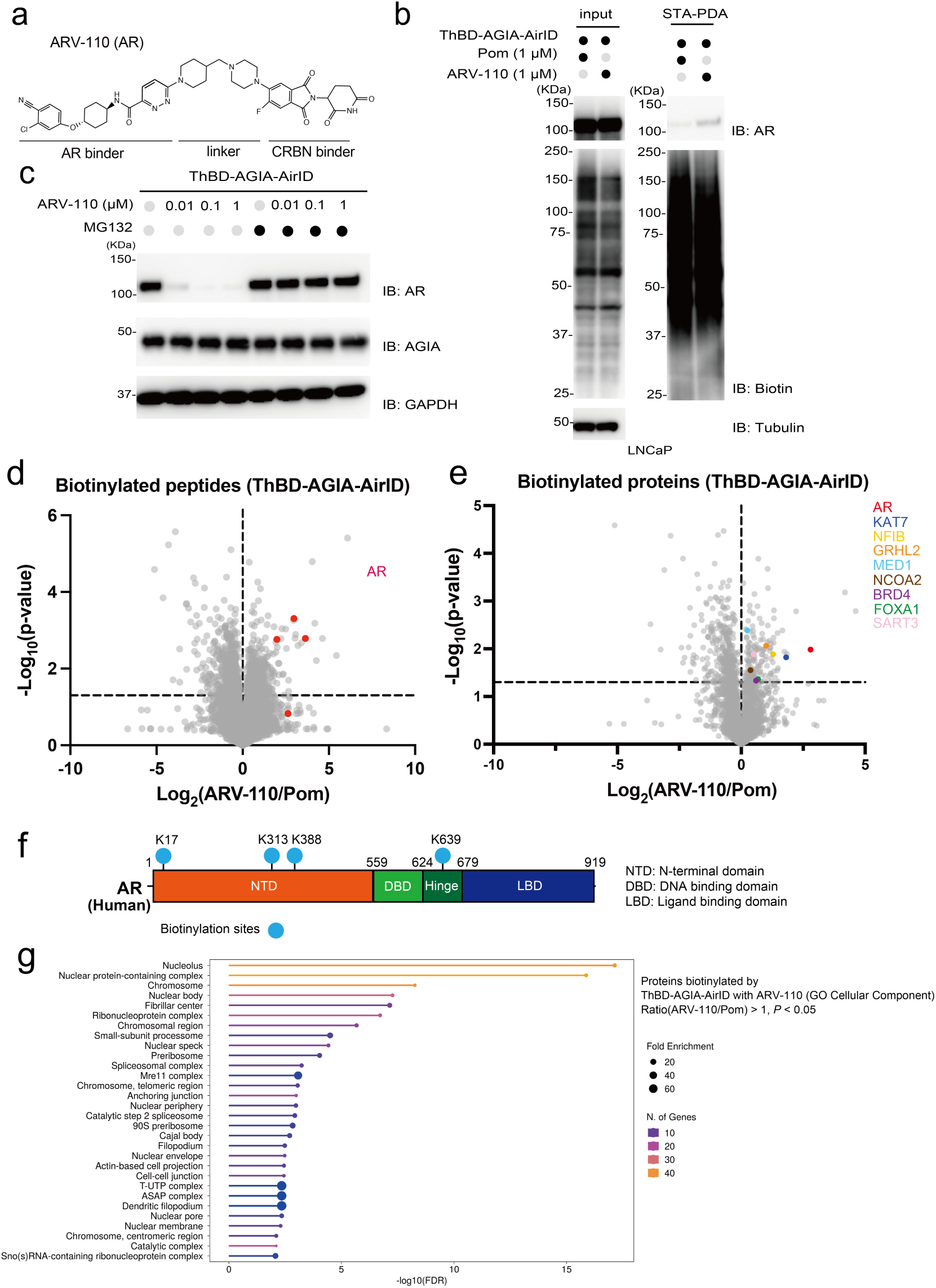
LC-MS/MS analysis of ARV-110-dependent proximity biotinylation by ThBD-AGIA-AirID. **a** Chemical structure of ARV-110. **b**, Streptavidin pull-down assay of ARV-110-dependent biotinylated AR in LNCaP cells stably expressing ThBD-AGIA-AirID. **c** Immunoblot analysis of ARV-110-dependent AR degradation. Cells were collected 6 h after adding the indicated concentrations of ARV-110 or DMSO together with 5 µM MG132 or DMSO to the medium. **d**, **e** Volcano plots of ARV-110-dependent biotinylated peptides and proteins detected via LC-MS/MS in LNCaP cells stably expressing ThBD-AGIA-AirID. d, Volcano plot of biotinylated peptides comparing ARV-110 treatment with pomalidomide treatment; e, Volcano plot of biotinylated proteins. Significant changes in the volcano plots were calculated using Student’s two-sided *t*-test, and the false discovery rate (FDR)-adjusted *P*-values calculated using Benjamini–Hochberg method are shown in Supplementary Data 9, 10. **f** Schematic showing AR, with light blue circles representing biotinylation sites. **g** Gene Ontology (GO) analysis of proteins biotinylated by ThBD-AGIA-AirID dependent on ARV-110 in LNCaP cells. GO analysis was performed using the ShinyGO software. The horizontal axis represents −log_10_ (FDR), the colour change represents the number of genes, and the size of the circle represents fold enrichment. GO cellular component analysis was performed.

## Discussion

PROTACs, molecules that are chemical fusions of an E3 binder and a target binder, are promising new drug modalities^1^. The E3 binders widely used in PROTACs are CRBN and VHL binders^11,12,13^. The majority of PROTACs in the clinical stage use CRBN binders^73^. Therefore, it is important to evaluate these two types of binders in the same system to increase the versatility of PROTACs. In this study, the two types of binders were compared using the same target binder with the proximity biotinylation enzyme AirID (Fig. 1 ARV-825 and MZ1, Fig. 5 MS154 and MS39). In the case of ARV-825 and MZ1, 31 biotinylated proteins were common. The common biotinylated proteins for ARV-825 and MZ1 included BRD3 and BRD protein-interacting proteins, such as ZNF687, indicating the high proximity specificity of AirID. Furthermore, using MS154 and MS39, seven proteins were found to be common. The common biotinylated proteins for MS154 and MS39 included EGFR and ERRFI1, which interacts with EGFR, confirming the biotinylation of proximity proteins using AirID in various PROTACs. In contrast, 97 and 184 differentially biotinylated proteins were found for ARV-825 and MZ1, respectively. For MS154 and MS39, 28 and 57 proteins differed, respectively. These results suggest that PROTACs with different E3 binders interact with many different proteins, even when the same target binder is used. These results indicate that PROTACs with different E3 ligases induce different interactions that may affect drug action. The data in this study show that the selection of the E3 binder in PROTAC development is important, as the interaction mode of the target protein (Fig. 1 and 5) and its interaction with proteins other than the target protein (Fig. 1, 4, and 5) are greatly affected by different E3 binders on PROTACs. Taken together, these results suggest that analysis of the drug-dependent interactome using proximity biotinylation enzymes, such as AirID, is very important for PROTAC drug discovery and binding agent selection.

We investigated whether the ThBD of CRBN can also be used for the analysis of heterobifunctional molecule-dependent biotinylation with AirID in cells. AirID-fused ThBD induced the biotinylation of 23 proteins in an ARV-825-dependent manner, whereas 130 proteins were biotinylated by AirID-fused full-length CRBN^26^. Even considering that MZ1-dependent proteins biotinylated by the full-length E3 form of VHL were 219, relatively few proteins were biotinylated by ThBD. Proteins biotinylated by ThBD-AGIA-AirID included BRD3, KMT5B, and ZNF687, which interact with BRD4. The number of proteins interacting with BRD4 among the proteins biotinylated by AirID-fused CRBN or ThBD in an ARV-825-dependent manner was 23 out of 130 for CRBN and 5 out of 23 for ThBD. Considering the same ratio of proteins interacting with BRD4 among biotinylated proteins, ThBD could also be used for PROTAC-induced interactome analysis. In addition, we emphasise that the expression levels of full-length CRBNs are lower than those of ThBD. Full-length E3 ligases have auto-ubiquitination capacity. Therefore, it is not easy to overexpress E3 ligases; however, ThBD lacks ubiquitination capacity because it lacks the Lon domain, which contains a binding region for DDB1. Therefore, ThBD is expected to be expressed at higher levels than full-length CRBNs. Additionally, biotinylation of CRL4 and CRL2 complex proteins by ThBD-AGIA-AirID in MM1.S and HCC-827-Luc cells was lower in terms of the number of biotinylation sites compared with that for AirID-CRBN and VHL-AirID (Supplementary Fig. 3c). However, it should be noted that the use of domain proteins is not applicable to all proteins, as VBD-fused AirID did not retain its function in the cell. In summary, these results show that the ligand-binding domains can be used to identify PROTAC target proteins and to analyse more focused PROTAC target-protein interactions.

ARV-110 is undergoing clinical trials; however, there are very few examples of interactome analyses. In this study, AirID-fused ThBD was used for interactome analysis induced by ARV-110 (Fig. 7). The ARV-110-dependent proximity biotinylation interactome showed that many biotinylated proteins, including AR, were localised in the nucleus. These data suggested that the interaction between AR and ARV-110 occurred mainly in the nucleus. This strongly suggests that by using AirID fusion ThBD, the subcellular localisation of interactions between heterobifunctional molecules and target proteins can be analysed. This indicates that the AirID-fused ThBD could be a tool for understanding the interaction mechanism between heterobifunctional molecules and target proteins in cells. In general, the interactome and subcellular localisation of small-molecule compounds bound to target proteins are essential for understanding the activity of drugs, which is also thought to be the case for PROTACs. Studies using FKBP12^F36V^ (dTAG) have shown that the degradation level differs depending on the protein with different localisations, even when the same PROTAC is used^16^. Therefore, the use of AirID-fused ThBD will provide the ability to analyse the target interactome and target localisation of PROTACs used in clinical trials and is expected to be very useful for future PROTAC development.

Currently, heterobifunctional molecules extend beyond proteolysis^74^. Heterobifunctional molecules other than those involved in proteolytic induction, such as DUBTAC and PhosTAC, are also being developed^55,56,57^. As shown in the present study, AirID can be used to analyse the expected and unexpected interacting proteins of heterobifunctional molecules in detail. In addition, this AirID evaluation system can be analysed using a standard mass spectrometry system; therefore, it can be implemented in various laboratories. In the near future, we hope that the AirID-based evaluation system demonstrated in this study will provide important critical information on biotinylated proteins, leading to the successful generation of heterobifunctional molecules suitable for therapeutic and basic research.

## Methods

### Reagents

Thalidomide (#T2524, Tokyo Chemical Industry), pomalidomide (#P2074, Tokyo Chemical Industry), lenalidomide (#126-06733, FUJIFILM Wako), VH032 (#HY-120217, MedChemExpress), ARV-825 (#HY-16954, MedChemExpress), MZ1 (#S8889, Selleck), MS154 (#7395, R&D Systems, Inc), MS39 (#7397, R&D Systems, Inc), ARV-110 (#S6965, selleck), and MG132 (#3175-v, Peptide Institute Inc., Osaka, Japan) were dissolved in DMSO (#13445–74, Nacalai tesque) at 1–100 mM and stored at −20 °C as stock solutions.

### Plasmids

The pcDNA3.1(+) and pCAGGS vectors were purchased from Invitrogen/Thermo Fisher Scientific and RIKEN, respectively. The pEU vector for wheat cell-free protein synthesis was constructed in our laboratory, as previously described^75^. The pcDNA3.1(+)-AGIA-MCS, pcDNA3.1(+)-HA-MCS, pCAGGS-MCS-FLAG, pCAGGS-HA-MCS, pEU-MCS, pEU-bls-MCS, and pEU-FLAG-GST-MCS plasmids were constructed using polymerase chain reaction (PCR) with the In-Fusion system (Takara Bio) or PCR and restriction enzymes. The pEU-FLAG-HSP90AA1, pEU-FLAG-GST-VHL, -HSPA8, -KMT5B, and -BRD4 plasmids were purchased from the Kazusa DNA Research Institute. ThBD (318–424 aa region of CRBN with C366S mutation)^37^, Venus, and DHFR were the genes used in a previous study^76,77^. The open-reading frames (ORFs) of *CRBN*, *ZNF687*, *RPN1*, *SSR4*, *SLC25A5*, *COPB2*, *MAD2L1BP*, *NPLOC4*, *PARG*, *TIPRL*, *ZNF114*, and *HSPA1B* were purchased from the Mammalian Gene Collection (MGC). The AirID-CRBN, VHL-AirID, AirID-VHL, AGIA-AirID-VBD, VBD-AGIA-AirID, ThBD-AGIA-AirID, and AirID-AGIA-ThBD were amplified, and restriction enzyme sites were added using PCR followed by cloning into pEU-MCS or pCSⅡ-CMV-IRES2-Bsd vector (RIKEN). The ORF of *BRD4* was amplified, restriction enzyme sites were added using PCR, and it was cloned into pEU-bls-MCS or pCAGGS-MCS-FLAG. Restriction enzyme sites were added to IKZF1, SALL4, Venus, DHFR, ZNF687, RPN1, SSR4, SLC25A5, COPB2, MAD2L1BP, NPLOC4, PARG, TIPRL, ZNF114, HSPA1B, and KMT5B using PCR. IKZF1, SALL4, DCAF13, and HSPA1B were cloned into the pcDNA3.1(+)-AGIA-MCS. DHFR, ZNF687, RPN1, SSR4, and HSPA1B were cloned into pEU-FLAG-GST-MCS. SLC25A5, COPB2, MAD2L1BP, NPLOC4, PARG, TIPRL, and ZNF114 were cloned into the pEU-MCS-FLAG vector. Venus was cloned into pcDNA3.1(+)-HA-MCS. KMT5B was cloned into the pCAGGS-HA-MCS vector.

### Cell culture and transfection

HEK293T (#CRL-3216, American Type Culture Collection [ATCC], RRID:CVCL_0063) cells were cultured in low-glucose DMEM (DMEM, #041-29775; FUJIFILM Wako) supplemented with 10% foetal bovine serum (#535-94155, FUJIFILM Wako), 100 U/mL penicillin, and 100 µg/mL streptomycin(#15140122, Thermo Fisher Scientific, MA, USA) at 37 °C under 5% CO_2_. HEK293T cells were transiently transfected with TransIT-LT1 transfection reagent (#MIR2304, Mirus Bio) or polyethyleneimine (PEI) Max (MW 40,000) (#2476, PolyScience, Inc.).

MM1.S cells (#CRL-2974, ATCC, RRID:CVCL_8792) were cultured in RPMI160 GlutaMAX medium (#72400047, Gibco) supplemented with 10% foetal bovine serum (#535-94155, FUJIFILM Wako), 100 U/mL penicillin, 100 µg/mL streptomycin, and 55 µM 2-mercaptoethanol (#21985023, Thermo Fisher Scientific) at 37 °C under 5% CO_2_.

B16F10 (RIKEN BRC, RRID:CVCL 0159), HCC-827-Luc (JCRB Cell Bank, RRID:CVCL_4W94), and LNCaP (RIKEN BRC, RRID:CVCL_0395) cells were cultured in RPMI160 GlutaMAX medium (#72400047, Gibco) supplemented with 10% foetal bovine serum (#535-94155, FUJIFILM Wako), 100 U/mL penicillin, 100 µg/mL streptomycin, at 37 °C under 5% CO_2_.

### Generation of stable cell lines using lentivirus

Each lentivirus was produced in HEK293T cells by transfection with the pCSII-CMV-ORF-IRES2-Bsd expression vector together with pCMV-VSV-G-RSV-Rev and pCAG-HIVgp as described in previous study^26^. HEK293T, HCC827-Luc, and LNCaP cells supplemented with 10 µg/mL polybrene (#12996-81, Nacalai Tesque) were infected with the appropriate lentivirus. In the case of MM1.S, the cells were suspended in 50 µL culture medium containing 8 µg/mL polybrene and lentivirus and infected by incubation at 37 °C for 30 min. After 24 h of infection, the culture medium was replaced, and 10 µg/mL blasticidin S (#ant-bl-10p, InvivoGen) selection was started 24 h thereafter.

### Antibodies

The following horseradish peroxidase (HRP)-conjugated antibodies were used in this study: FLAG (Sigma-Aldrich, #A8592, 1:5000), AGIA (produced in our laboratory, 1:10000)^36^, α-tubulin (MBL, #PM054-7,1:5000), biotin (Cell Signaling Technology, #7075,1:1000), and anti-GAPDH (MBL, #M171-7, 1:5000). The following primary antibodies were used in this study: Anti-BRD4 (Cell Signaling Technology, #13440, 1:1000), anti-BirA (Agrisera, #AS20 4440, 1:10000), anti-VHL (Cell Signaling Technology, #68547, 1:1000), anti-SALL4 (Santa Cruz Biotechnology, #sc-101147, 1:500), anti-IKZF1 (Cell Signaling Technology, #14859, 1:1000), anti-BRD3 (Santa Cruz Biotechnology, #sc-81202, 1:500), anti-EGFR (produced in our laboratory, EmAb-134, 1:1000)^78,79^, anti-HSP90 (Cell Signaling Technology, #4874, 1:1000), and anti-AR (Cell Signaling Technology, #5153, 1:1000). Anti-rabbit IgG (HRP-conjugated, Cell Signaling Technology, #7074, 1:10000), anti-mouse IgG (HRP-conjugated, Cell Signaling Technology, #7076, 1:10000).

### Immunoblot analysis

Each cell pellet was lysed in RIPA buffer (25 mM Tris-HCl [pH 8.0], 150 mM NaCl, 1% NP-40, 0.5% sodium deoxycholate, 0.1% sodium dodecyl sulphate (SDS), and 1 mM EDTA) containing a protease inhibitor cocktail (#P8340, Sigma-Aldrich). The cell lysates were centrifuged at 16,100 × *g* for 15 min, and the protein concentration in the supernatant was quantified using a BCA assay kit (#23227, Thermo Fisher Scientific). Protein samples were separated using SDS-PAGE and transferred onto polyvinylidene difluoride (PVDF) membranes (#IPVH00010, Millipore). The membranes were blocked using 5% skim milk (#4273437, Megmilk Snow Brand) in TBST (20 mM Tris-HCl [pH 7.5], 150 mM NaCl, 0.05% Tween20) at 27 °C for 1 h, and then treated with the appropriate antibodies. Immobilon (#WBKLS0500, Millipore), ImmunoStar LD (#290-69904, FUJIFILM Wako), or EzWestLumi plus (#2332638, Atto) were used as substrates for HRP, and the luminescence signal was detected using an ImageQuant LAS 4000 mini (GE Healthcare, version 1.1). For some blots, the membrane was stripped with a stripping solution (#193-16375, FUJIFILM Wako) and reprobed with other antibodies.

### Wheat cell-free protein synthesis

The recombinant proteins were synthesised using a wheat cell-free system^75^. In vitro transcription and wheat cell-free protein synthesis were performed using the WEPRO7240 expression kit (Cell-Free Sciences). Transcription was performed using SP6 RNA polymerase, with plasmids or DNA fragments as templates.

### In vitro PROTAC-dependent biotinylation of POI and streptavidin pull-down assay

The AirID-fused and FG-fused proteins synthesised using the wheat cell-free protein synthesis system were each mixed in a volume of 20 µL. For the biotinylation assay of SALL4 dependent on pomalidomide by AirID-fused CRBN or −ThBD, DMSO, or 2 µM pomalidomide were mixed in the presence of 500 nM biotin and 100 mM NaCl at a final volume of 50 µL and incubated at 27 °C for 3 h. For the biotinylation assay of BRD4 dependent on MZ1 by AirID-fused VHL or −VBD, DMSO, 1 µM VH032, or 1 µM MZ1 were mixed in the presence of 500 nM biotin and 100 mM NaCl at a final volume of 50 µL and incubated at 27 °C for 3 h. For biotinylated samples, a tenfold volume of 1% SDS buffer (50 mM Tris-HCl pH 7.5, 150 mM, 1% SDS) and 20 µL of STA-Sepharose suspended in 200 µL Triton lysis buffer were added. After 2 h of rotation at room temperature and three washes with 500 µL of Triton lysis buffer, biotinylated proteins were eluted by boiling with 50 µL of 2× sample buffer containing 5% 2-mercaptoethanol.

### AlphaScreen-based biochemical assays using in vitro PROTACs dependent biotinylation samples

The AirID-fused and FG-fused proteins synthesised using the wheat cell-free protein synthesis system were each mixed in a volume of 20 µL. Then, DMSO or pomalidomide (final 1-20 µM), 1 µM ARV-825, 1 µM MS154, 1 µM MS39 were mixed in the presence of 500 nM biotin and 100 mM NaCl in a final volume of 50 µL and incubated at 27 °C for 3 h. Biotinylated mixtures (1 µL) were prepared in 20 µL AlphaScreen buffer containing 100 mM Tris (pH 8.0), 0.1% Tween20, 100 mM NaCl, and 1 mg/mL BSA. Subsequently, the mixtures were added to a 384-well AlphaPlate (PerkinElmer). Then, 5 µL detection mixture containing 0.2 µg/mL anti-DYKDDDDK mouse mAb (#012-22384, FUJIFILM Wako), 0.08 µL streptavidin-coated donor beads, and 0.08 µL Protein A-coated acceptor beads in AlphaScreen buffer were added to each well. After incubation at 27 °C for 1 h, luminescence signals were detected using an EnVision plate reader (PerkinElmer version 1.12).

### Analysis of E3, PROTAC, and POI ternary complex interaction using the AlphaScreen method

Biotinylated GST-bls-ThBD mixtures (1 µL) were prepared in 10 µL AlphaScreen buffer containing 100 mM Tris (pH 8.0), 0.1% Tween20, 100 mM NaCl, and 1 mg/mL BSA. PROTAC mixtures at the indicated concentrations were prepared in 5 µL AlphaScreen buffer. Substrate mixtures (5 µL) containing 1 µL FLAG-HSP90AA1 in AlphaScreen buffer were prepared. Subsequently, the three solutions were mixed and incubated at 27 °C for 1 h in a 384-well AlphaPlate (PerkinElmer). Thereafter, 5 µL detection mixture containing 0.2 µg/mL anti-DYKDDDDK mouse mAb (#012-22384, FUJIFILM Wako), 0.08 µL streptavidin-coated donor beads, and 0.08 µL Protein A-coated acceptor beads (µL) in AlphaScreen buffer were added to each well. After incubation at 27 °C for 1 h, luminescence signals were detected using an EnVision plate reader (PerkinElmer version 1.12).

### Cell biotinylation assay with AirID-fused proteins

HEK293T cells stably expressing AirID-AGIA-ThBD or ThBD-AGIA-AirID were cultured in 6-well plates and transfected with 500 ng pcDNA3.1(+)-AGIA-IKZF1 and 500 ng pcDNA3.1(+)-AGIA-SALL4 together with 1 µg pcDNA3.1(+). After 24 h of transfection, the cells were treated with 10 µM pomalidomide or DMSO in the presence of 10 µM biotin and 5 µM MG132 or DMSO for 6 h. The cells were harvested in PBS. The cell pellets were washed with 1× PBS and lysed in 300 µL 2% SDS lysis buffer (50 mM Tris-HCl pH 7.5, 2% SDS) containing a protease inhibitor cocktail (Sigma-Aldrich), and the lysates were denatured by boiling at 98 °C for 15 min. The lysates were then diluted with 300 µL of 50 mM Tris-HCl (pH 7.5), and clarified by centrifugation at 16,100 × *g* for 15 min. Thereafter, 560 µL of lysate was added to 500 µL RIPA lysis buffer containing 30 µL streptavidin Sepharose High Performance (#90100484, Cytiva). After rotating at 4 °C overnight, the beads were washed three times with 500 µL 1% SDS buffer (50 mM Tris-HCl pH 7.5, 1% SDS), and the proteins were eluted by boiling with 50 µL of 1× sample buffer containing 5% 2-mercaptoethanol.

### Streptavidin pull-down assay of PROTACs in cells

For the STA-PDA using ARV-825 or MZ1, HEK293T, MM1.S, and B16F10 cells stably expressing AirID-CRBN, ThBD-AGIA-AirID, AirID-AGIA-ThBD, VHL-AirID, AirID-VHL, or VBD-AGIA-AirID were cultured in 10 cm dishes and treated with DMSO, 1 µM pomalidomide, 3 µM MZ1, 1 µM ARV-825, or GO-HSP heterobifunctional molecules in the presence of 10 µM biotin and 5 µM MG132 for 6 h.

For the assay using MS154, HCC-827-Luc cells stably expressing ThBD-AGIA-AirID were cultured in 10 cm dishes. Next, the medium was changed to serum-free RPMI1640 and treated with 5 µM MG132 for 1 h. Subsequently, pomalidomide (final 1 µM) or MS154 (final 1 µM) was added together with biotin (final 10 µM) and the mixture was incubated for 6 h.

For the assay using MS39, HCC-827-Luc cells stably expressing VBD-AGIA-AirID or VHL-AirID were cultured in 10 cm dishes. Next, cells were treated with 5 µM MG132 and incubated for 1 h, after which VH032 (final 3 µM) or MS39 (final 3 µM) was added with biotin (final 10 µM) and incubated for 6 h.

For the assay using ARV-110, LNCaP cells stably expressing ThBD-AGIA-AirID were cultured in 10 cm dishes. Next, cells were treated with 5 µM MG132 and incubated for 1 h, after which pomalidomide (final 1µM) or ARV-110 (final 1µM) was added with biotin (final 10 µM) and incubated for 6 h.

The cells were harvested by suspending in 1 mL PBS. The cell pellets were washed with 1× PBS and lysed in 400 µL SDS lysis buffer (50 mM Tris-HCl pH 7.5, 150 mM NaCl, 2% SDS) containing a protease inhibitor cocktail (Sigma-Aldrich), and the lysates were denatured by boiling at 98 °C for 15 min. Then, the lysate was sonicated (ON 60 s; OFF 30 s) and clarified by centrifugation at 16,100 × *g* for 15 min and diluted 2-fold with 400 µL of 50 mM Tris-HCl (pH 7.5). Thereafter, 760 µL of lysate was added to 400 µL Triton lysis buffer (150 mM NaCl, 25 mM Tris-HCl pH7.5, 1mM EDTA, 1% Triton-X 100) containing 20 µL streptavidin Sepharose beads or streptavidin Dynabeads™ MyOne™ Streptavidin C1 (#DB65001, VERITAS). After rotating overnight at 4 °C, the beads were washed three times with 500 µL Triton lysis buffer, and the proteins were eluted by boiling with 50 µL of 2× sample buffer containing 5% 2-mercaptoethanol.

### CellTiter-Glo assay in HEK293T

HEK293T cells were seeded at 2 × 10^5^ cells/well in 96-well plates. The day after seeding, the drug was added at the indicated concentration, and the cells were incubated for 24 h. Living cells were counted using the CellTiter-Glo assay kit (#G7571, Promega).

### Preparation of cell lysates treated with PROTACs for enrichment of biotinylated peptides

Cells were cultured independently in three dishes, and biotinylation reactions were performed in each dish. After washing three times with 10 mL of HEPES saline buffer (20 mM HEPES-NaOH, pH 7.5, 137 mM NaCl), the cells were lysed in 300 µL (MM1.S cells, HEK293T cells, and HCC827-luc cells) or 500 µL (LNCaP cells) Gdm-TCEP buffer (6 M guanidine-HCl, 100 mM HEPES-NaOH [pH 7.5], 10 mM TCEP, 40 mM chloroacetamide). Cell lysates were pooled in one tube and divided into three tubes of 300 or 500 µL each.

### Enrichment of biotinylated peptides using Tamavidin 2-REV

All experiments were performed in triplicates for each treatment. The cell lysates in Gdm-TCEP buffer were dissolved by heating and sonication and then centrifuged at 20,000 × *g* for 15 min at 4 °C. The supernatants were recovered, and proteins were purified by methanol–chloroform precipitation and solubilised in 150 µL of PTS buffer (12 mM SDC, 12 mM SLS, 100 mM Tris-HCl, pH8.0). After sonication and heating, the protein solution was diluted 5-fold with 100 mM Tris-HCl, pH8.0 and digested overnight with trypsin (MS grade, Thermo Fisher Scientific) at 37 °C. The resulting peptide solutions were diluted 2-fold with TBS (50 mM Tris-HCl, pH 7.5, 150 mM NaCl). Biotinylated peptides were captured on a 15 µL slurry of MagCapture HP Tamavidin 2-REV magnetic beads (#133-18611, FUJIFILM Wako) by incubation for 3 h at 4°C. After washing with TBS five times, the biotinylated peptides were eluted twice with 100 µL of 1 mM biotin in TBS for 15 min at 37 °C. The combined eluates were desalted using GL-Tip SDB (#7820-11200, GL Sciences), evaporated in a SpeedVac concentrator (Thermo Fisher Scientific), and re-dissolved in 0.1% TFA and 3% acetonitrile (ACN).

### Data-dependent LC-MS/MS analysis

LC-MS/MS analysis of the biotinylated peptides was performed using an EASY-nLC 1200 UHPLC system connected to an Orbitrap Fusion mass spectrometer through a nanoelectrospray ion source (Thermo Fisher Scientific). The peptides were separated on a 150 mm C18 reversed-phase column with an inner diameter of 75 µm (Nikkyo Technos) with a linear 4–32% ACN gradient for 0–60 min, followed by an increase to 80% ACN for 10 min and a final hold at 80% ACN for 10 min. The mass spectrometer was operated in data-dependent acquisition mode with a maximum duty cycle of 3 s. The MS1 spectra were measured at a resolution of 120,000, an automatic gain control (AGC) target of 4 × 10^5^, and a mass range of 375–1500 *m/z*. HCD MS/MS spectra were acquired using a linear ion trap with an AGC target of 1 × 10^4^, an isolation window of 1.6 *m/z*, a maximum injection time of 200 ms, and a normalised collision energy of 30. Dynamic exclusion was set to 10 s. Raw data were directly analysed against the Swiss-Prot database restricted to *Homo sapiens* using Proteome Discoverer version 2.5 (Thermo Fisher Scientific) with the Sequest HT search engine. The search parameters were as follows: (a) trypsin as an enzyme with up to two missed cleavages, (b) precursor mass tolerance of 10 ppm, (c) fragment mass tolerance of 0.6 Da; (d) carbamidomethylation of cysteine as a fixed modification, and (e) acetylation of protein N-terminus, oxidation of methionine, and biotinylation of lysine as variable modifications. Peptides were filtered at a false discovery rate (FDR) of 1% using a percolator node. Label-free quantification was performed based on the intensities of the precursor ions using a precursor-ion quantifier node. Normalisation was performed such that the sum of the abundance values for each sample was the same for all peptides. For statistical analyses of the MS data, the *P*-values in each volcano plot were calculated using Student’s *t*-test in Microsoft Excel (version 16.78.3). The adjusted *P*-values were calculated by controlling the FDR using Microsoft Excel (version 16.78.3) and are shown in Supplementary Data.

### Immunoprecipitation

HEK293T cells were cultured in 10 cm dishes and transfected with 10 µg pcDNA3.1(+)-AGIA-Venus, DCAF13, or HSPA1B. After incubation for 24 h, the cells were lysed in 1 mL of Triton Lysis buffer containing a protease inhibitor cocktail (#P8340, Sigma-Aldrich) and Benzonase Nuclease (#E1014, Sigma-Aldrich). The lysate was rotated at 4 °C for 1 h, then centrifuged at 16,100 × *g* for 5 min. The supernatant was collected and mixed with 1 µg of anti-AGIA antibody. After 30 min rotation at 4 °C, 12 µg Dynabeads Protein G (#DB10004, VERITAS) suspended in 50 µL Triton Lysis buffer was added. After 2 h rotation at 4 °C, the samples were washed two times with 500 µL PBS and one time with 500 µL Triton Lysis buffer, and the proteins were eluted by boiling with 50 µL of 2× sample buffer containing 5% 2-mercaptoethanol.

HEK293T cells were cultured in 10 cm dishes and transfected with 5 µg pcDNA3.1(+)-HA-Venus or pCAGGS-HA-KMT5B together with 5 µg pCAGGS-BRD4-FLAG. After incubation for 24 h, the medium was replaced with fresh medium. After 48 h of transfection, the cells were lysed in 400 µL of NP-40 Lysis buffer (20 mM HEPES NaOH pH7.5, 400 mM NaCl, 1 mM EDTA, 1.5 mM MgCl_2_, 0.5% NP-40) containing protease inhibitor cocktail (#P8340, Sigma-Aldrich) and Benzonase Nuclease (#E1014, Sigma-Aldrich). The lysate was rotated at 4 °C for 1 h, and then centrifuged at 16,100 × *g* for 5 min. The supernatant was collected and mixed with 1 µg of anti-DYKDDDK antibody. After 30 min rotation at 4 °C, 15 µg Dynabeads Protein A (DB100008, VERITAS) suspended in 100 µL NP-40 Lysis buffer was added to the sample. After 3 h rotation at 4 °C, the samples were washed three times with 500 µL NP-40 Lysis buffer, and the proteins were eluted by boiling with 40 µL of 2× sample buffer containing 5% 2-mercaptoethanol.

### Gene ontology and pathway analyses

GO analysis was performed using ShinyGO 0.77 (http://bioinformatics.sdstate.edu/go77/). Pathway diagrams of protein–protein interaction networks based on data analysed with STRING (https://string-db.org) were generated using the Cytoscape software (https://cytoscape.org).

### Chemical synthesis of geldanamycin-thalidomide derivatives

The synthesis methods of geldanamycin-thalidomide derivatives and their intermediates are provided in the Supplementary Information (**General information for chemical synthesis**).

### Statistical analyses

Significant changes were analysed by Student’s *t*-test using Microsoft Excel spreadsheets with a basic statistical program, two-way ANOVA with Bonferroni’s multiple comparison test or one-way ANOVA followed by Tukey’s post-hoc test using the GraphPad Prism 9 software (GraphPad, Inc.). For all tests, *P* < 0.05 was considered statistically significant. Immunoblot analyses and streptavidin pull-down assays were repeated more than twice, with similar results.

## Acknowledgments

We would like to thank the Applied Protein Research Laboratory of Ehime University. We thank Prof. Y. Imai for providing us with the LNCaP cell line. This work was supported by a grant from the Platform Project for Supporting Drug Discovery and Life Science Research (Basis for Supporting Innovative Drug Discovery and Life Science Research [BINDS]) from AMED under Grant Numbers JP24ama121010J0003 (T.S.) and JP24ama121040J0003 (Y.I.); KAKENHI (21K19230 and 24H00560 to T.S.) from the Japan Society for the Promotion of Science (JSPS); a Grant-in-Aid for JSPS Fellows (24KJ1755 for K.Y.) from JSPS; and Joint Usage and Joint Research Programs of the Institute of Advanced Medical Sciences of Tokushima University. We would like to thank Editage (www.editage.com) for English language editing.

## Author contributions

K.Y. performed cloning and characterisation of AirID-fused proteins, in vitro assays, cell-based assays, GO analysis, and biotinylation assay; S.Y established and characterised cell lines stably expressing AirID-CRBN; H.K. performed enrichment of biotinylated peptides and LC-MS/MS analyses; H.Y. and Y. I performed chemical synthesis of GA-TDs; K.Y. and T.S. analysed the data and wrote the draft paper; T.S. conceived the research and designed the study; K.Y., H.K., and T.S. designed the experiments, wrote the paper, and all authors revised the manuscript.

## Corresponding authors

Correspondence and requests for materials should be addressed to Tatsuya Sawasaki or Hidetaka Kosako.

