## Supplementary Information for "In-cell proximity target validation methods for heterobifunctional molecules with CRBN- or VHL-binder using AirID"

*[1]Division of Cell-Free Life Science, Proteo-Science Center, Ehime University, 3 Bunkyo-cho, Matsuyama, Ehime 790-8577, Japan. [2]Division of Proteo-Interactome, Proteo-Science Center, Ehime University, 3 Bunkyo-cho, Matsuyama, Ehime 790-8577, Japan. [3]Graduate School of Pharmaceutical Sciences, Tohoku University, 6-3, Aoba, Aramaki, Aoba-ku, Sendai 980-8578, Japan. [4]Division of Cell Signaling, Institute of Advanced Medical Sciences, Tokushima University, Tokushima 770-8503, Japan.*

*Corresponding Authors:

Tatsuya Sawasaki

Proteo-Science Center, Ehime University, Matsuyama 790-8577, Japan

Hidetaka Kosako

Institute of Advanced Medical Sciences, Tokushima University, Tokushima 770-8503, Japan

**

**a**

Proteins biotinylated by VHL-AirID

| Protein name | Uniprot code |
|---|---|
| AATF | Q9NY61 |
| ACTB | P60709 |
| ACTR2 | P61160 |
| ADAM8 | P78325-2 |
| AFG3L2 | Q9Y4W6 |
| AHSA1 | O95433 |
| AKNA | Q7Z591 |
| ALDOA | P04075 |
| ANKHD1 | Q8IWZ3 |
| ARHGAP21 | Q5T5U3 |
| ARID5A | Q03989 |
| ARMC6 | Q6NXE6 |
| ASNS | P08243 |
| ATAD5 | Q96QE3 |
| ATF1 | P18846 |
| ATP13A1 | Q9HD20 |
| ATRIP | Q8WXE1 |
| ATXN2L | Q8WWM7 |
| AVEN | Q9NQS1 |
| B2M | P61769 |
| BCOR | Q6W2J9 |
| BRD2 | P25440 |
| BRD3 | Q15059 |
| BRD4 | O60885 |
| BSG | P35613 |
| CAMKK2 | Q96RR4 |
| CANX | P27824 |
| CBX4 | O00257 |
| CCAR2 | Q8N163 |
| CCNI | Q14094 |
| CCNT2 | O60583 |
| CCPG1 | Q9ULG6 |
| CCT3 | P49368 |
| CCT6A | P40227 |
| CCT8 | P50990 |
| CD2AP | Q9Y5K6 |
| CD74 | P04233 |
| CDK1 | P06493 |
| CDKAL1 | Q5VV42 |
| CEP192 | Q8TEP8 |
| CEP350 | Q5VT06 |
| CHAMP1 | Q96JM3 |
| CLGN | O14967 |
| CLOCK | O15516 |
| CMTR2 | Q8IYT2 |
| COPG1 | Q9Y678 |
| CORO7 | P57737 |
| CRYBB1 | P53674 |
| CSNK1A1 | P48729 |
| CTR9 | Q6PD62 |
| CYB5R1 | Q9UHQ9 |
| CYP51A1 | Q16850 |
| DDHD1 | Q8NEL9 |
| DDOST | P39656 |
| DDX3X | O00571 |
| DDX5 | P17844 |
| DIDO1 | Q9BTC0 |
| DNAJC2 | Q99543 |
| DOCK4 | Q8N1I0 |
| DPYD | Q12882 |
| DUSP5 | Q16690 |
| DUSP6 | Q16828 |
| EEF2 | P13639 |
| EIF2B5 | Q13144 |
| EIF2S3 | P41091 |
| EIF3F | O00303 |
| ELF2 | Q15723-1 |
| ENO1 | P06733 |
| EPRS1 | P07814 |
| ERGIC3 | Q9Y282 |
| ERLEC1 | Q96DZ1 |
| ERN1 | O75460 |
| ESS2 | Q96DF8 |
| EXOC4 | Q96A65 |

| Protein name | Uniprot code |
|---|---|
| FBL | P22087 |
| FDFT1 | P37268 |
| FNDC3B | Q53EP0 |
| G3BP1 | Q13283 |
| GMPPA | Q96IJ6 |
| GNAS | Q5JWF2 |
| GNPTG | Q9UJJ9 |
| GTF2F1 | P35269 |
| HAX1 | O00165 |
| HCFC1 | P51610 |
| HDAC9 | Q9UKV0 |
| HERC4 | Q5GLZ8 |
| HLA-B | P01889 |
| HLA-E | P13747 |
| HNRNPA0 | Q13151 |
| HP1BP3 | Q5SSJ5 |
| HSP90AA1 | P07900 |
| HSP90AB1 | P08238 |
| HSP90B1 | P14625 |
| HSPA1B | P0DMV9 |
| HSPA4L | O95757 |
| HSPA5 | P11021 |
| HSPA6 | P17066 |
| HSPA8 | P11142 |
| HSPB1 | P04792 |
| HSPH1 | Q92598 |
| IARS1 | P41252 |
| IDH1 | O75874 |
| IGF2BP2 | Q9Y6M1 |
| IGLL5 | B9A064 |
| INAVA | Q3KP66 |
| INO80 | Q9ULG1 |
| IRS2 | Q9Y4H2 |
| ITGB7 | P26010 |
| ITM2C | Q9NQX7 |
| JCHAIN | P01591 |
| JMJD1C | Q15652 |
| KDM4B | O94953 |
| KIAA0232 | Q92628 |
| KMT2A | Q03164 |
| KMT5B | Q4FZB7 |
| LAMA5 | O15230 |
| LIN54 | Q6MZP7 |
| MACF1 | Q9UPN3 |
| MAGED1 | Q9Y5V3 |
| MARF1 | Q9Y4F3 |
| MARS1 | P56192 |
| MDH2 | P40926 |
| MED1 | Q15648 |
| MMGT1 | Q8N4V1 |
| MTHFD2 | P13995 |
| MVP | Q14764 |
| MYL6 | P60660 |
| NBR1 | Q14596 |
| NFATC2IP | Q8NCF5 |
| NFRKB | Q6P4R8 |
| NIPBL | Q6KC79 |
| NLRP7 | Q8WX94 |
| NMT1 | P30419 |
| NOMO2 | Q5JPE7 |
| NONO | Q15233 |
| NSD3 | Q9BZ95 |
| NUP88 | Q99567 |
| OLAH | Q9NV23 |
| P4HA1 | P13674 |
| PABPC1 | P11940 |
| PARP4 | Q9UKK3 |
| PBXIP1 | Q96AQ6 |
| PCBP2 | Q15366 |
| PCNT | O95613 |
| PFN1 | P07737 |
| PHF8 | Q9UPP1 |
| PI4KA | P42356 |
| PLOD3 | O60568 |

| Protein name | Uniprot code |
|---|---|
| POLD1 | P28340 |
| POLR2A | P24928 |
| PPIB | P23284 |
| PPIP5K2 | O43314 |
| PPP2R5D | Q14738 |
| PPP6C | O00743 |
| PRKDC | P78527 |
| PSAP | P07602 |
| PSAT1 | Q9Y617 |
| PSMB10 | P40306 |
| PSMB3 | P49720 |
| PSMD2 | Q13200 |
| PSMD3 | O43242 |
| PTBP3 | O95758-1 |
| PTPN6 | P29350 |
| PYGB | P11216 |
| RANBP2 | P49792 |
| RBBP5 | Q15291 |
| RBM39 | Q14498 |
| REPS1 | Q96D71 |
| RESF1 | Q9HCM1 |
| RPL10 | P27635 |
| RPL13A | P40429 |
| RPL8 | P62917 |
| RPN1 | P04843 |
| RPS21 | P63220 |
| RRAGC | Q9HB90 |
| SCRN3 | Q0VDG4 |
| SDAD1 | Q9NVU7 |
| SEC11B | P0C7V7 |
| SEC11C | Q9BY50 |
| SEC24D | O94855 |
| SLFN13 | Q68D06 |
| SMAP | O00193 |
| SMC1A | Q14683 |
| SMC4 | Q9NTJ3 |
| SMC5 | Q8IY18 |
| SPATA2 | Q9UM82 |
| SPATS2 | Q86XZ4 |
| SPCS2 | Q15005 |
| SREBF2 | Q12772 |
| SSR2 | P43308 |
| SSR4 | P51571 |
| STAT2 | P52630 |
| TAF1 | P21675 |
| TAF3 | Q5VWG9 |
| TAPBP | O15533 |
| TARS2 | Q9BW92 |
| TBL3 | Q12788 |
| TCIRG1 | Q13488 |
| TDG | Q13569 |
| TFIP11 | Q9UBB9 |
| TIPRL | O75663 |
| TMBIM6 | P55061 |
| TMEM199 | Q8N511 |
| TMEM248 | Q9NWD8 |
| TMEM9 | Q9P0T7 |
| TMPO | P42167 |
| TNRC6B | Q9UPQ9 |
| TONSL | Q96HA7 |
| TRAPPC9 | Q96Q05 |
| TRIM4 | Q9C037 |
| TRMT112 | Q9UI30 |
| TXNDC5 | Q8NBS9 |
| UBIAD1 | Q9Y5Z9 |
| VPS11 | Q9H270 |
| VPS26C | O14972 |
| WWOX | Q9NZC7 |
| ZNF518A | Q6AHZ1 |
| ZNF687 | Q8N1G0 |
| ZNF841 | Q6ZN19-3 |

**b**

Proteins biotinylated by AirID-CRBN

| Protein name | Uniprot code |
|---|---|
| ADTRP | Q96IZ2 |
| AHR | P35869 |
| ANKRD12 | Q6UB98 |
| ARID5A | Q03989 |
| ATF6 | P18850 |
| ATF6B | Q99941 |
| ATG3 | Q9NT62 |
| ATP1A1 | P05023 |
| B2M | P61769 |
| BCOR | Q6W2J9 |
| BRD2 | P25440 |
| BRD3 | Q15059 |
| BRD4 | O60885 |
| C12orf45 | Q8N5I9 |
| C19orf47 | Q8N9M1 |
| CASP10 | Q92851 |
| CASP3 | P42574 |
| CCNT1 | O60563 |
| CCNT2 | O60583 |
| CDK1 | P06493 |
| CDK2 | P24941 |
| CDK9 | P50750 |
| CENPC | Q03188 |
| CEP350 | Q5VT06 |
| CHD1L | Q86WJ1 |
| CORO1C | Q9ULV4 |
| CRBN | Q96SW2 |
| CTR9 | Q6PD62 |
| DEPTOR | Q8TB45 |
| DIDO1 | Q9BTC0 |
| DUSP5 | Q16690 |
| ELAC2 | Q9BQ52 |
| ELF2 | Q15723 |
| FTL | P02792 |
| GNAS | P63092 |
| HDAC1 | Q13547 |
| HERPUD1 | Q15011 |
| HMGCR | P04035 |
| HNRNPL | P14866 |
| HSP90AB1 | P08238 |
| HSPA5 | P11021 |
| HSPA8 | P11142 |
| HSPD1 | P10809 |
| HSPH1 | Q92598 |
| INVS | Q9Y283 |
| ITGB7 | P26010 |
| ITM2C | Q9NQX7 |
| JMJD1C | Q15652 |
| JSRP1 | Q96MG2 |
| KDM3A | Q9Y4C1 |
| KDM4B | O94953 |
| KMT5B | Q4FZB7 |
| LATS2 | Q9NRM7 |
| LSS | P48449 |
| MAGED1 | Q9Y5V3 |
| MARF1 | Q9Y4F3 |
| MBD4 | O95243 |
| MFAP1 | P55081 |
| MGA | Q8IWI9 |
| MIS18BP1 | Q6P0N0 |
| MKI67 | P46013 |
| MOB3B | Q86TA1 |
| MSL2 | Q9HCI7 |
| MXD1 | Q05195 |
| MYOM2 | P54296 |
| N4BP1 | O75113 |
| NABP2 | Q9BQ15 |
| NEB | P20929 |
| NECAP2 | Q9NVZ3 |
| NELFA | Q9H3P2 |
| NFIL3 | Q16649 |
| NFKB2 | Q00653 |
| NKRF | O15226 |
| NOMO2 | Q5JPE7 |
| NPAT | Q14207 |
| NSD3 | Q9BZ95 |

| Protein name | Uniprot code |
|---|---|
| PAF1 | Q8N7H5 |
| PHRF1 | Q9P1Y6 |
| PLOD3 | O60568 |
| POLR2A | P24928 |
| PPIL2 | Q13356 |
| PRDM1 | O75626 |
| PROSER1 | Q86XN7 |
| PSMB7 | Q99436 |
| PTPN2 | P17706 |
| RFX5 | P48382 |
| RLF | Q13129 |
| RNASEH1 | O60930 |
| RPL10 | P27635 |
| RPL13A | P40429 |
| RPL18A | Q02543 |
| RPL28 | P46779 |
| RPL4 | P36578 |
| RPN2 | P04844 |
| RPS11 | P62280 |
| RPS18 | P62269 |
| RPS24 | P62847 |
| RSL24D1 | Q9UHA3 |
| RTEL1 | Q9NZ71 |
| SEPTIN1 | Q8WYJ6 |
| SERBP1 | Q8NC51 |
| SETX | Q7Z333 |
| SH2D3C | Q8N5H7 |
| SLBP | Q14493 |
| SMARCC1 | Q92922 |
| SMC5 | Q8IY18 |
| SND1 | Q7KZF4 |
| SNRK | Q9NRH2 |
| SPEN | Q96T58 |
| STAT2 | P52630 |
| SULF1 | Q8IWU6 |
| TCIRG1 | Q13488 |
| TENT5C | Q5VWP2 |
| TFCP2 | Q12800 |
| TIPARP | Q7Z3E1 |
| TMCO1 | Q9UM00 |
| TMPO | P42166 |
| TRIM37 | O94972 |
| TSHZ1 | Q6ZSZ6 |
| TUFT1 | Q9NNX1 |
| UIMC1 | Q96RL1 |
| XRCC6 | P12956 |
| YEATS2 | Q9ULM3 |
| ZFAND6 | Q6FIF0 |
| ZHX2 | Q9Y6X8 |
| ZMYND8 | Q9ULU4 |
| ZNF106 | Q9H2Y7 |
| ZNF451 | Q9Y4E5 |
| ZNF592 | Q92610 |
| ZNF687 | Q8N1G0 |

**Supplementary Fig. 1 Proteins biotinylated by AirID-CRBN or VHL-AirID in an ARV-825- or MZ1-dependent manner**

**a** List of MZ1-dependent proteins predominantly biotinylated by VHL-AirID in MM1.S cells (MZ1/VH032 peptides ratio >2, $P < 0.05$). **b** List of ARV-825-dependent proteins predominantly biotinylated by AirID-CRBN in MM1.S cells (ARV-825/pomalidomide peptides ratios >2, $P < 0.05$). ARV-825-dependent biotinylated proteins are listed in the table based on data from Yamanaka et al., *Nat Commun*. (2022)[26].

a b

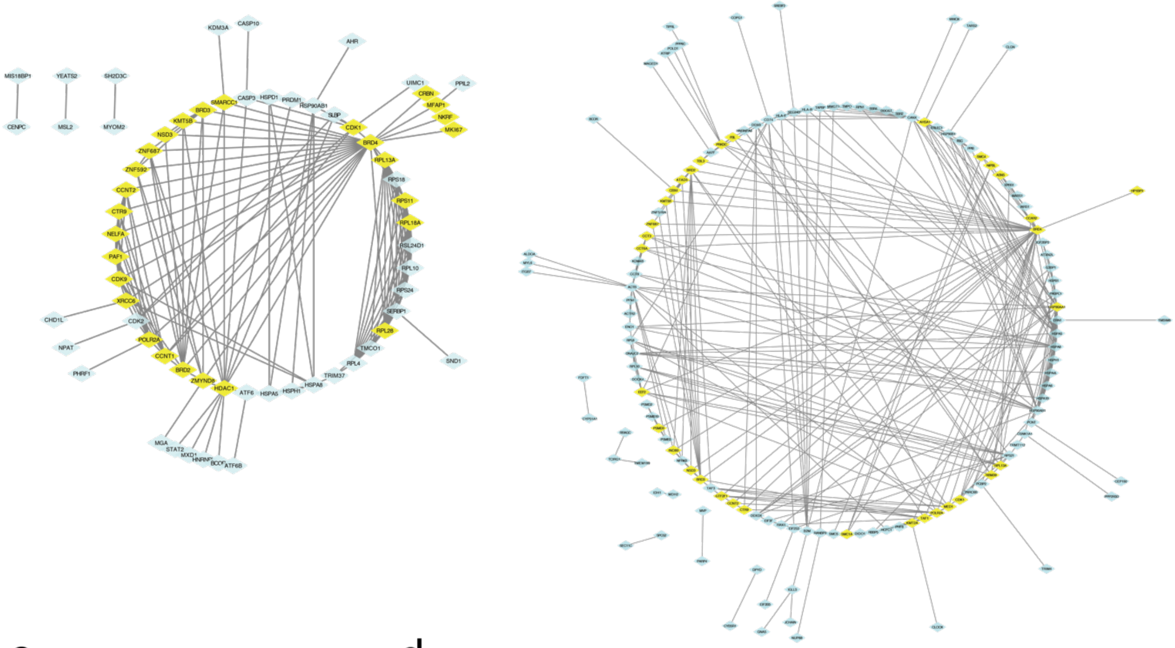

c

Proteins biotinylated by AirID-CRBN and VHL-AirID

| Protein name | Uniprot code |
| --- | --- |
| ARID5A | Q03989 |
| B2M | P61769 |
| BCOR | Q6W2J9 |
| BRD2 | P25440 |
| BRD3 | Q15059 |
| BRD4 | O60885 |
| CCNT2 | O60583 |
| CDK1 | P06493 |
| CEP350 | Q5VT06 |
| CTR9 | Q6PD62 |
| DIDO1 | Q9BTC0 |
| DUSP5 | Q16690 |
| GNAS | Q5JWF2 |
| HSP90AB1 | P08238 |
| HSPA5 | P11021 |
| HSPA8 | P11142 |
| HSPH1 | Q92598 |
| ITGB7 | P26010 |
| ITM2C | Q9NQX7 |
| JMJD1C | Q15652 |
| KDM4B | O94953 |
| KMT5B | Q4FZB7 |
| MAGED1 | Q9Y5V3 |
| MARF1 | Q9Y4F3 |
| NOMO2 | Q5JPE7 |
| NSD3 | Q9BZ95 |
| PLOD3 | O60568 |
| POLR2A | P24928 |
| RPL10 | P27635 |
| RPL13A | P40429 |
| SMC5 | Q8IY18 |
| STAT2 | P52630 |
| TCIRG1 | Q13488 |
| TMPO | P42167 |
| ZNF687 | Q8N1G0 |

d

Proteins biotinylated by AirID-CRBN and VHL-AirID (GO Biological Process)

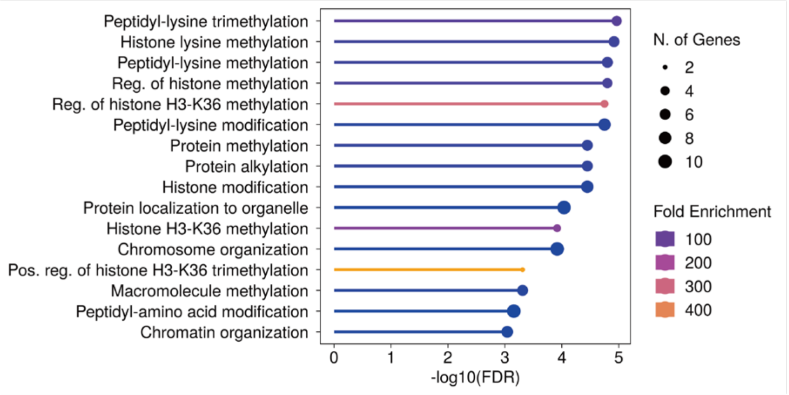


**Supplementary Fig. 2 Pathway and gene ontology analyses of proteins biotinylated by AirID-CRBN or VHL-AirID depending on ARV-825 or MZ1**

**a** Pathway diagram of MZ1-dependent proteins predominantly biotinylated by VHL-AirID in MM1.S cells. Each edge represents a protein–protein interaction based on the STRING database. **b** Pathway diagram of ARV-825-dependent proteins predominantly biotinylated by AirID-CRBN in MM1.S cells. Each edge represents a protein–protein interaction based on the STRING database. **c** List of common proteins biotinylated by AirID-CRBN or VHL-AirID dependent on ARV-825 or MZ1. **d** Gene ontology (GO) analysis of proteins biotinylated by AirID-CRBN and VHL-AirID dependent on ARV-110 and MZ1 in MM1.S cells. GO analysis was performed using the ShinyGO software. The horizontal axis represents $-\log_{10}$ (FDR), the colour change represents the number of genes, and the size of the circle represents fold enrichment. GO biological process analysis was performed.

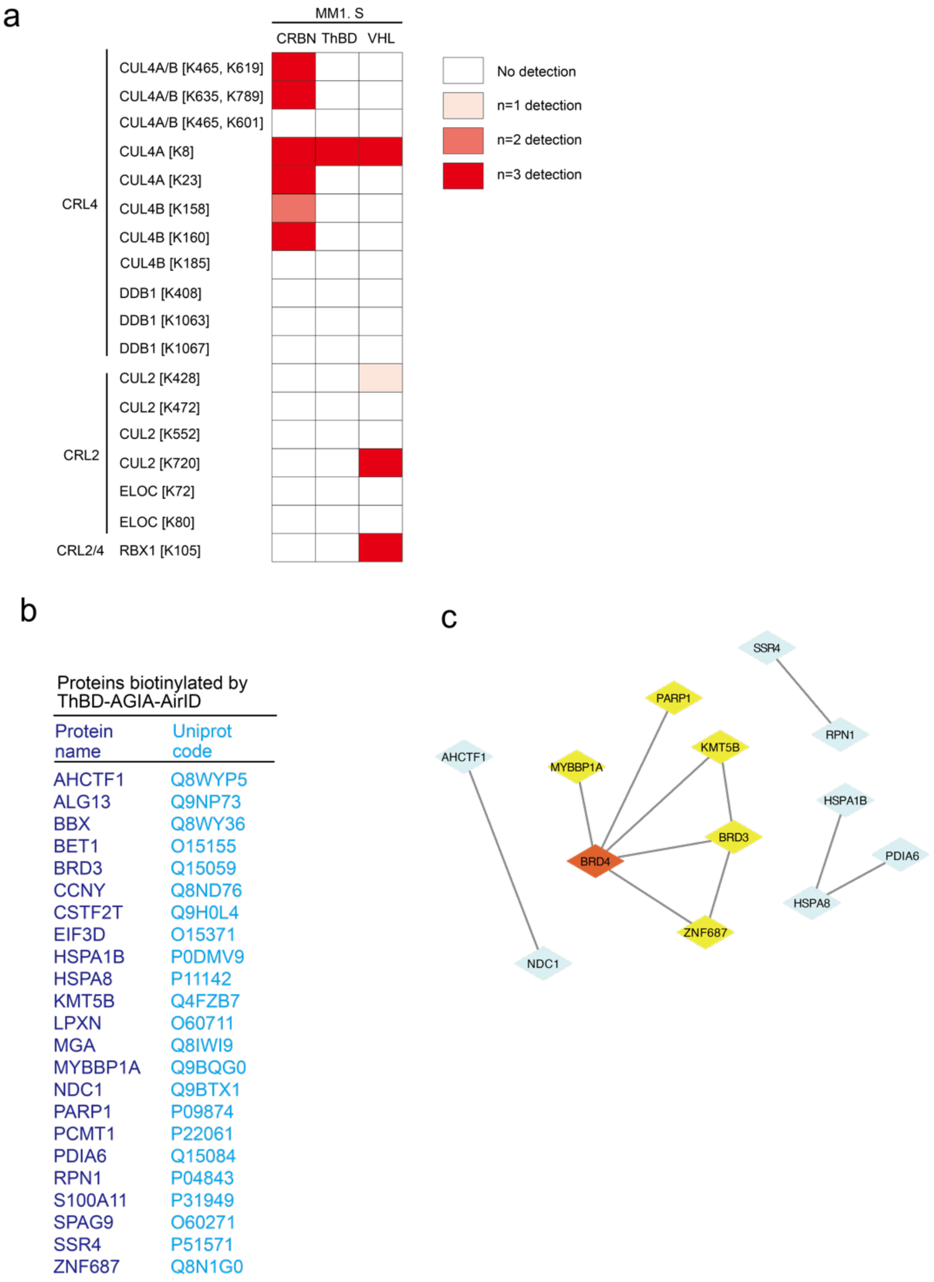


**Supplementary Fig. 3 ARV-825-dependent protein biotinylation using MM1.S cells expressing AirID-fused ThBD**

**a** Heat map showing CRL2/4 complex proteins biotinylated by AirID-CRBN, ThBD-AGIA-AirID, or VHL-AirID in MM1.S cells. Heatmap diagrams were generated based on the data from each cell line treated with DMSO. The colour change represents the number of biotinylated peptides detected. **b** List of ARV-825-dependent proteins predominantly biotinylated by ThBD-AGIA-AirID in MM1.S cells (ARV-825/pomalidomide peptides ratios >2, $P < 0.05$). **c** Pathway diagram of ARV-825-dependent proteins predominantly biotinylated by ThBD-AGIA-AirID in MM1.S cells. Each edge represents a protein–protein interaction based on the STRING database.

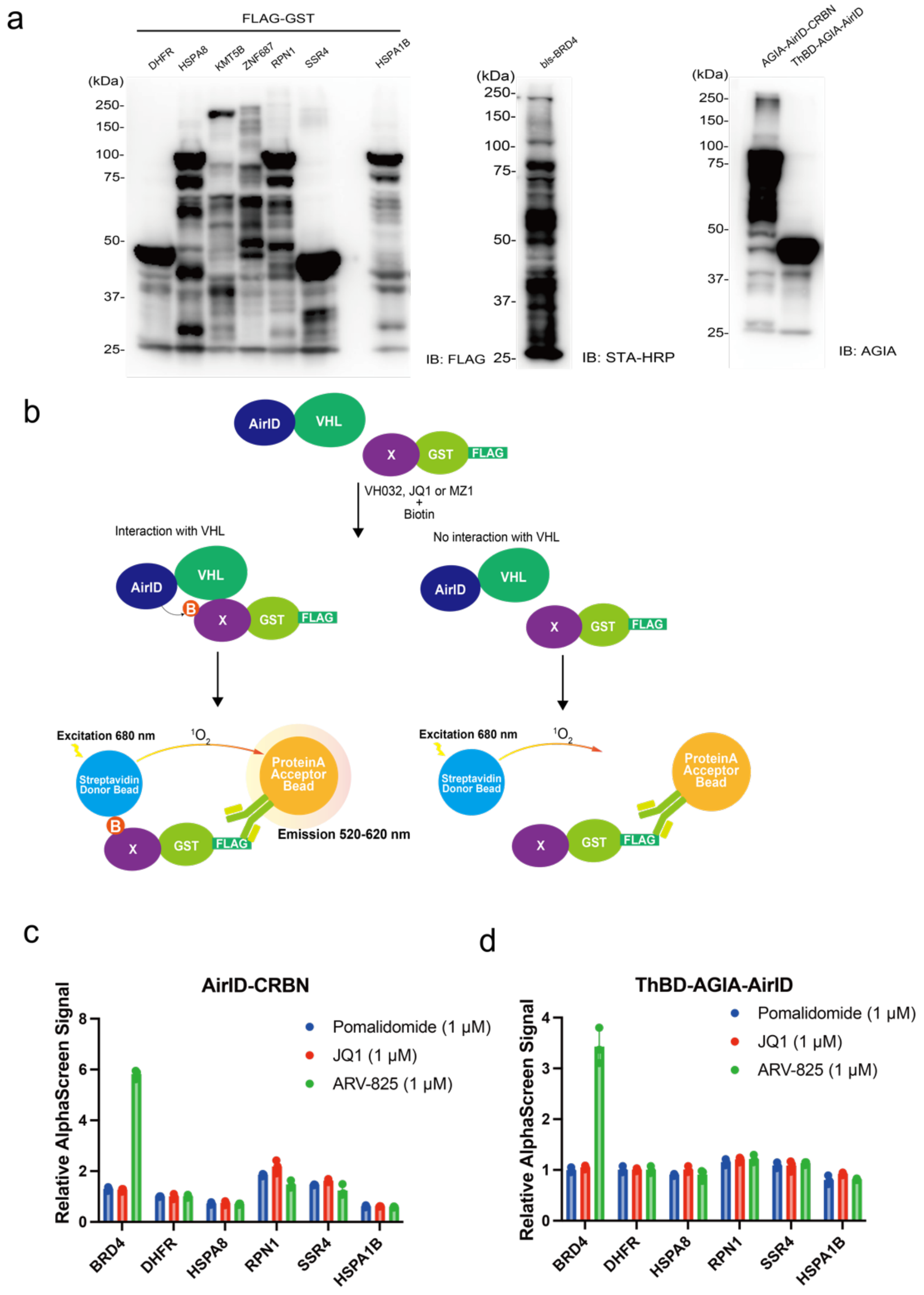


**Supplementary Fig. 4 ARV-825 and MZ1-dependent biotinylated protein synthesis in a wheat cell-free system and detection of biotinylated proteins using the AlphaScreen method**

**a** Expression of the indicated proteins in a wheat cell-free system was confirmed using immunoblotting. **b** Schematic of the detection of proteins biotinylated by VHL-AirID with VH032, JQ1, or MZ1 using the AlphaScreen assay. **c**, **d** AlphaScreen detection of proteins biotinylated by AirID-CRBN or ThBD-AGIA-AirID in vitro (c, AirID-CRBN; d, ThBD-AGIA-AirID). All AlphaScreen signals represent the relative values of FLAG-GST-DHFR with AirID-CRBN or ThBD-AGIA-AirID for each drug treatment. Error bars denote standard deviations (independent experiments, $n = 3$).

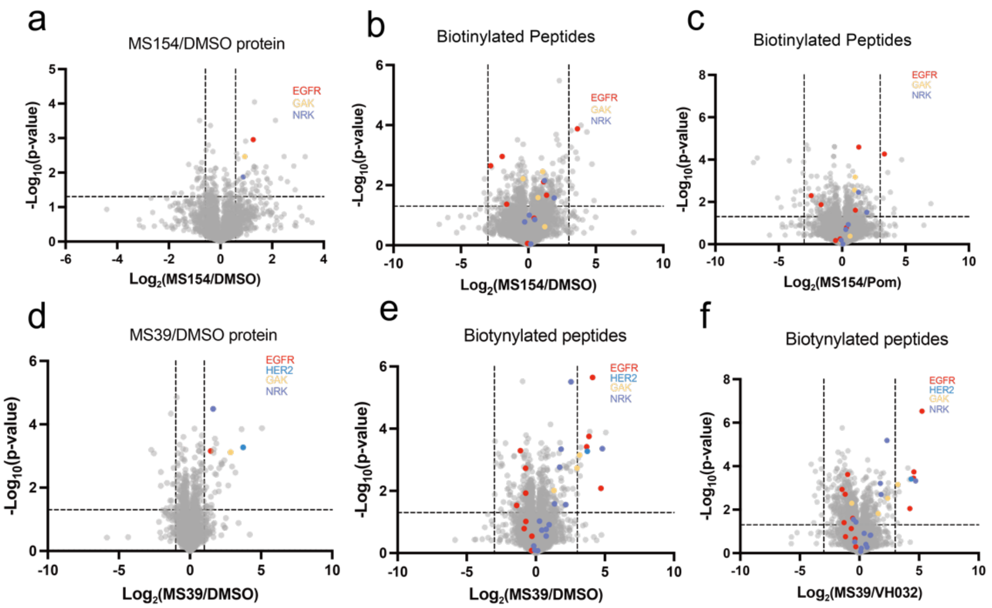


g

Proteins biotinylated by
ThBD-AGIA-AirID with MS154

| Protein name | Uniprot code |
|---|---|
| AKT1S1 | Q96B36 |
| ARHGEF12 | Q9NZN5 |
| CDC42BPA | Q5VT25 |
| COPS7A | Q9UBW8 |
| EGFR | P00533 |
| EIF2S1 | P05198 |
| ERRFI1 | Q9UJM3 |
| FHIP1A | Q05DH4 |
| GAK | O14976 |
| GTF2F2 | P13984 |
| HJURP | Q8NCD3 |
| HP1BP3 | Q5SSJ5 |
| LAT2 | Q9GZY6 |
| MYO18A | Q92614 |
| NANS | Q9NR45 |
| NCAPH | Q15003 |
| NRK | Q7Z2Y5 |
| NUP155 | O75694 |
| NUP54 | Q7Z3B4 |
| PABPC4 | Q13310 |
| PARD3 | Q8TEW0-2 |
| PLEKHG1 | Q9ULL1 |
| RBM26 | Q5T8P6-2 |
| RPS9 | P46781 |
| SLC4A7 | Q9Y6M7-7 |
| SMARCD1 | Q96GM5 |
| SPATA20 | Q8TB22 |
| SRSF2 | Q01130 |
| STK24 | Q9Y6E0 |
| TMEM185B | Q9H7F4 |
| TMEM51 | Q9NW97 |
| TTC27 | Q6P3X3 |
| WDR55 | Q9H6Y2 |
| ZNF724 | A8MTY0 |

h

Proteins biotinylated by
VHL-AirID with MS39

| Protein name | Uniprot code | Protein name | Uniprot code |
|---|---|---|---|
| ATG2B | Q96BY7 | PC | P11498 |
| CCNY | Q8ND76 | PCBP1 | Q15365 |
| CDA | P32320 | PCDH1 | Q08174-2 |
| COL17A1 | Q9UMD9 | PCDH7 | O60245 |
| COPA | P53621 | POGK | Q9P215 |
| CYB5B | O43169 | PRPF4 | O43172 |
| DSC2 | Q02487 | PSD3 | Q9NYI0 |
| DSG2 | Q14126 | PSEN1 | P49768 |
| EGFR | P00533 | RAB27B | O00194 |
| EMD | P50402 | RASEF | Q8IZ41 |
| EPB41L4B | Q9H329 | RIPK2 | O43353 |
| ERBB2 | P04626 | SCRN3 | Q0VDG4 |
| ERRFI1 | Q9UJM3 | SHB | Q15464 |
| FAM120A | Q9NZB2 | SHC1 | P29353 |
| FAM83F | Q8NEG4 | SLITRK6 | Q9H5Y7 |
| FLVCR1 | Q9Y5Y0 | SRSF11 | Q05519 |
| GAK | O14976 | STX6 | O43752 |
| GCFC2 | P16383 | TMC5 | Q6UXY8 |
| GPRC5A | Q8NFJ5 | TMEM51 | Q9NW97 |
| GPRC5C | Q9NQ84 | TMEM87A | Q8NBN3 |
| GSDME | O60443 | UACA | Q9BZF9 |
| HTT | P42858 | USP48 | Q86UV5 |
| IGF2R | P11717 | VAMP1 | P23763 |
| ITGB1 | P05556 | VAMP5 | O95183 |
| KDF1 | Q8NAX2 | ZDHHC5 | Q9C0B5 |
| KIRREL1 | Q96J84 | ZFYVE16 | Q7Z3T8 |
| LAT2 | Q9GZY6 | | |
| MAP3K20 | Q9NYL2 | | |
| MARK3 | P27448 | | |
| MUC13 | Q9H3R2 | | |
| NPM3 | O75607 | | |
| NRK | Q7Z2Y5 | | |
| PABPC4 | Q13310 | | |
| PAK6 | Q9NQU5 | | |

i

Proteins biotinylated by
ThBD-AGIA-AirID with MS154 (GO molecular Function)
Ratio(MS154/Pom) > 1.5, $P < 0.05$

j

Proteins biotinylated by
VHL-AirID with MS39 (GO molecular Function)
Ratio(MS39/VHL) > 1.5, $P < 0.05$

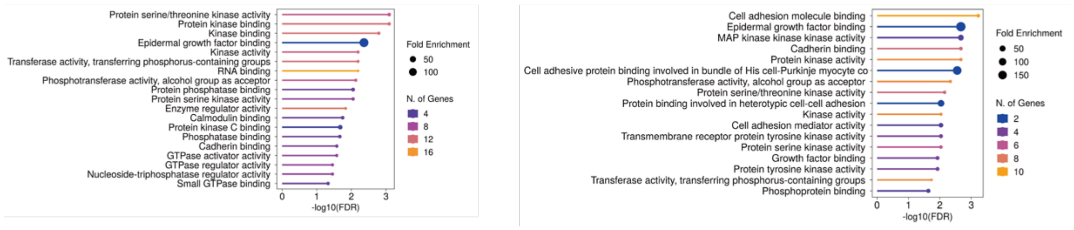


**Supplementary Fig. 5 Analysis of proteins biotinylated by ThBD-AGIA-AirID or VHL-AirID dependent of MS154 or MS39**

**a** Volcano plot of MS154-dependent proteins biotinylated detected via LC-MS/MS in HCC827-Luc cells stably expressing ThBD-AGIA-AirID. **b**, **c** Volcano plots of MS154-dependent biotinylated peptides detected via LC-MS/MS in cells stably expressing ThBD-AGIA-AirID. Volcano plot comparing **b** MS154 treatment with DMSO treatment; and **c** MS154 treatment with pomalidomide treatment. **d** Volcano plot of MS39-dependent biotinylated proteins detected via LC-MS/MS in HCC827-Luc cells stably expressing VHL-AirID. **e, f** Volcano plots of MS39-dependent biotinylated peptides detected via LC-MS/MS in cells stably expressing VHL-AirID. Volcano plot comparing **e** MS39 treatment with DMSO treatment; and **f** MS39 treatment with VH032 treatment. **g**, **h** List of MS154- or MS39-dependent proteins predominantly biotinylated by ThBD-AGIA-AirID or VHL-AirID in HCC827-Luc cells (MS154/pomalidomide proteins ratios >1.5, $P < 0.05$) (MS39/VH032 proteins ratios >1.5, $P < 0.05$). **i, j** Gene ontology (GO) analysis of proteins biotinylated by ThBD-AGIA-AirID and VHL-AirID dependent on MS154 and MS39 in HCC827-Luc cells. GO analysis was performed using the ShinyGO software. The horizontal axis represents $-\log_{10}$ (FDR), the colour change represents the number of genes, and the size of the circle represents fold enrichment. GO molecular function analysis was performed. **a, b**, **c, d, e, f** Significant changes in the volcano plots were calculated using Student's two-sided *t*-test, and the false discovery rate (FDR)-adjusted *P*-values calculated using Benjamini–Hochberg method are shown in Supplementary Data 3-6.

**a**

Ratio (VH032/DMSO) > 2.0 and
Ratio (MZ1/DMSO) > 2.0
(*P*<0.05)

| Protein name | Uniprot code |
|---|---|
| SCRN3 | Q0VDG4 |
| SLC25A5 | P05141 |

Ratio (MZ1/VH032) > 2.0
(*P*<0.05)

**b**

Ratio (VH032/DMSO) > 1.5 and
Ratio (MS39/DMSO) > 1.5
(*P*<0.05)

| Protein name | Uniprot code |
|---|---|
| AGK | Q53H12 |
| COPB2 | P35606 |
| MAD2L1BP | Q15013 |
| NPLOC4 | Q8TAT6 |
| PARG | Q86W56 |
| TIPRL | O75663 |
| TUBA1B | P68363 |
| TUBB | P07437 |
| TUBB4B | P68371 |
| WDR11 | Q9BZH6 |
| ZNF114 | Q8NC26 |

Ratio (MS39/VH032) > 1.5
(*P*<0.05)

**c**

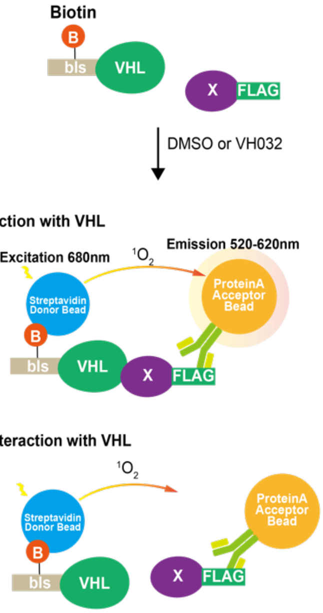


**d**

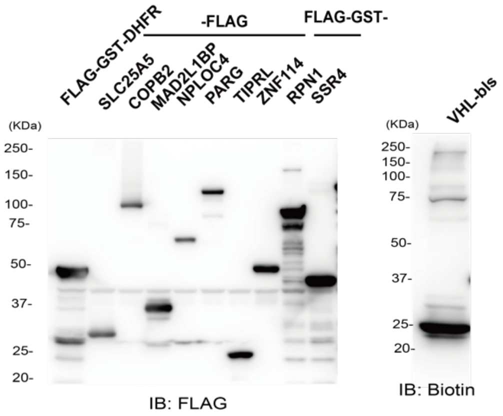


**e**

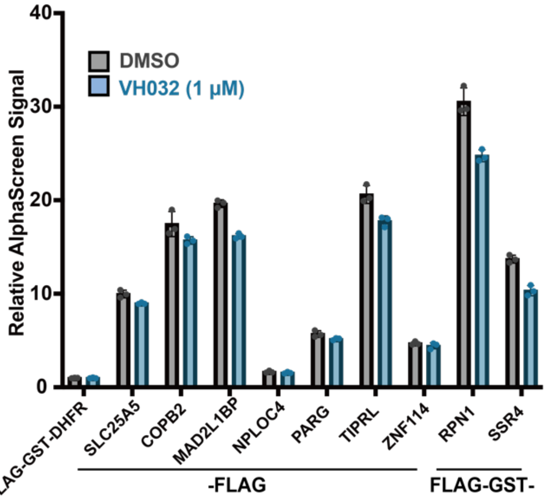


**Supplementary Fig. 6 Neo-substrate exploration of VHL binder VH032**

**a** List of VH032-dependent proteins predominantly biotinylated by VHL-AirID in MM1.S cells (VH032/DMSO and MZ1/DMSO proteins ratios >2, $P < 0.05$). **b** List of VH032-dependent proteins predominantly biotinylated by VHL-AirID in HCC-827-Luc cells (VH032/DMSO and MS39/DMSO proteins ratios >1.5, $P < 0.05$). **c** Schematic of AlphaScreen detection of the interaction between VHL and VHL candidate neosubstrates via VH032. **d** Expression confirmed in a synthetic wheat cell-free system of proteins used in AlphaScreen. **e** AlphaScreen assay detection of the VHL-bls interaction protein. All VHL-bls interaction analyses with candidate VHL-interacting proteins were performed in the presence of DMSO or VH032. All AlphaScreen signals represent the relative values of FG-DHFR and VHL-bls proteins.

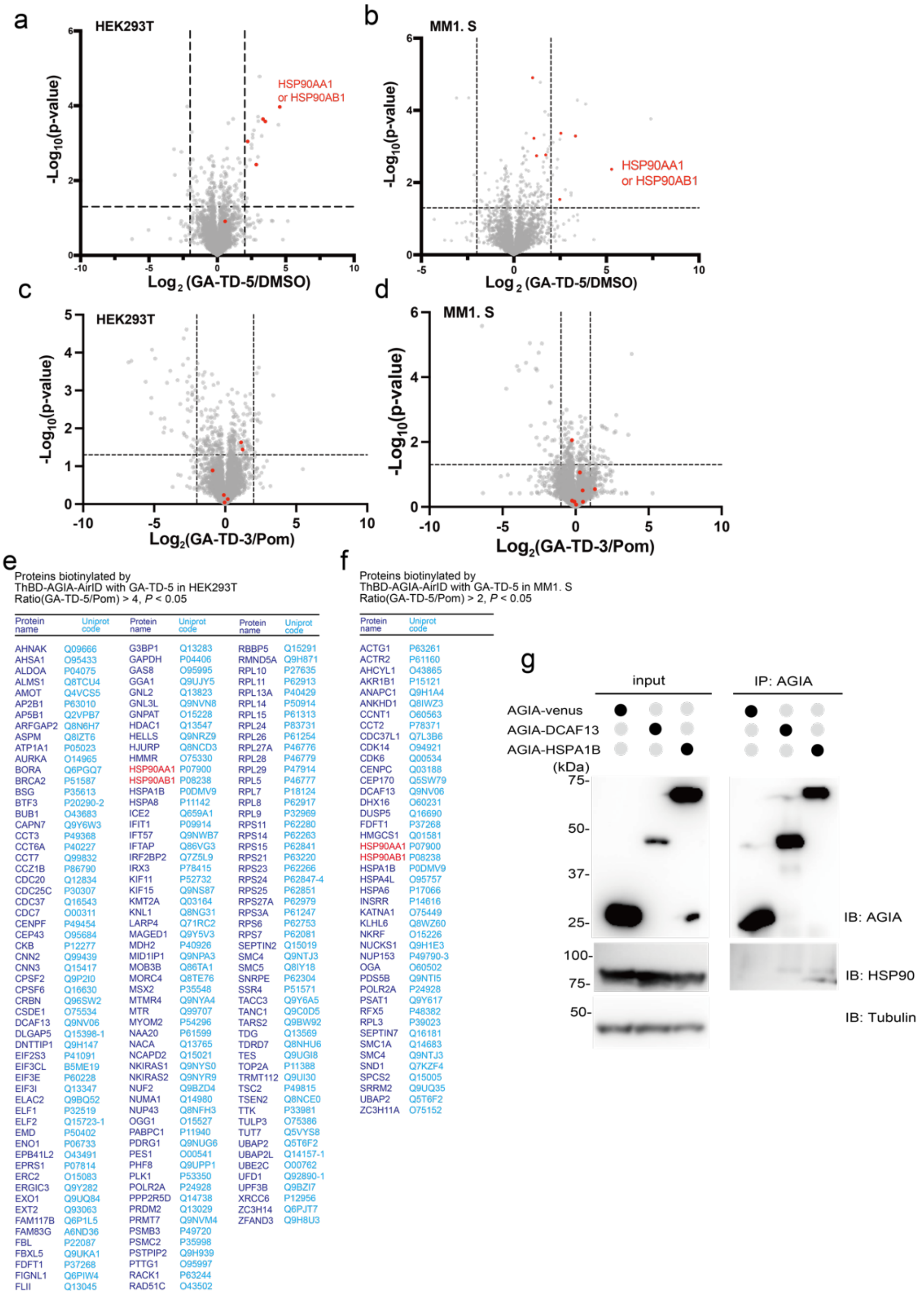


**Supplementary Fig. 7 Analysis of proteins biotinylated by ThBD-AGIA-AirID dependent on GA-TD-5 or GA-TD-3**

**a, b** Volcano plots of GA-TD-5-dependent biotinylated peptides detected via LC-MS/MS in HEK293T or MM1.S cells stably expressing ThBD-AGIA-AirID. **c, d** Volcano plots of GA-TD-3-dependent biotinylated peptides detected via LC-MS/MS in HEK293T or MM1.S cells stably expressing ThBD-AGIA-AirID. **e, f** List of GA-TD-5-dependent proteins predominantly biotinylated by ThBD-AGIA-AirID in HEK293T or MM1.S cells (e, GA-TD-5/Pomalidomide peptides ratios > 4, $P$ < 0.05; f, GA-TD-5/Pomalidomide peptides ratios > 2, $P$ < 0.05). **g** Co-immunoprecipitation. HEK293T cells were transfected with each gene. The transfected Venus gene was used as a control. Immunoprecipitation was performed using an anti-AGIA antibody. The immunoprecipitated proteins were detected using anti-AGIA and anti-HSP90 antibodies. **a, b**, **c, d** Significant changes in the volcano plots were calculated using Student's two-sided *t*-test, and the false discovery rate (FDR)-adjusted *P*-values calculated using Benjamini–Hochberg method are shown in Supplementary Data 7-8.

a

Proteins biotinylated by
ThBD-AGIA-AirID with ARV-110 in LNCaP
Ratio(ARV-110/Pom) > 1, $P < 0.05$

| Protein name | Uniprot code | Protein name | Uniprot code | Protein name | Uniprot code |
|---|---|---|---|---|---|
| ABI1 | Q8IZP0 | LAD1 | O00515 | SCRIB | Q14160 |
| ACIN1 | Q9UKV3 | LIMCH1 | Q9UPQ0 | SF3B1 | O75533 |
| ACIN1 | Q9UKV3-5 | MAD1L1 | Q9Y6D9 | SHROOM2 | Q13796 |
| AIMP2 | Q13155 | MAP2 | P11137-4 | SKI | P12755 |
| ANAPC1 | Q9H1A4 | MARK3 | P27448 | SLC16A1 | P53985 |
| ANKLE2 | Q86XL3 | MAVS | Q7Z434 | SLC35A2 | P78381-4 |
| AR | P10275 | MED1 | Q15648 | SNRNP200 | O75643 |
| ARHGAP21 | Q5T5U3 | MME | P08473 | SOAT1 | P35610 |
| ARL6IP5 | O75915 | MRE11 | P49959 | SORBS2 | O94875 |
| ATP2A2 | P16615 | MTA1 | Q13330 | SPTBN2 | O15020 |
| BCAP31 | P51572 | MYBBP1A | Q9BQG0-2 | SRSF11 | Q05519 |
| BICRA | Q9NZM4 | MYEF2 | Q9P2K5 | SSX2IP | Q9Y2D8 |
| BMS1 | Q14692 | MYO18A | Q92614 | STEAP1 | Q9UHE8 |
| BRD4 | O60885 | N4BP2 | Q86UW6 | SUCO | Q9UBS9 |
| CACTIN | Q8WUQ7 | NBN | O60934 | SUGP1 | Q8IWZ8 |
| CCDC47 | Q96A33 | NCOA2 | Q15596 | SURF6 | O75683 |
| CCDC86 | Q9H6F5 | NEBL | O76041-2 | SYNE2 | Q8WXH0 |
| CD2AP | Q9Y5K6 | NELFA | Q9H3P2 | TAF15 | Q92804 |
| CDCA8 | Q53HL2 | NFIB | O00712 | TBL3 | Q12788 |
| CDKN2AIP | Q9NXV6 | NFXL1 | Q6ZNB6 | TDRKH | Q9Y2W6 |
| CDV3 | Q9UKY7 | NOL11 | Q9H8H0 | THOC3 | Q96J01 |
| CEBPZ | Q03701 | NOL6 | Q9H6R4 | TMEM131 | Q92545 |
| CGNL1 | Q0VF96 | NOP58 | Q9Y2X3 | TMEM230 | Q96A57 |
| CHD2 | O14647 | NUP107 | P57740 | TMPO | P42166 |
| CLCC1 | Q96S66 | NUP153 | P49790-3 | TOX4 | O94842 |
| CMAS | Q8NFW8 | NUP155 | O75694 | TRA2A | Q13595 |
| CNNM3 | Q8NE01 | NUP214 | P35658 | TRDN | Q13061 |
| COA7 | Q96BR5 | NVL | O15381 | USO1 | O60763 |
| COBLL1 | Q53SF7 | ORC1 | Q13415 | USP22 | Q9UPT9 |
| CWC15 | Q9P013 | ORC4 | O43929 | USP30 | Q70CQ3 |
| DCAF8 | Q5TAQ9 | PARK7 | Q99497 | UTRN | P46939 |
| DDRGK1 | Q96HY6 | PC | P11498 | VAPB | O95292 |
| DDX31 | Q9H8H2 | PDCD11 | Q14690 | VIM | P08670 |
| DKC1 | O60832 | PDS5A | Q29RF7 | VTI1B | Q9UEU0 |
| DNTTIP2 | Q5QJE6 | PELP1 | Q8IZL8 | WDR43 | Q15061 |
| DOCK7 | Q96N67-5 | PEX14 | O75381 | WDR75 | Q8IWA0 |
| DPF2 | Q92785 | PIKFYVE | Q9Y2I7 | ZC3H18 | Q86VM9 |
| DSN1 | Q9H410 | PNISR | Q8TF01 | ZC3HC1 | Q86WB0 |
| ESF1 | Q9H501 | POLR2A | P24928 | ZNF326 | Q5BKZ1 |
| EXOSC9 | Q06265 | POLR3A | O14802 | | |
| FAM120C | Q9NX05 | PPP4R3A | Q6IN85 | | |
| FBL | P22087 | PPRC1 | Q5VV67 | | |
| FLVCR1 | Q9Y5Y0 | PRPF31 | Q8WWY3 | | |
| FMR1 | Q06787 | PSPC1 | Q8WXF1 | | |
| FOXA1 | P55317 | RAB11FIP5 | Q9BXF6 | | |
| GRHL2 | Q6ISB3 | RAD50 | Q92878 | | |
| HABP2 | Q14520 | RAI14 | Q9P0K7 | | |
| HEATR1 | Q9H583 | RBM15 | Q96T37 | | |
| HMGA1 | P17096 | RBM19 | Q9Y4C8 | | |
| HNRNPA2B1 | P22626 | RBM26 | Q5T8P6 | | |
| HNRNPF | P52597 | RBM34 | P42696 | | |
| ILF2 | Q12905 | RECQL | P46063 | | |
| INTS1 | Q8N201 | REPS1 | Q96D71 | | |
| ITGB1 | P05556 | REXO4 | Q9GZR2 | | |
| ITPRID2 | P28290 | RIDA | P52758 | | |
| KAT14 | Q9H8E8 | RRP9 | O43818 | | |
| KAT7 | O95251 | RSBN1 | Q5VWQ0 | | |
| KCNMA1 | Q12791 | SAP130 | Q9H0E3-3 | | |
| KCTD18 | Q6PI47 | SARG | Q9BW04 | | |
| KRT18 | P05783 | SART3 | Q15020 | | |
| KRT8 | P05787 | SCARB1 | Q8WTV0-2 | | |

b

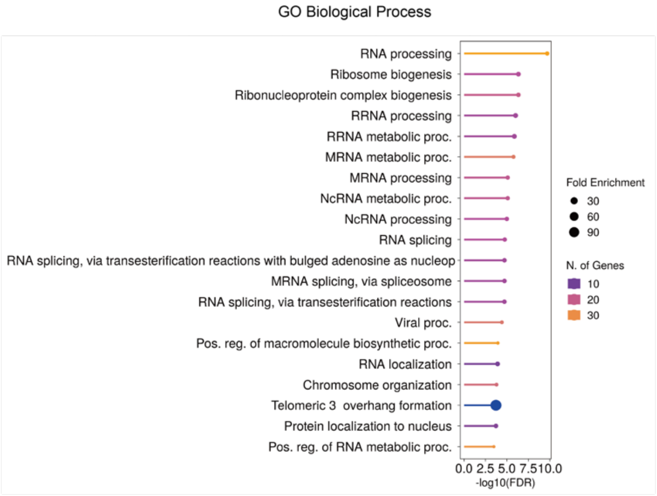


c

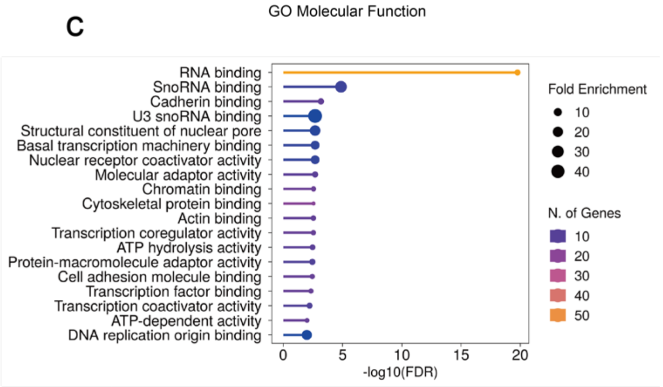


**Supplementary Fig. 8 List of ARV-110-dependent proteins biotinylated by ThBD-AGIA-AirID and gene ontology analysis**

**a** List of ARV-110-dependent proteins predominantly biotinylated by ThBD-AGIA-AirID in LNCaP cells (ARV-110/Pomalidopmide proteins ratio >1, $P < 0.05$). **b, c** Gene ontology (GO) analysis of proteins biotinylated by ThBD-AGIA-AirID dependent on ARV-110 in LNCaP cells. GO analysis was performed using the ShinyGO software. The horizontal axis represents $-\log_{10}$ (FDR), the colour change represents the number of genes, and the size of the circle represents fold enrichment. (b, GO biological process analysis; c, GO molecular function analysis).

### General information for chemical synthesis

Proton nuclear magnetic resonance ($^{1}$H NMR) spectra were recorded with tetramethylsilane ($\delta_H$ 0.00) as an internal standard. Coupling constants (*J*) are reported in hertz (Hz). Abbreviations of multiplicity are as follows: s, singlet; d, doublet; t, triplet; quint, quintet; m, multiplet; br, broad. The data are presented as follows: chemical shift, multiplicity, coupling constants, and integration. Carbon nuclear magnetic resonance ($^{13}$C NMR) spectra were recorded with $CDCl_3$ ($\delta_C$ 77.16) as an internal standard. Column chromatography was carried out on silica gel 60 N (63–210 μm or 40–50 μm). Analytical thin layer chromatography (TLC) was carried out with 0.25-mm silica gel plates. Visualization was accomplished with ultraviolet light and anisaldehyde or phosphomolybdic acid stain, followed by heating. Reagents and solvents were purified by standard means or used as received, unless otherwise noted. Dehydrated tetrahydrofuran (THF, stabilizer-free) and *N*,*N*-dimethylformamide (DMF) were purchased. All reactions were conducted in an argon atmosphere, unless otherwise noted.

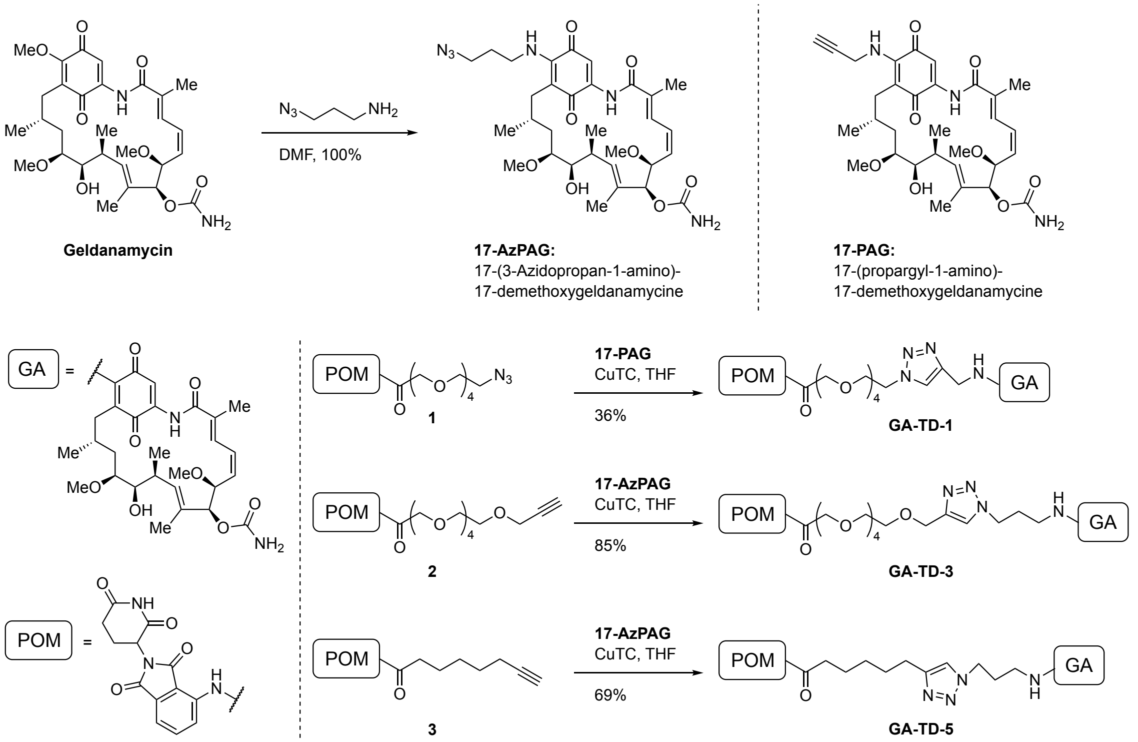

**Scheme 1.**

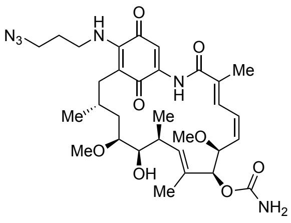


**17-(3-Azidopropan-1-amino)-17-demethoxygeldanamycine (17-AzPAG):** 3-Azidopropan-1-amine (31.3 mg, 0.312 mmol) in DMF (2.0 mL) was added to geldanamycin (98.4 mg, 0.176 mmol) in the round bottom flask, and the mixture was stirred for 12 h in the dark. The reaction mixture was concentration in vacuo followed by column chromatography ($CHCl_3$ → 50:1 $CHCl_3$/MeOH) afforded **17-AzPAG** (118 mg, containing DMF, >100%) as a purple solid. **17-AzPAG** was used without further purifications.

$^1$H NMR (594 MHz, $CDCl_3$) δ 9.16 (s, 1H), 7.29 (s, 1H), 6.96 (d, *J* = 11.6 Hz, 1H), 6.59 (t, *J* = 11.6 Hz, 1H), 6.35 (t, J = 5.7 Hz, 1H), 5.88 (dd, *J* = 11.6, 9.8 Hz, 1H), 5.86 (d, *J* = 10.5 Hz, 1H), 5.18 (s, 1H), 5.06 (brs, 2H), 4.32 (d, J = 9.8 Hz, 1H), 4.18 (brs, 1H), 3.68 (m, 1H), 3.63–3.56 (m, 2H), 3.53–3.42 (m, 3H), 3.37 (s, 3H), 3.27 (s, 3H), 2.75 (m, 1H), 2.70 (d, J = 14.0 Hz, 1H), 2.40 (dd, J = 14.0, 10.7 Hz, 1H), 2.03 (s, 3H), 1.94 (m, 3H), 1.80 (m, 5H), 1.73 (s, 1H), 1.00 (t, J = 6.6 Hz, 3H), 0.99 (t, J = 6.6 Hz, 3H); $^{13}$C NMR (149 MHz, $CDCl_3$) δ 183.9, 180.9, 168.4, 156.3, 144.8, 141.3, 136.0, 135.0, 133.7, 133.0, 127.1, 126.6, 109.0, 108.8, 81.6, 81.5, 81.3, 72.7, 57.1, 56.8, 48.9, 43.3, 35.1, 34.4, 32.4, 29.2, 28.7, 22.9, 12.9, 12.7, 12.5; HRMS (FAB) *m*/*z* $[M + K]^+$ calcd for $C_{31}H_{44}N_6O_8K$ 667.2852; found 667.2826.

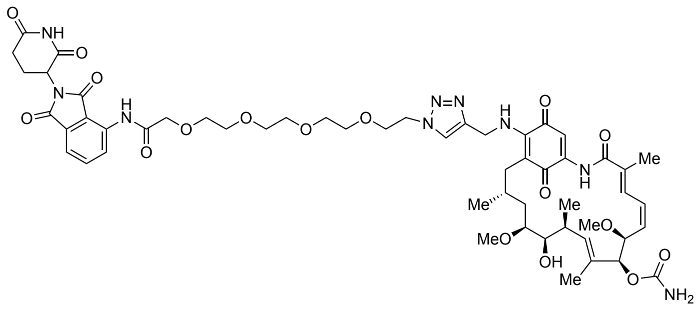


**GA-TD-1**: CuTC (8.27 mg, 43.4 μmol) was added to a mixture of **17-PAG** (24.7 mg, 42.3 μmol) and azide **1** (23.5 mg, 43.9 μmol) in THF (2.0 mL). After 30 min of stirring, the reaction was quenched with half saturated aqueous $NH_4Cl$ (20 mL), and the resulting mixture was extracted with $CHCl_3$ (20 mL). The organic extract was dried over anhydrous $Na_2SO_4$. Filtration and evaporation in vacuo furnished the crude product, which was purified by flash column chromatography (silica gel 10 g, a concave concentration gradient of MeOH in $CHCl_3$) followed by preparative thin layer chromatography ($CHCl_3$/MeOH 9:1) to give **GA-TD-1** (17.5 mg, 36%) as a purple solid. The compound exists as a 1:1 mixture of diastereomers arising from the racemic E3 ligand.

$^1$H NMR (594 MHz, $CDCl_3$) δ 10.46 (s, 0.5H), 10.41 (s, 0.5H), 9.21 (brs, 0.5H), 9.20 (s, 0.5H), 9.14 (s, 0.5H), 8.98 (brs, 0.5H), 8.82 (d, *J* = 8.5 Hz, 0.5H), 8.80 (d, *J* = 8.5 Hz, 0.5H), 7.85 (s, 0.5H), 7.79 (s, 0.5H), 7.71 (dd, *J* = 8.5, 7.3 Hz, 0.5H), 7.70 (dd, *J* = 8.5, 7.3 Hz, 0.5H) 7.57 (d, *J* = 7.3 Hz, 1H), 7.53 (d, *J* = 7.3 Hz, 1H), 7.24 (s, 0.5H), 7.15 (s, 0.5H), 6.94 (d, *J* = 11.4 Hz, 0.5H), 6.93 (d, *J* = 11.4 Hz, 0.5H), 6.64 (m, 1H), 6.56 (t, *J* = 11.4 Hz, 1H), 5.89 (m, 1H), 5.85 (m, 1H), 5.26 (s, 0.5H), 5.21 (s, 0.5H), 5.05–4.76 (m, 5H), 4.56 (m, 2H), 4.36 (brs, 0.5H), 4.32 (d, *J* = 9.7 Hz, 1H), 4.26–4.07 (m, 2.5H), 3.94–3.54 (m, 15H), 3.43 (m, 1H), 3.36 (s, 1.5H), 3.35 (s, 1.5H), 3.28 (s, 1.5H), 3.27 (s, 1.5H), 2.88 (m, 1H), 2.83–2.61 (m, 4H), 2.42 (m, 1H), 2.15 (m, 1H), 1.99 (s, 1.5H), 1.98 (s, 1.5H), 1.80 (s, 1.5H), 1.77–1.68 (m, 3H), 1.03–0.97 (m, 6H); $^{13}$C NMR (149 MHz, $CDCl_3$) δ 184.1, 184.0, 181.2, 181.1, 171.4, 171.3, 169.5, 169.3, 168.7, 168.6, 168.5, 168.30, 168.26, 166.8, 166.7, 156.3, 156.2, 144.8, 144.6, 143.7, 143.5, 141.13, 141.11, 136.9, 136.8, 136.54, 136.46, 136.2, 134.9, 134.6, 134.1, 133.9, 133.0, 132.8, 131.5,

131.4, 127.6, 127.3, 126.7, 126.6, 125.3, 123.6, 123.5, 119.0, 116.3, 116.2, 109.9, 109.7, 109.1, 82.2, 81.9, 81.57, 81.56, 81.43, 81.38, 72.74, 72.68, 71.7, 71.2, 71.1, 70.83, 70.80, 70.79, 70.75, 70.72, 70.68, 70.57, 70.52, 70.49, 69.4, 57.3, 56.9, 50.6, 49.4, 49.3, 41.7, 35.3, 35.1, 34.6, 32.5, 32.3, 31.5, 28.8, 28.6, 23.2, 23.1, 23.0, 22.9, 12.94, 12.89, 12.7, 12.5, 12.4; HRMS (FAB) *m*/*z* $[M + H]^+$ calcd for $C_{54}H_{70}N_9O_{17}$ 1116.4884; found 1116.4889.

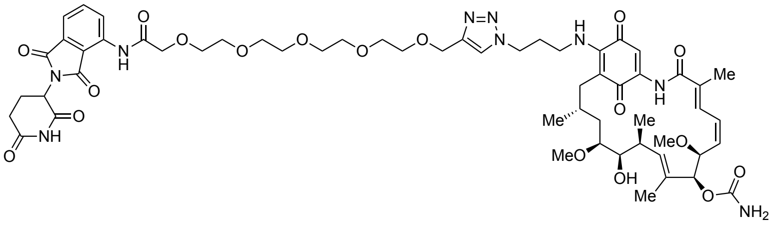

**GA-TD-3**: CuTC (7.2 mg, 37.8 μmol) was added to a mixture of **17-AzPAG** (20.3 mg, 32.3 μmol) and alkyne **2** (20.7 mg, 37.9 μmol) in THF (1.0 mL). After 1 h of stirring, the reaction was quenched with half saturated aqueous $NH_4Cl$ (20 mL), and the resulting mixture was extracted with $CHCl_3$ (2 × 20 mL). The organic extracts were dried over anhydrous $Na_2SO_4$. Filtration and evaporation in vacuo furnished the crude product, which was purified by flash column chromatography ($CHCl_3$/MeOH 100:3) followed by preparative thin layer chromatography ($CHCl_3$/MeOH 9:1) to give **GA-TD-3** (17.3 mg, 36%) as a purple solid. The compound exists as a 1:1 mixture of diastereomers arising from the racemic E3 ligand. $^1$H NMR (600 MHz, $CDCl_3$) δ 10.50 (s, 0.5H), 10.49 (s, 0.5H), 9.18–9.06 (m, 2H), 8.83 (m, 1H), 7.71 (m, 1H), 7.65 (s, 1H), 7.56 (m, 1H), 7.25 (s, 0.5H), 7.24 (s, 0.5H), 6.94 (d, *J* = 11.7 Hz, 1H), 6.57 (m, 1H), 6.32 (t, *J* = 6.0 Hz, 1H), 5.88 (m, 1H), 5.87 (m, 1H), 5.21 (s, 1H), 4.95 (m, 1H), 4.67 (s, 2H), 4.47 (t, *J* = 6.7 Hz, 2H), 4.31 (d, *J* = 9.8 Hz, 1H), 4.25–4.09 (m, 3H), 3.85–3.62 (m, 16H), 3.61–3.53 (m, 3H), 3.43 (m, 1H), 3.36 (s, 3H), 3.27 (s, 3H), 2.88 (m, 1H), 2.84–2.74 (m, 2H), 2.73 (m, 1H), 2.64 (m, 1H), 2.26 (m, 2H), 2.22 (m, 1H), 2.15 (m, 1H), 2.02 (s, 3H), 1.79 (s, 3H), 1.75 (m, 2H), 1.67 (m, 1H), 0.996 (d, *J* = 7.0 Hz, 1.5H), 0.994 (d, *J* = 7.0 Hz, 1.5H), 0.909 (d, *J* = 6.7 Hz, 1.5H), 0.904 (d, *J* = 6.7 Hz,

1.5H); $^{13}$C NMR (151 MHz, $CDCl_3$) δ 184.07, 184.05, 181.03, 181.02, 171.42, 171.40, 169.5, 169.4, 168.60, 168.59, 168.57, 168.56, 168.3, 166.95, 166.92, 156.21, 156.19, 145.8, 144.82, 144.80, 141.24, 141.22, 136.89, 136.88, 136.45, 136.42, 136.2, 134.98, 134.96, 133.9, 132.9, 131.55, 131.54, 127.3, 126.7, 126.6, 125.3, 123.0, 118.9, 116.30, 116.29, 109.6, 109.5, 109.04, 109.02, 81.8, 81.6, 81.3, 72.7, 71.84, 71.81, 71.08, 71.06, 70.78, 70.76, 70.75, 70.62, 70.60, 70.58, 70.57, 70.56, 69.90, 69.89, 64.7, 57.3, 56.9, 49.4, 47.4, 42.6, 35.13, 35.12, 34.4, 32.47, 32.46, 31.6, 30.42, 30.39, 28.8, 23.0, 22.89, 22.86, 13.0, 12.7, 12.6; HRMS (FAB) *m/z* $[M + K]^+$ calcd for $C_{57}H_{75}N_9O_{18}K$ 1212.4862; found 1212.4865.

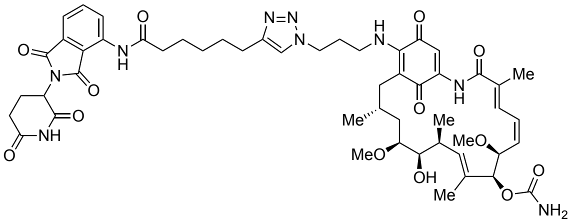


**GA-TD-5**: CuTC (3.3 mg, 17 μmol) was added to a mixture of **17-AzPAG** 19.6 mg, 31.2 μmol) and alkyne **3** (16.7 mg, 42.2 μmol) in THF (1.0 mL). After 30 min of stirring, the reaction was quenched with half saturated aqueous $NH_4Cl$ (20 mL), and the resulting mixture was extracted with AcOEt (20 mL). The organic extract was washed with $H_2O$ (20 mL) followed by brine (20 mL) and dried over anhydrous $Na_2SO_4$. Filtration and evaporation in vacuo furnished the crude product, which was purified by flash column chromatography ($CHCl_3$ → 50:1 $CHCl_3$/MeOH → 97:3 $CHCl_3$/MeOH) to give **GA-TD-5** (21.9 mg, 69%) as a purple solid. The compound exists as a 1:1 mixture of diastereomers arising from the racemic E3 ligand.

$^1$H NMR (594 MHz, $CDCl_3$) δ 9.41 (s, 1H), 9.142 (s, 0.5H), 9.136 (s, 0.5H), 8.792 (d, *J* = 8.5 Hz, 0.5H), 8.786 (d, *J* = 8.5 Hz, 0.5H), 8.72 (brs, 0.5H), 8.59 (brs, 0.5H), 7.69 (t, *J* = 8.5 Hz, 0.5H), 7.67 (t, *J* = 8.5 Hz, 0.5H), 7.53 (d, *J* = 8.5 Hz, 0.5H), 7.51 (d, *J* = 8.5 Hz, 0.5H), 7.36 (s, 0.5H), 7.32 (s, 0.5H), 7.25 (s, 0.5H), 7.24 (s, 0.5H), 6.94 (d, *J* = 11.7 Hz, 0.5H), 6.92 (d, *J* = 11.7 Hz, 0.5H), 6.57 (t, *J* = 11.7 Hz, 0.5H),

6.55 (t, $J$ = 11.7 Hz, 0.5H), 6.24 (t, $J$ = 6.0 Hz, 1H), 5.88 (m, 1H), 5.86 (m, 1H), 5.24 (s, 0.5H), 5.21 (s, 0.5H), 4.93 (m, 1H), 4.43 (m, 2.5H), 4.31 (d, $J$ = 9.6 Hz, 0.5H), 4.30 (d, $J$ = 9.6 Hz, 0.5H), 4.11 (brs, 0.5H), 3.55 (m, 3H), 3.43 (m, 1H), 3.354 (s, 1.5H), 3.350 (s, 1.5H), 3.27 (s, 1.5H), 3.26 (s, 1.5H), 2.91 (m, 1H), 2.82–2.70 (m, 5H), 2.60 (m, 1H), 2.52–2.44 (m, 2H), 2.25 (m, 2H), 2.23–2.08 (m, 2H), 2.00 (s, 3H), 1.85–1.58 (m, 7H), 1.79 (s, 3H), 1.47 (quint, $J$ = 7.7 Hz, 2H), 0.99 (d, $J$ = 6.9 Hz, 3H), 0.90 (d, $J$ = 6.7 Hz, 1.5H), 0.87 (d, $J$ = 6.7 Hz, 1.5H); $^{13}$C NMR (149 MHz, $CDCl_3$) δ 184.1, 181.1, 181.0, 172.4, 172.3, 171.03, 170.96, 169.3, 169.2, 168.7, 168.1, 168.0, 166.8, 166.7, 156.3, 156.2, 148.53, 148.47, 144.7, 144.6, 141.3, 138.0, 136.6, 136.4, 136.3, 136.2, 134.9, 134.7, 134.0, 133.8, 132.93, 132.86, 131.23, 131.17, 127.5, 127.4, 126.64, 126.57, 125.34, 125.27, 121.2, 121.1, 118.64, 118.60, 115.5, 109.6, 109.5, 109.0, 81.9, 81.5, 81.3, 72.8, 72.7, 57.3, 56.9, 49.44, 49.40, 47.31, 47.27, 42.7, 37.9, 37.8, 35.2, 35.1, 34.35, 34.29, 32.4, 31.5, 30.5, 30.4, 29.1, 28.9, 28.7, 28.6, 25.5, 25.3, 25.0, 24.9, 23.0, 22.9, 22.84, 22.82, 13.0, 12.9, 12.68, 12.67, 12.6, 12.5; HRMS (FAB) $m/z$ $[M + H]^+$ calcd for $C_{52}H_{66}N_9O_{13}$ 1024.4775; found 1024.4766.

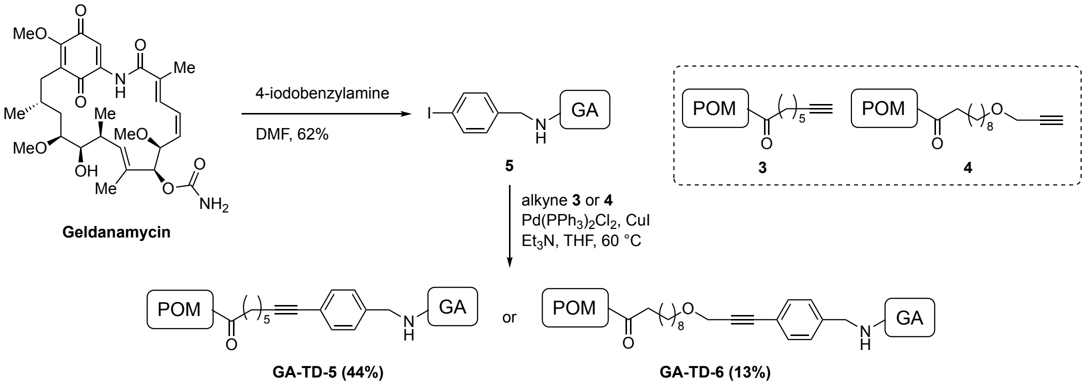


**Scheme 2.**

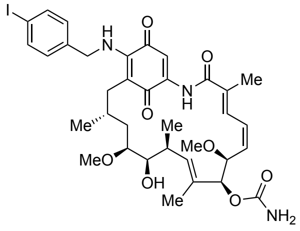


(4*E*,6*Z*,8*S*,9*S*,10*E*,12*S*,13*R*,14*S*,16*R*)-13-Hydroxy-19-((4-iodobenzyl)amino)-8,14-dimethoxy -4,10,12,16-tetramethyl-3,20,22-trioxo-2-azabicyclo[16.3.1]docosa-1(21),4,6,10,18-pentaen-9-yl carbamate (**5**): (4-Iodophenyl)methanamine (55.4 mg, 0.238 mmol) was added to geldanamycin (58.7 mg, 0.105 mmol) in DMF (0.35 mL), and the mixture was stirred for 1 h in the dark. The reaction mixture was diluted with $CH_2Cl_2$ (5 mL). The resulting solution was purified by silica gel column chromatography ($CHCl_3$ → 100:1 $CHCl_3$/MeOH → 50:1 $CHCl_3$/MeOH) to give iodide **5** (112 mg, containing DMF, 62%) as a purple solid. Iodide **5** was used without further purifications.

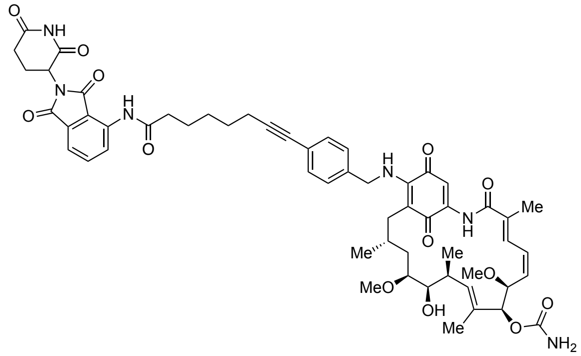


**GA-TD-6**: $Pd(PPh_3)_2Cl_2$ (3.52 mg, 5.01 μmol), $Et_3N$ (20 μL, 150 μmol), and CuI (2.8 mg, 15 μmol) were added to a solution of iodide **5** (53.5 mg, 70.2 μmol) and alkyne **3** (70.0 mg, 177 μmol) in THF (1.0 mL) at room temperature. After 4 h of stirring at 60 °C, the reaction mixture was cooled to room temperature and diluted with $CH_2Cl_2$. The resulting solution was filtered through a Celite pad, and the filtrate was evaporated in vacuo. The crude product was purified by flash column chromatography ($CHCl_3$ → 100:1 $CHCl_3$/MeOH→ 50:1 $CHCl_3$/MeOH → 40:1 $CHCl_3$/MeOH), followed by gel permeation chromatograph to give **GA-TD-6** (22.4 mg, 44%) as a purple solid. The compound exists as a 1:1 mixture of diastereomers arising from the racemic E3 ligand.

$^{1}$H NMR (600 MHz, CDCl$_3$) δ 9.43 (s, 1H), 9.20 (s, 0.5H), 9.17 (s, 0.5H), 8.82 (m, 1H), 8.56 (brs, 0.5H), 8.48 (brs, 0.5H), 7.70 (m 1H), 7.53 (m, 1H), 7.38 (d, *J* = 8.3 Hz, 2H), 7.31 (s, 0.5H), 7.30 (s, 0.5H), 7.17 (d, *J* = 8.3 Hz, 1H), 7.16 (d, *J* = 8.3 Hz, 1H), 6.96 (d, *J* = 11.3 Hz, 1H), 6.58 (t, *J* = 11.3 Hz, 1H), 6.49 (t, *J* = 5.9 Hz, 0.5H), 6.46 (t, *J* = 5.9 Hz, 0.5H), 5.89–5.84 (m, 2H), 5.20 (s, 0.5H), 5.19 (s, 0.5H), 4.95 (m, 1H), 4.88 (brs, 2H), 4.72 (dd, *J* = 14.9, 5.9 Hz, 1H), 4.62 (m, 1H), 4.32 (m, 1H), 4.18 (brs, 0.5H), 4.13 (brs, 0.5H), 3.58 (m, 1H), 3.44 (m, 1H), 3.362 (s, 1.5H), 3.359 (s, 1.5H), 3.28 (s, 3H), 2.90 (m, 1H), 2.83 – 2.71 (m, 3H), 2.62 (m, 2H), 2.50 (m, 2H), 2.45 (t, *J* = 6.9 Hz, 2H), 2.37 (m, 1H), 2.15 (m, 1H), 2.02 (s, 3H), 1.81 (m, 2H), 1.80 (s, 3H), 1.75 (m, 1H), 1.75 (m, 2H), 1.73 (m, 1H), 1.70–1.65 (m, 4H), 1.59 (m, 2H), 1.01 (d, *J* = 7.9 Hz, 3H), 0.99 (d, *J* = 7.0 Hz, 3H); $^{13}$C NMR (151 MHz, CHLOROFORM-*D*) δ 184.01, 183.98, 181.28, 181.25, 172.3, 171.0, 170.9, 169.32, 169.30, 168.6, 168.5, 168.1, 168.0, 166.8, 156.3, 156.2, 144.8, 144.7, 141.4, 141.3, 137.99, 137.98, 136.6, 136.2, 136.14, 136.08, 135.01, 134.95, 133.8, 132.98, 132.96, 132.4, 131.24, 131.23, 127.6, 127.5, 127.3, 127.2, 126.7, 125.40, 125.38, 124.3, 124.2, 118.6, 115.4, 109.4, 109.03, 109.00, 90.99, 90.96, 81.8, 81.6, 81.5, 81.4, 81.3, 80.5, 80.4, 72.81, 72.79, 57.3, 56.8, 49.8, 49.7, 49.4, 38.0, 37.9, 35.1, 34.5, 32.5, 32.4, 31.5, 28.7, 28.41, 28.36, 24.89, 24.86, 23.1, 22.8, 19.4, 12.9, 12.7, 12.5; HRMS (ESI) *m/z* [M + Na]$^+$ calcd for $C_{56}H_{64}N_6O_{13}Na$ 1051.4424; found 1051.4423.

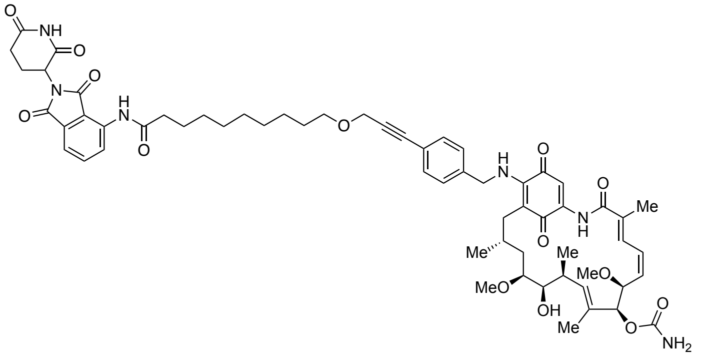


**GA-TD-7**: Pd(PPh$_3$)$_2$Cl$_2$ (3.82 mg, 5.44 μmol), Et$_3$N (20 μL, 150 μmol), and CuI (2.8 mg, 15 μmol)

were added to a solution of iodide **5** (58.0 mg, 76.2 μmol) and alkyne **4** (35.9 mg, 74.6 μmol) in THF (1.0 mL) at room temperature. After 12 h of stirring at 60 °C, the reaction mixture was cooled to room temperature and diluted with $CH_2Cl_2$. The resulting solution was filtered through a Celite pad, and the filtrate was evaporated in vacuo. The crude product was purified by flash column chromatography ($CHCl_3$ → 100:1 $CHCl_3$/MeOH→ 50:1 $CHCl_3$/MeOH → 40:1 $CHCl_3$/MeOH), followed by gel permeation chromatograph to give **GA-TD-6** (10.9 mg, 13%) as a purple solid. The compound exists as a 1:1 mixture of diastereomers arising from the racemic E3 ligand.

$^1$H NMR (594 MHz, $CDCl_3$) δ 9.41 (s, 1H), 9.16 (s, 0.5H), 9.15 (s, 0.5H), 8.83 (d, *J* = 8.5 Hz, 1H), 8.29 (brs, 0.5H), 8.27 (brs, 0.5H), 7.71 (dd, *J* = 8.5, 7.3 Hz, 1H), 7.54 (d, *J* = 7.3 Hz, 1H), 7.47 (d, *J* = 8.2 Hz, 2H), 7.30 (s, 1H), 7.22 (d, *J* = 8.2 Hz, 2H), 6.96 (d, *J* = 11.6 Hz, 1H), 6.58 (t, *J* = 11.6 Hz, 1H), 6.46 (m, 1H), 5.89 (dd, *J* = 5.9, 5.3 Hz, 1H), 5.87 (m, 1H), 5.20 (s, 1H), 4.95 (m, 1H), 4.74 (dd, *J* = 15.2, 5.9 Hz, 1H), 4.64 (dd, *J* = 15.2, 5.3 Hz, 1H), 4.36 (s, 2H), 4.32 (d, *J* = 9.8 Hz, 3H), 4.10 (brs, 1H), 3.58 (m, 3H), 3.44 (m, 1H), 3.36 (s, 3H), 3.28 (s, 3H), 2.92 (m, 1H), 2.84–2.71 (m, 3H), 2.63 (m, 1H), 2.44 (m, 2H), 2.37 (m, 1H), 2.16 (m, 1H), 2.02 (s, 3H), 1.80 (s, 3H), 1.78 – 1.69 (m, 5H), 1.63 (quint, *J* = 6.7 Hz, 2H), 1.45–1.28 (m, 10H), 1.02–0.98 (m, 6H); $^{13}$C NMR (149 MHz, $CDCl_3$) δ 184.0, 181.4, 172.58, 172.56, 170.8, 169.3, 168.6, 167.94, 167.92, 166.8, 156.2, 144.8, 141.3, 138.1, 137.1, 136.6, 136.1, 135.0, 133.9, 133.0, 132.7, 131.2, 127.7, 127.2, 126.7, 125.4, 123.1, 118.5, 115.4, 109.4, 109.0, 86.6, 85.4, 81.9, 81.6, 81.4, 72.8, 70.5, 58.9, 57.3, 56.9, 49.8, 49.4, 38.2, 35.2, 34.5, 32.5, 31.5, 29.8, 29.7, 29.5, 29.4, 29.3, 28.7, 26.2, 25.4, 23.1, 22.8, 12.9, 12.7, 12.5; HRMS (ESI) *m*/*z* $[M + Na]^+$ calcd for $C_{61}H_{74}N_6O_{14}Na$ 1137.5155; found 1137.5138.

**Copies of $^1$H and $^{13}$C NMR spectra**

| parameters | |
|---|---|
| Comment | 17-AzPAG 1H |
| Instrument | ECA 600SL |
| Solvent | CHLOROFORM-D |
| Temperature | 25.4 |
| Number of Scans | 16 |
| Spectrometer Frequency | 594.17 |
| Spectral Width | 8912.7 |
| Lowest Frequency | -1460.0 |
| Nucleus | 1H |
| Digital Resolution | 0.08 |

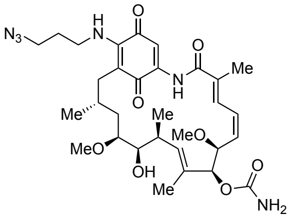


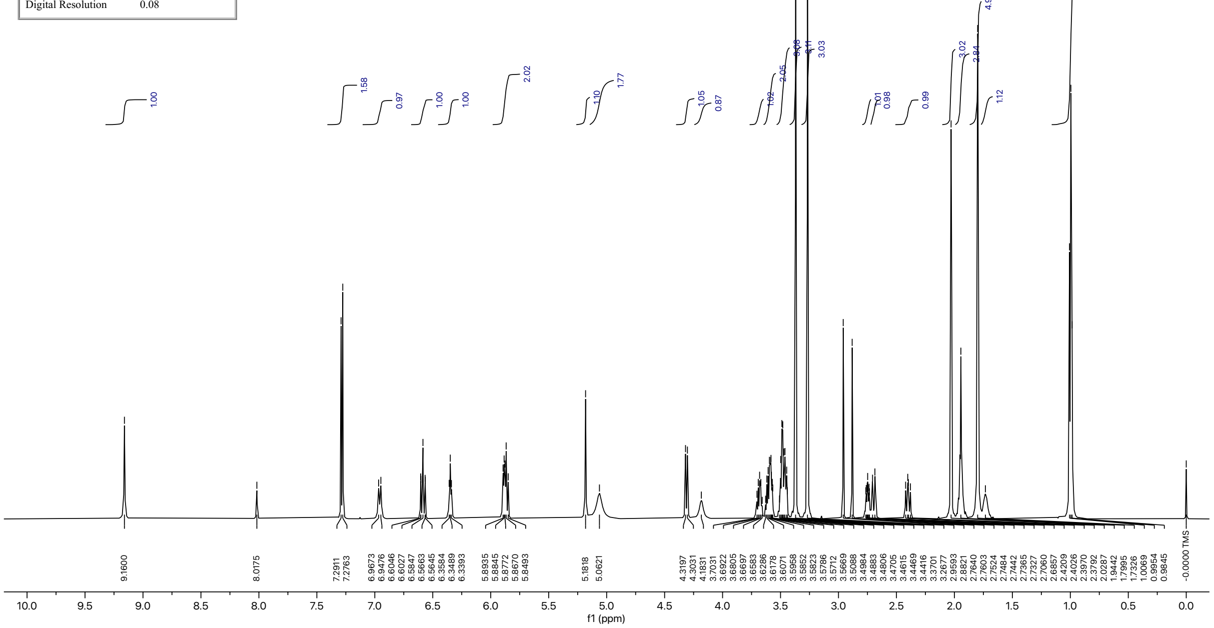

| parameters | |
| --- | --- |
| Comment | 17-AzPAG 13C |
| Instrument | ECA 600SL |
| Solvent | CHLOROFORM-D |
| Temperature | 26.5 |
| Number of Scans | 4000 |
| Spectrometer Frequency | 149.40 |
| Spectral Width | 37594.0 |
| Lowest Frequency | -3856.3 |
| Nucleus | 13C |
| Digital Resolution | 0.72 |

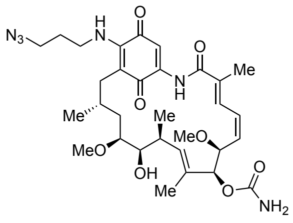


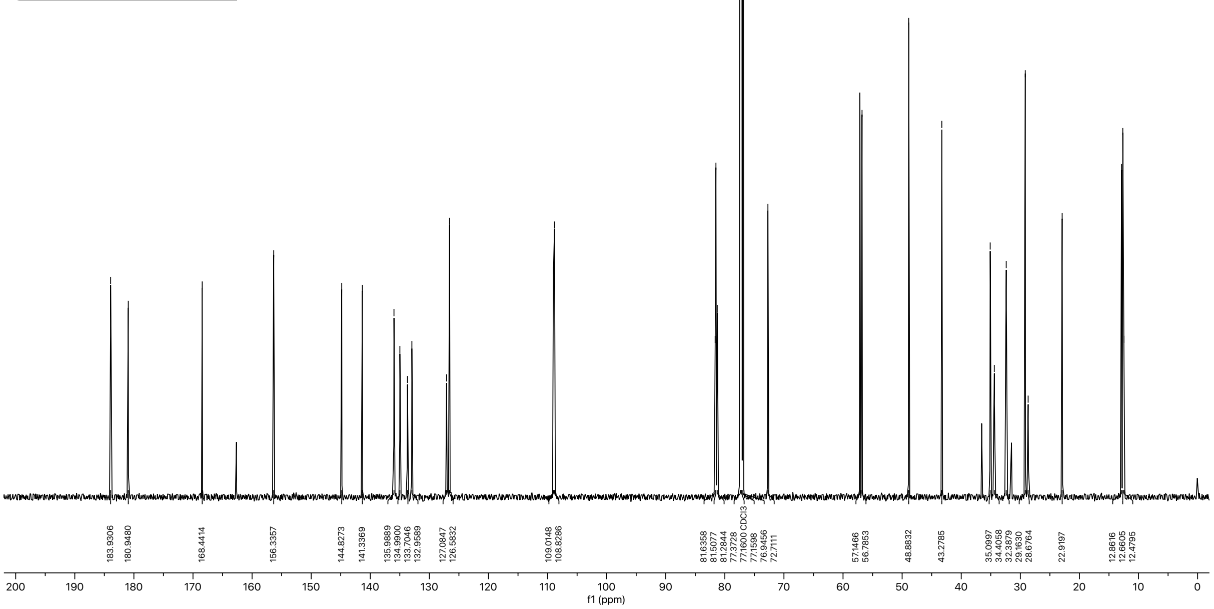

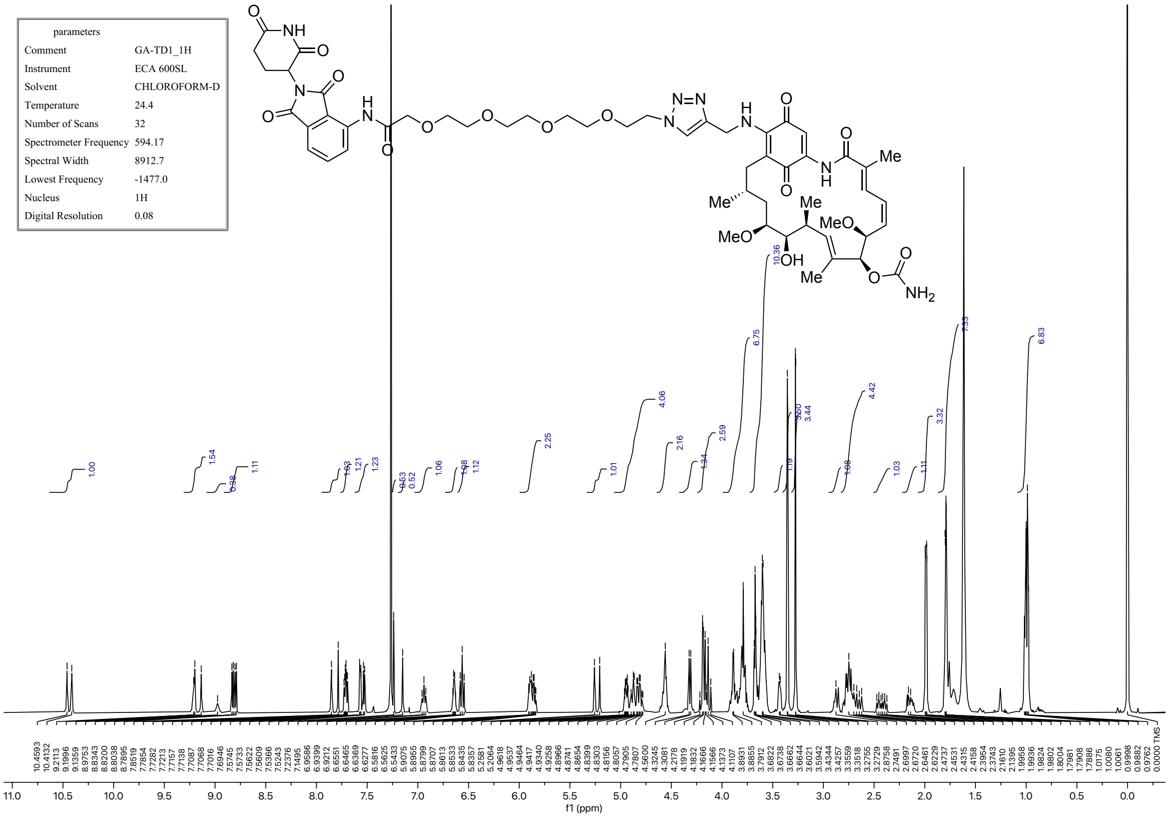
parameters
Comment GA-TD1_1H
Instrument ECA 600SL
Solvent CHLOROFORM-D
Temperature 24.4
Number of Scans 32
Spectrometer Frequency 594.17
Spectral Width 8912.7
Lowest Frequency -1477.0
Nucleus 1H
Digital Resolution 0.08
f1 (ppm)

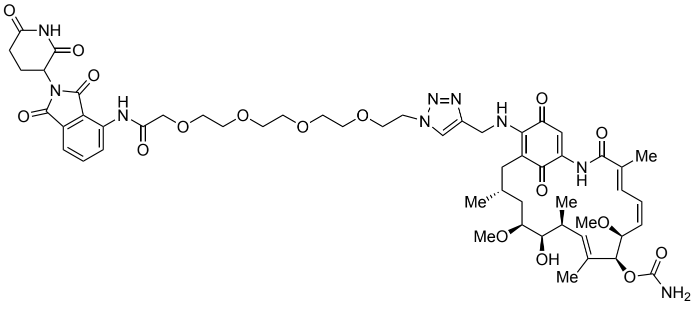

| parameters | |
| --- | --- |
| Comment | GA-TD1_13C |
| Instrument | ECA 600SL |
| Solvent | CHLOROFORM-D |
| Temperature | 25.8 |
| Number of Scans | 50000 |
| Spectrometer Frequency | 149.40 |
| Spectral Width | 37594.0 |
| Lowest Frequency | -3846.8 |
| Nucleus | 13C |
| Digital Resolution | 0.72 |

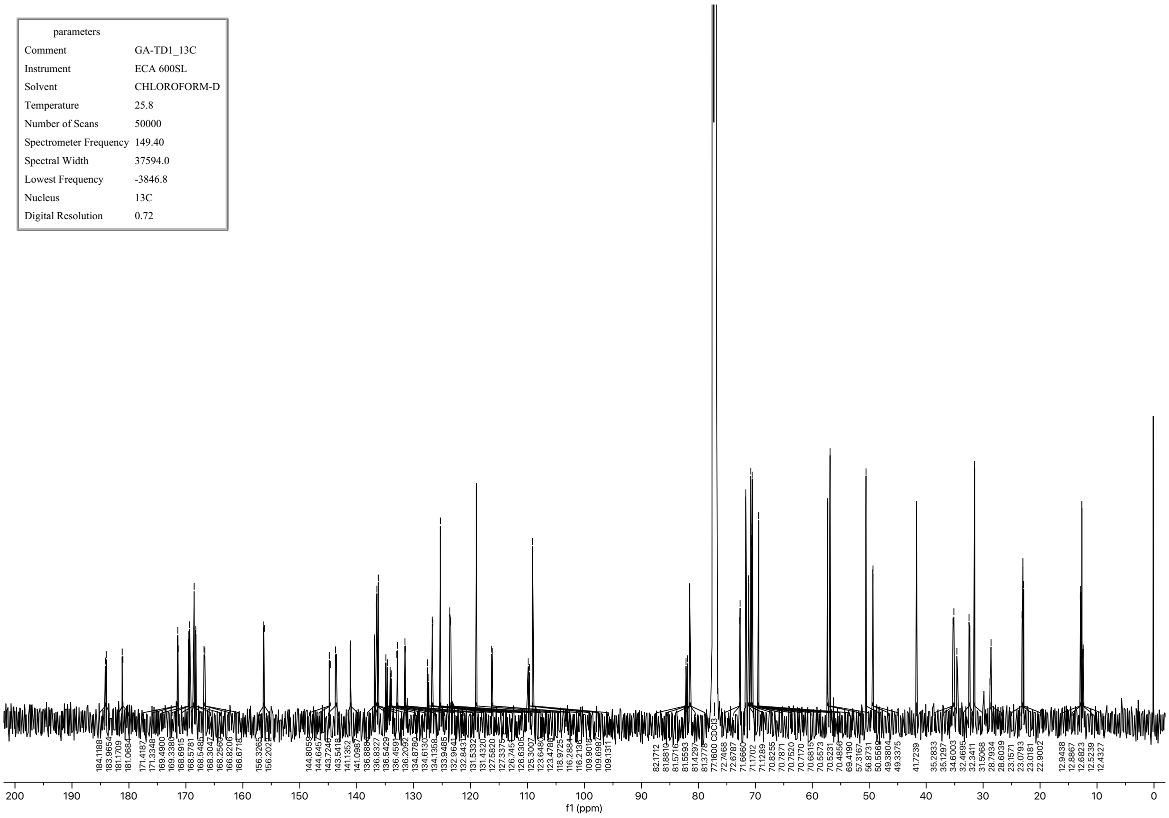

| parameters | |
|---|---|
| Comment | GA-TD3_1H |
| Instrument | JNM-ECZ600R/ S1 |
| Solvent | CHLOROFORM-D |
| Temperature | 23.3 |
| Number of Scans | 16 |
| Spectrometer Frequency | 600.17 |
| Spectral Width | 9025.3 |
| Lowest Frequency | -1502.1 |
| Nucleus | 1H |
| Digital Resolution | 0.09 |

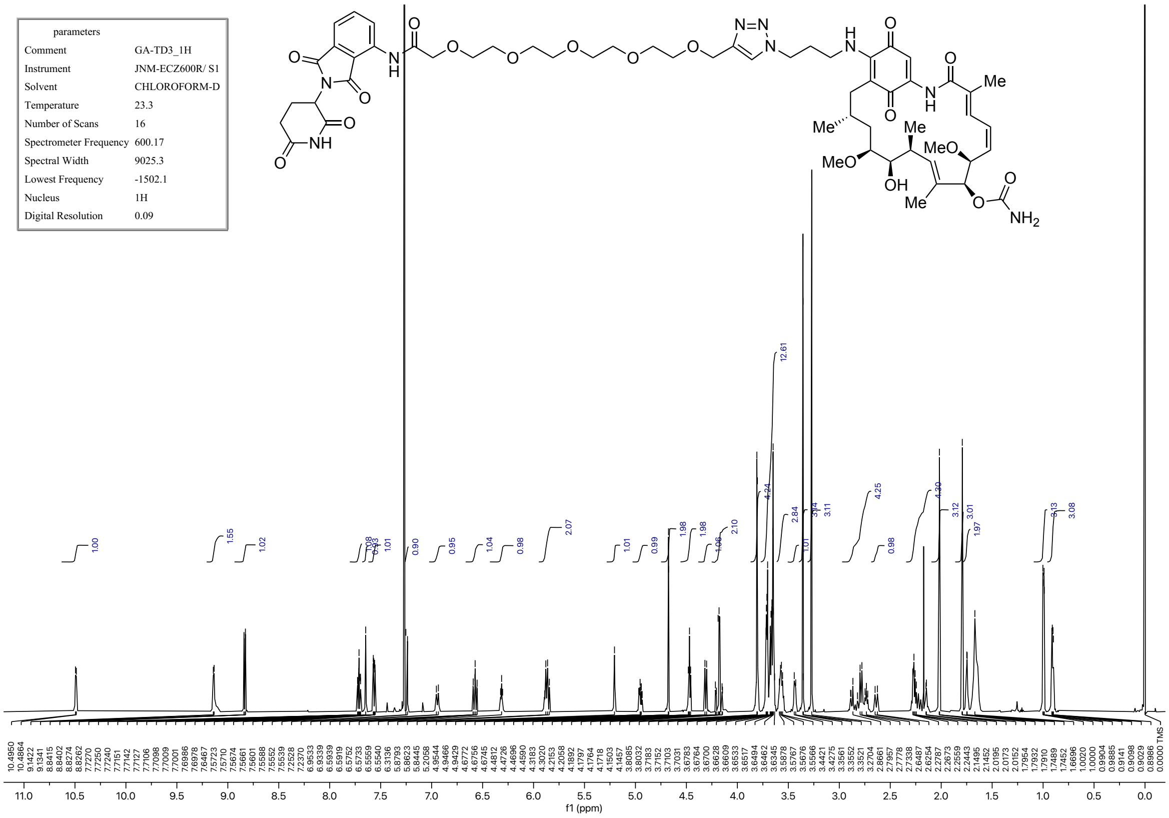

| parameters | |
|---|---|
| Comment | GA-TD3_13C |
| Instrument | JNM-ECZ600R/ S1 |
| Solvent | CHLOROFORM-D |
| Temperature | 22.3 |
| Number of Scans | 18040 |
| Spectrometer Frequency | 150.91 |
| Spectral Width | 37878.8 |
| Lowest Frequency | -3839.4 |
| Nucleus | 13C |
| Digital Resolution | 0.36 |

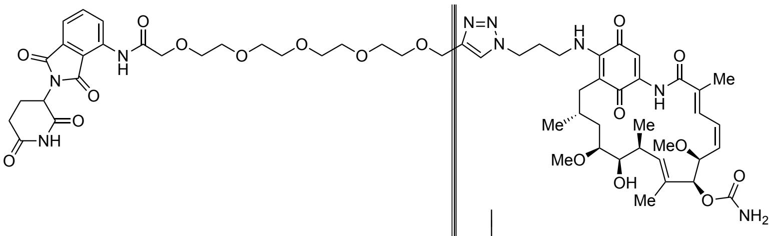


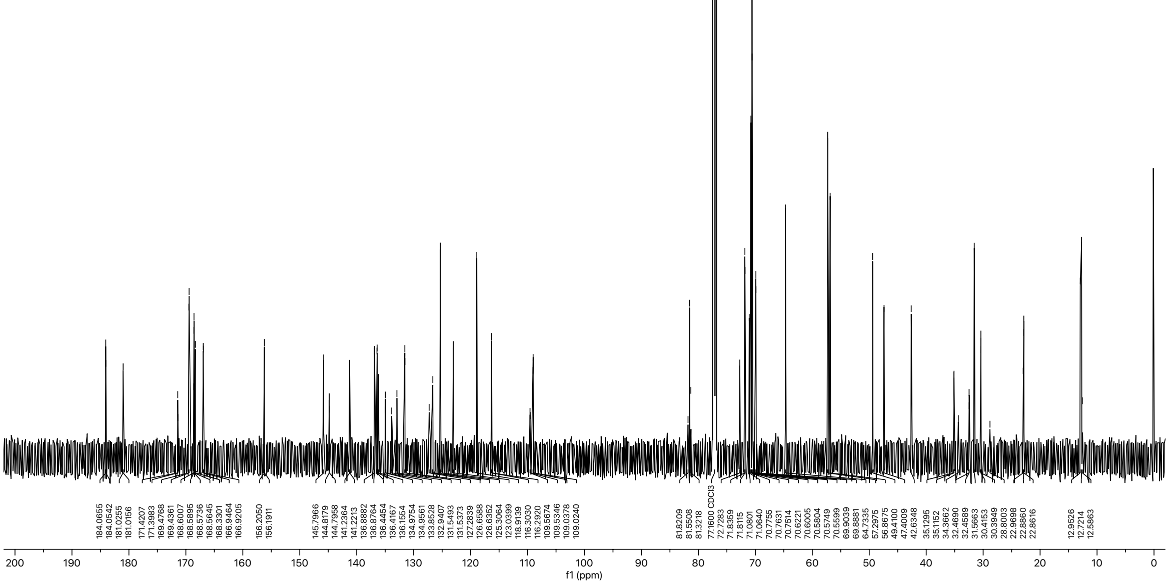

| parameters | |
|---|---|
| Comment | GA-TD5_1H |
| Instrument | ECA 600SL |
| Solvent | CHLOROFORM-D |
| Temperature | 24.5 |
| Number of Scans | 16 |
| Spectrometer Frequency | 594.17 |
| Spectral Width | 8912.7 |
| Lowest Frequency | -1476.5 |
| Nucleus | 1H |
| Digital Resolution | 0.08 |

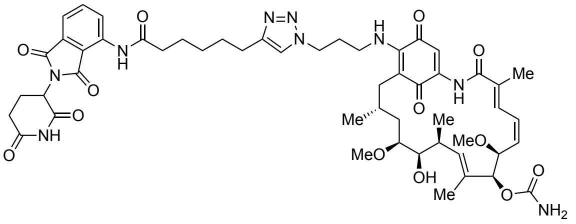


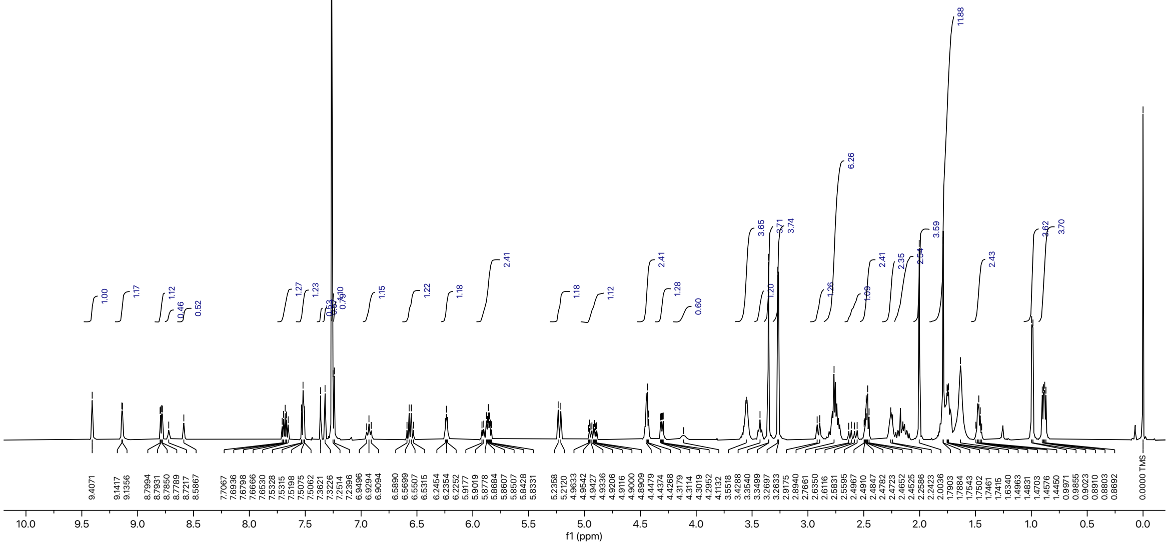

| parameters | |
|---|---|
| Comment | GA-TD5-13C |
| Instrument | ECA 600SL |
| Solvent | CHLOROFORM-D |
| Temperature | 25.4 |
| Number of Scans | 20601 |
| Spectrometer Frequency | 149.40 |
| Spectral Width | 37594.0 |
| Lowest Frequency | -3847.7 |
| Nucleus | 13C |
| Digital Resolution | 0.72 |

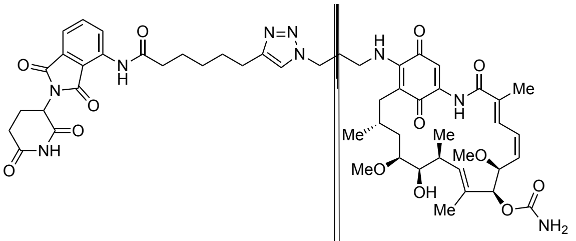


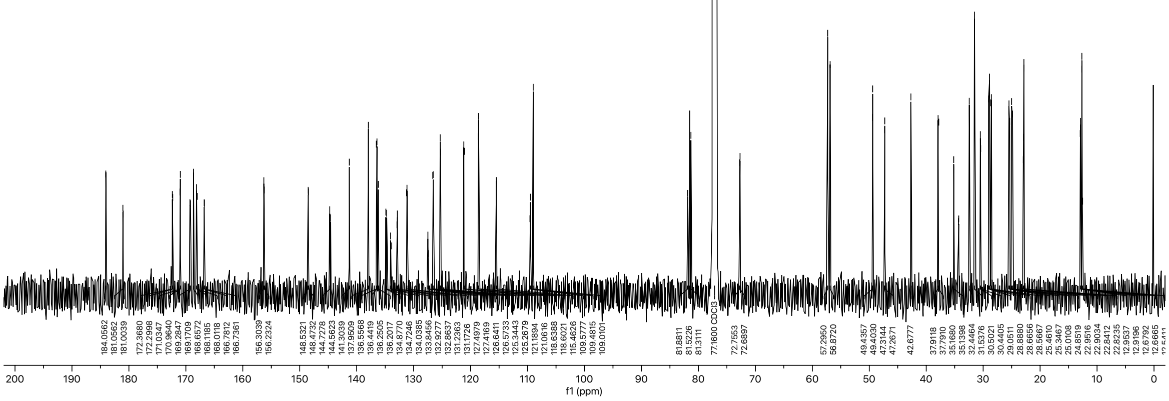

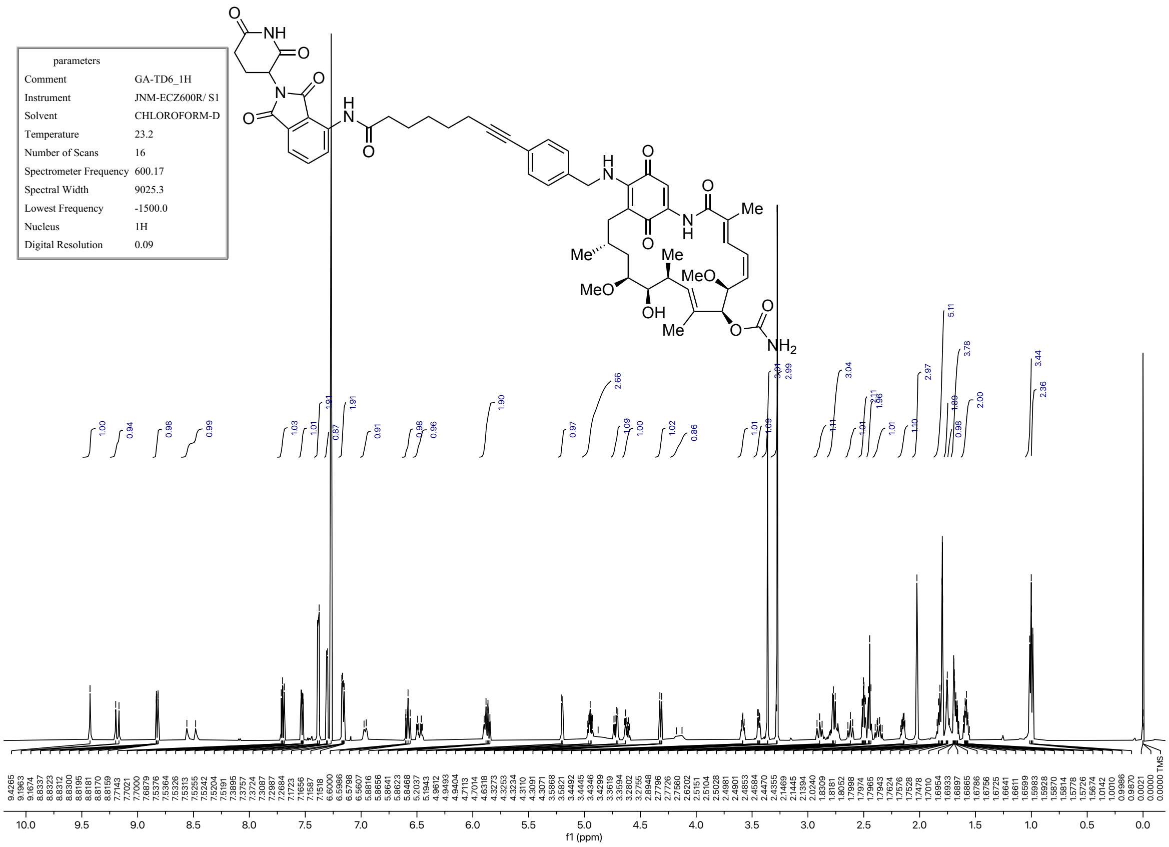


| parameters | |
|---|---|
| Comment | GA-TD6_1H |
| Instrument | JNM-ECZ600R/ S1 |
| Solvent | CHLOROFORM-D |
| Temperature | 23.2 |
| Number of Scans | 16 |
| Spectrometer Frequency | 600.17 |
| Spectral Width | 9025.3 |
| Lowest Frequency | -1500.0 |
| Nucleus | 1H |
| Digital Resolution | 0.09 |

| parameters | |
|---|---|
| Comment | GA-TD6_13C |
| Instrument | JNM-ECZ600R/ S1 |
| Solvent | CHLOROFORM-D |
| Temperature | 23.4 |
| Number of Scans | 2048 |
| Spectrometer Frequency | 150.91 |
| Spectral Width | 37878.6 |
| Lowest Frequency | -3840.9 |
| Nucleus | 13C |
| Digital Resolution | 0.36 |

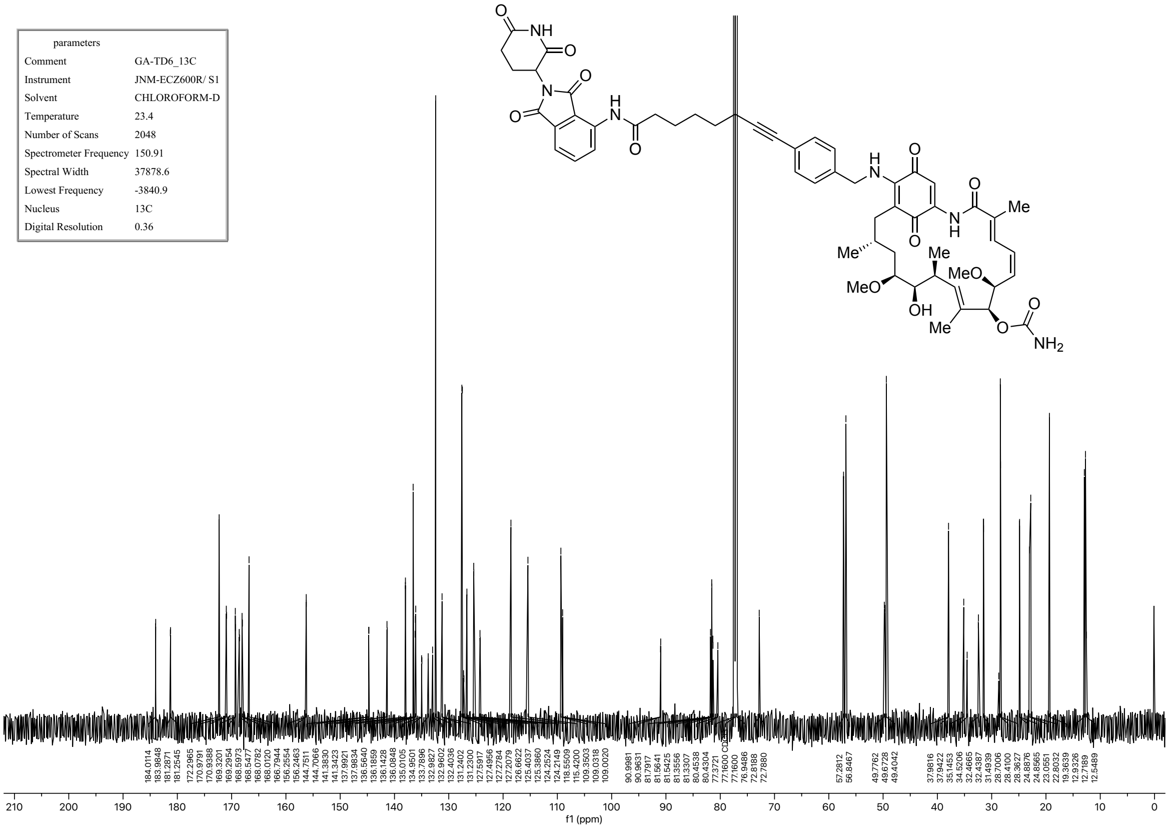

| parameters | |
|---|---|
| Comment | GA-TD7_1H |
| Instrument | ECA 600SL |
| Solvent | CHLOROFORM-D |
| Temperature | 24.4 |
| Number of Scans | 64 |
| Spectrometer Frequency | 594.17 |
| Spectral Width | 8912.7 |
| Lowest Frequency | -1478.2 |
| Nucleus | 1H |
| Digital Resolution | 0.08 |

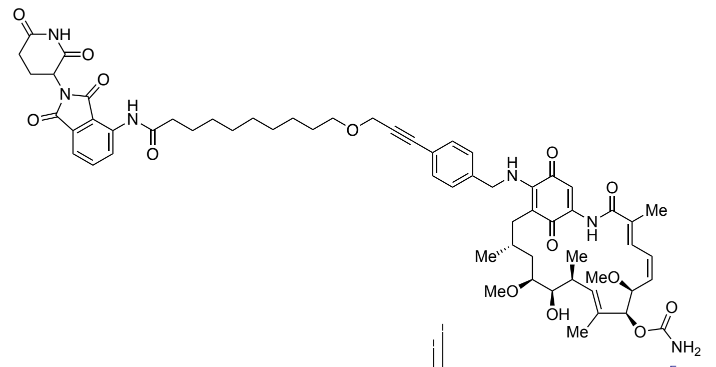

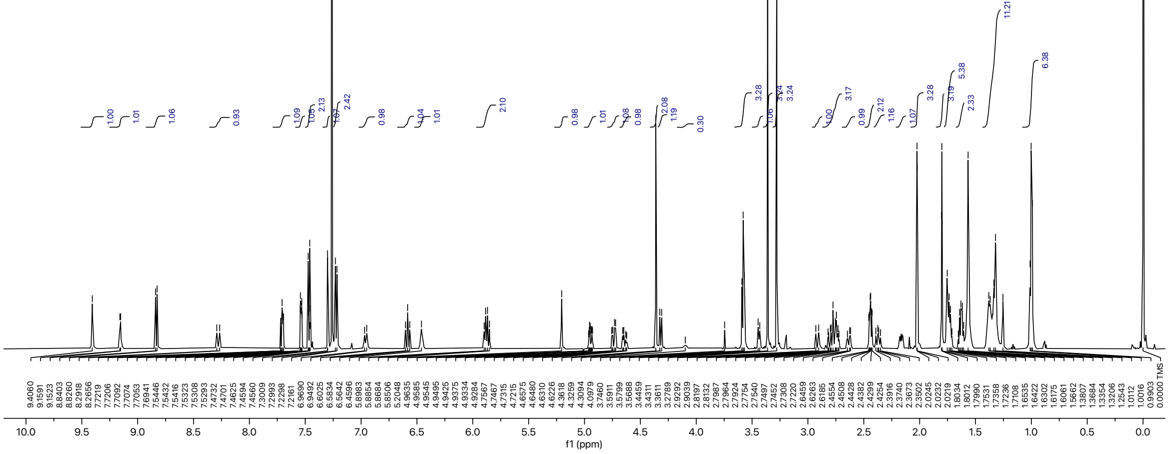

| parameters | |
|---|---|
| Comment | GA-TD7_13C |
| Instrument | ECA 600SL |
| Solvent | CHLOROFORM-D |
| Temperature | 25.2 |
| Number of Scans | 37301 |
| Spectrometer Frequency | 149.40 |
| Spectral Width | 37594.0 |
| Lowest Frequency | -3846.5 |
| Nucleus | 13C |
| Digital Resolution | 0.72 |

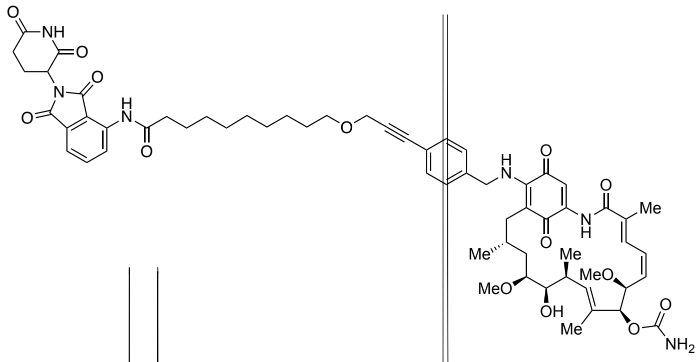


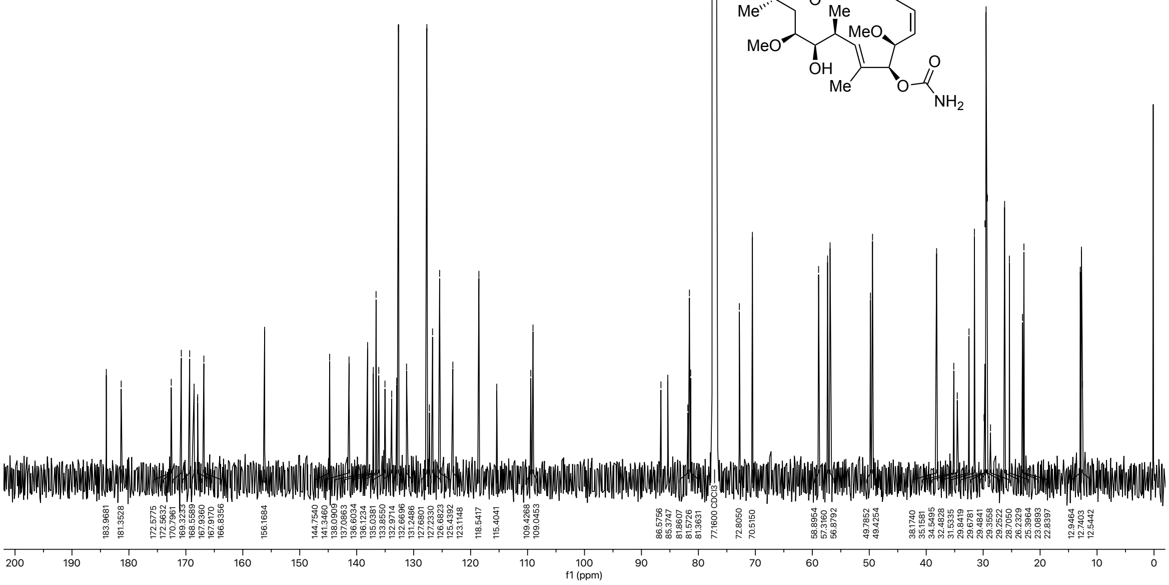
